# Two FtsZ proteins orchestrate archaeal cell division through distinct functions in ring assembly and constriction

**DOI:** 10.1101/2020.06.04.133736

**Authors:** Yan Liao, Solenne Ithurbide, Christian Evenhuis, Jan Löwe, Iain G. Duggin

**Affiliations:** The ithree institute, University of Technology Sydney, Ultimo, NSW, 2007, Australia; MRC Laboratory of Molecular Biology, Francis Crick Avenue, Cambridge, CB2 0QH, UK

**Keywords:** FtsZ, tubulin, archaea, cell division, cytokinesis, cell envelope, cell wall, S-layer, cell morphology

## Abstract

The tubulin homolog FtsZ assembles a cytokinetic ring in bacteria and plays a key role in the machinery that constricts to divide the cells. Many archaea encode two FtsZ proteins from distinct families, FtsZ1 and FtsZ2, of previously unclear functions. Here we show that *Haloferax volcanii* cannot divide properly without either or both, but DNA replication continues, and cells proliferate in alternative ways via remarkable envelope plasticity. FtsZ1 and FtsZ2 co-localize to form the dynamic division ring. However, FtsZ1 can assemble rings independently of FtsZ2, and stabilizes FtsZ2 in the ring, whereas FtsZ2 functions primarily in the constriction mechanism. FtsZ1 also influenced cell shape suggesting it forms a hub-like platform at midcell for the assembly of shape-related systems too. Both FtsZ1 and FtsZ2 are widespread in archaea with a single S-layer envelope, but archaea with a pseudomurein wall and division septum only have FtsZ1. FtsZ1 is therefore likely to provide a fundamental recruitment role in diverse archaea, and FtsZ2 is required for constriction of a flexible S-layer envelope, where an internal constriction force might dominate the division mechanism, in contrast to the single-FtsZ bacteria and archaea that divide primarily by wall ingrowth.

---

FtsZ is the most conserved bacterial cell division protein and among the first molecules to assemble at the incipient division site. It forms a dynamic ring-like structure around the cell that acts as a platform to initiate recruitment of about a dozen core components and up to ∼30 other factors involved in division^1,2^. Within the ring, FtsZ polymerizes dynamically in a GTP-dependent manner near the inner surface of the cytoplasmic membrane, directing the inward synthesis of septal cell wall peptidoglycan to drive cell constriction, and may also provide a direct influence on membrane constriction^3-7^. The septal wall splits to separate the two new cells in a process that is either continuous with septal wall synthesis and ring closure or is activated after the septal cross-wall is complete^8,9^.

An *ftsZ* homolog was first identified in several archaea by DNA hybridization^10,11^ and GTP-binding screens^12^. Archaeal FtsZ showed GTPase activity *in vitro*^11^ and was localized to mid-cell by immunofluorescence^11,13^. In bacteria such as *E. coli*, the actin homolog FtsA forms filaments that, together with ZipA and several regulatory or stabilization proteins, link FtsZ filaments to the inner membrane^2,14^. However, homologs of these and most other bacterial (and eukaryotic) cell division proteins appear to be largely absent in archaea. One exception and likely candidate is the archaeal homolog of SepF, which can polymerize and participates with FtsA in anchoring FtsZ for Gram-positive bacterial division^15^, and shows strong phylogenetic association with FtsZ in archaea^16^.

The first complete archaeal genome sequence, of *Methanocaldococcus jannaschii*,^17^ revealed multiple *ftsZ* homologs, whereas almost all bacterial genomes contain only one *ftsZ*. Some archaea do not contain FtsZ, including the phylum Crenarchaeota, where division is instead orchestrated by proteins related to the ESCRT-III complexes involved in membrane remodelling in eukaryotes^18,19^. Many of the archaea that contain FtsZ also have one or more tubulin-superfamily proteins from the CetZ family^20,21^. The genome of the archaeal model organism *Haloferax volcanii*^22^ encodes eight tubulin-superfamily proteins—six CetZs, and two FtsZs. Work in *H. volcanii* showed that the CetZs were not individually required for division, but at least one is required for rod cell shape development^20^. In the same study, an FtsZ1-GFP fusion protein was observed to localize at midcell. The results also suggested *H. volcanii* as an excellent model for archaeal cell biology, as it has relatively large and flat cells that show a variety of regulated morphologies and are well-suited to light and electron microscopy^20,23,24^. Here we use *H. volcanii* to demonstrate that FtsZ1 and FtsZ2 have different functions in the mechanism of archaeal cell division.

## Results

### Characteristics of two FtsZ families in archaea

We performed a molecular phylogenetic analysis of 149 tubulin-superfamily proteins identified in 60 diverse archaeal genomes. Archaeal FtsZ1, FtsZ2, and the well-studied bacterial/plant FtsZ form three distinct families that are approximately equally divergent from one another (Fig. S1a). A majority of the archaea encoded both an FtsZ1 and an FtsZ2 homolog, including most Euryarchaeota (*e*.*g*., Halobacteria, Thermococci and Archaeoglobi) and the DPANN and Asgard superphyla (Table S3). The Crenarchaeota and Thaumarchaeota lacked specific members of the FtsZ families, whereas methanogenic archaea, that characteristically have a pseudomurein wall not found in other species, were found to only possess FtsZ1 (Fig. S1, Table S3).

We noticed conserved differences between archaeal FtsZ1 and FtsZ2 that are primarily in the vicinity of the GTPase active site and longitudinal subunit interface in polymers (see Supplementary Results and Discussion, Fig. S1c-e). Conserved motifs were also identified in the N- and C-terminal tails that differed from bacterial/plant FtsZ. These differences are likely to manifest as fundamental functional characteristics of the three FtsZ families but await further detailed investigations.

### Depletion of FtsZ1 or FtsZ2 causes division defects in H. volcanii

Given that *ftsZ* genes are nearly ubiquitous and important for survival in bacteria^25^, we first engineered *H. volcanii* strains in which expression of the genomic copy of the *ftsZ1* (HVO_0717) or *ftsZ2* (HVO_0581) gene was placed under the control of the highly specific tryptophan-regulated promoter *p*.*tna*^26^ (Fig. S2); expression of each *ftsZ* could be maintained during strain construction and growth by the inclusion of tryptophan (Trp) in the growth medium, or depleted by placing the cells into medium without Trp. The two corresponding strains grew well in the presence of 2 mM Trp and appeared of approximately normal size and shape (Fig. 1a, 0 h, leftmost panels). However, resuspension of growing cells in liquid medium without Trp caused a large increase in cell size over the 24 h sampling period (biomass doubling times of ∼4.4 h over the first 9 h of log growth), as shown by phase-contrast microscopy (Fig. 1a) and Coulter cytometry cell-volume distributions (Fig. S3). This is morphologically analogous to the classical bacterial filamentous phenotype^27^, and signifies a defect in the regulation or mechanism of cell division in both the *ftsZ1*- and *ftsZ2*-depletion strains. The *ftsZ2* depletion resulted in a stronger division defect (significantly larger cells) compared to the *ftsZ1* depletion (Fig. 1a, S3). Western blotting showed that the cellular amount of FtsZ1 had decreased substantially by ∼6-9 h after withdrawal of Trp from the *p*.*tna-ftsZ1* strain, whereas in the *p*.*tna-ftsZ2* strain, FtsZ2 had almost completely disappeared after 3 h (Fig. 1b). Whether the cell size difference is due to the different rates of depletion or differing roles and importance of the two FtsZ proteins to cell division is addressed further below.

**Figure 1.**
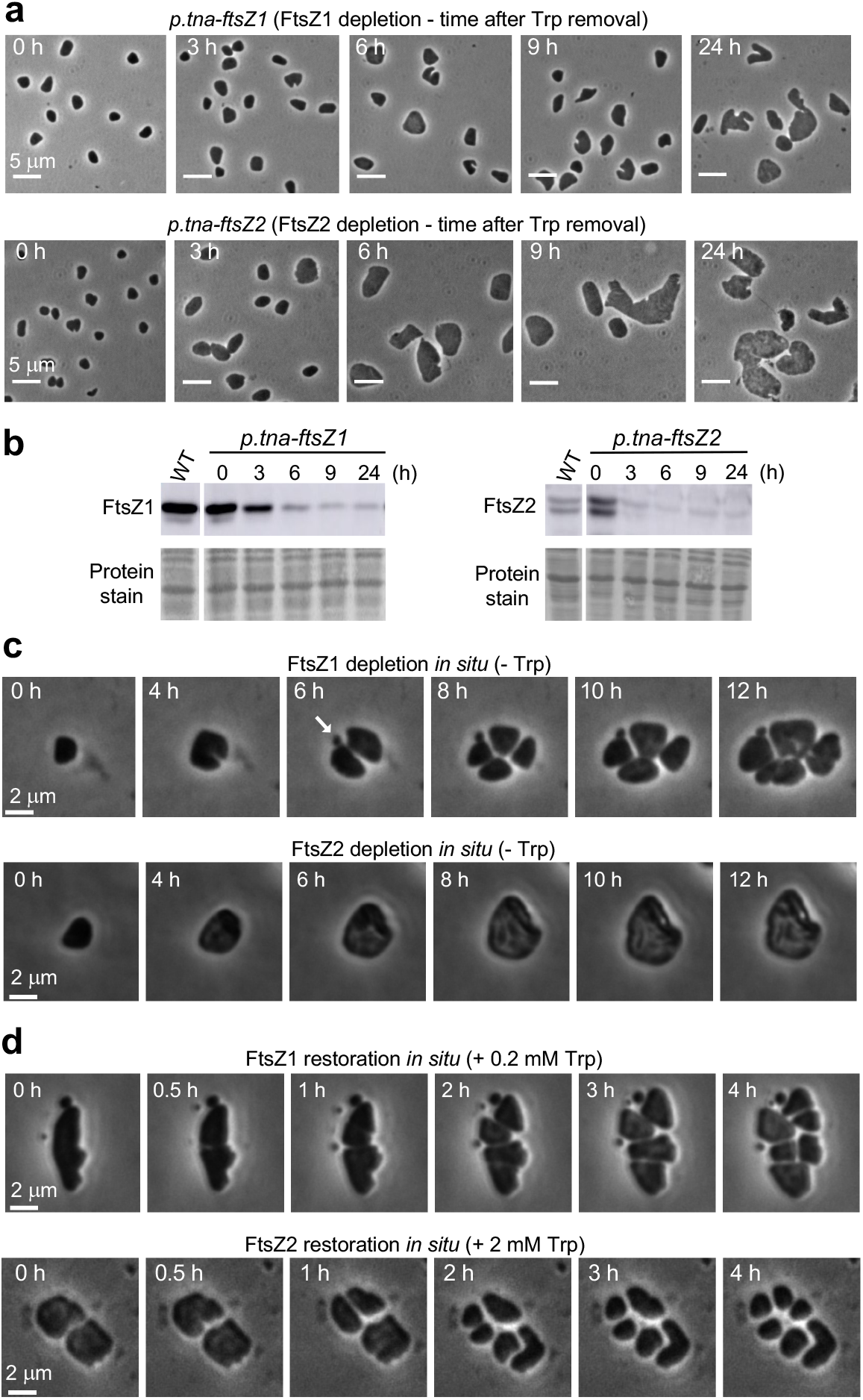
Depletion of FtsZ1 or FtsZ2 results in defects in cell division. (a) Phase-contrast microscopy of samples from cultures (*H. volcanii* ID56 - *p*.*tna-ftsZ1*, and ID57 - *p*.*tna-ftsZ2*) immediately prior to (0 h) or at the indicated time-points after resuspension of cells (to OD_600_ = 0.25) in growth medium without Trp. (b) Western blots probed with FtsZ1 or FtsZ2 antisera during depletion with the corresponding Ponceau S pre-staining of total protein. The separate wild-type (H26) panels were from the same blots for each protein; FtsZ1 (∼40 kDa) and FtsZ2 (∼43 kDa) bands are indicated. The double band for FtsZ2 suggests a modified/clipped form. (c) *In situ* time-lapse imaging of FtsZ depletion. Cells cultured in Hv-Cab + 2 mM Trp were washed and placed onto agarose media pads without Trp and imaged over time. Morphology of giant cells differs from the liquid culture (a), most likely due to the support provided by growth on agarose. (d) *In situ* time-lapse imaging of FtsZ restoration. The respective *p*.*tna-ftsZ* strains were initially grown for ∼2 days without Trp inducer, and then cells were washed and placed onto agarose media (Hv-Cab) pads including 0.2 mM Trp for FtsZ1, and 2 mM Trp for FtsZ2. Scale bars (c-d), 2 µm.

Time-lapse microscopy of growing cells showed some FtsZ1-depleted cells dividing, apparently inefficiently, whereas FtsZ2 depletion appeared to strongly block division (*e*.*g*., Fig. 1c). After the onset of depletion, some *p*.*tna-ftsZ2* cells initially developed a constriction but were unable to complete division; the partial constriction then reversed with further cell growth yet had a lingering influence on cell shape (Fig. S4). Later in depletion, budding-like processes were observed that resulted in apparent vesicles or minicells of variable sizes (Video S1, *p*.*tna-ftsZ2*). In some of the enlarged FtsZ1-depleted cells, a small cell fragment appeared at one side of the division plane during cell separation (Fig. 1c, arrow) or in others appeared to be ‘carved out’ from a cell edge (Video S2). In contrast, FtsZ2-depleted cells were not seen actively dividing, although some showed blebbing from cellular lobes in these very large misshapen cells (Video S3).

FtsZ-depleted cells that were re-supplied with Trp at the commencement of time-lapse imaging (FtsZ restoration) showed that the giant cells then became capable of dividing (Fig. 1d, Video S1). Some underwent multiple simultaneous divisions, generating large cells that then underwent further division events to eventually re-establish a population of normal-sized cells. It was also clear that the division process itself was sometimes asymmetric (unilateral), where cell constriction occurs from only one edge (or predominately from one edge), and the plane of division was sometimes noticeably acentral or not straight nor perpendicular to the cell envelope (Video S1). Irregular positioning of division sites may be expected based on the sensitivity of division site positioning mechanisms to cell size and shape^28^, and this resulted in noticeable pleomorphology in the progeny.

### DNA composition of H. volcanii cells depleted of FtsZ1 or FtsZ2

To assess the positioning and replication of DNA in cells after depletion of each FtsZ, we stained cells with SYTOX Green (SG) DNA stain. The giant cells showed DNA-SG staining that appeared diffusely present throughout the cytoplasm, similar to the staining intensity and distribution in wild-type cells (Fig. S5a). Flow cytometry showed that DNA-SG fluorescence correlated with the side-scatter signal as a proxy for cell size (Fig. S5b). Some of the largest mutant cells (*e*.*g*., after *ftsZ2*-depletion) showed ∼100-fold higher DNA-SG fluorescence and side-scatter than wild-type, corresponding to an expected genome copy number of ∼2000 or more in the largest mutant cells, compared to the typical polyploidy of ∼20 in unmodified *H. volcanii*^29^. These results indicate that DNA synthesis continues in approximate proportion to cell biomass increase during inhibition of cell division caused by FtsZ depletion, and are consistent with the previously observed simple correlation between DNA content and cell size in wild-type *H. volcanii*^30^ that reflects an absence of distinct cell-cycle phases (Fig. S5b).

### H. volcanii propagates without FtsZ1 and/or FtsZ2

The *p*.*tna-ftsZ1* and *p*.*tna-ftsZ2* strains were grown continuously at a range of Trp concentrations (from 0 to 2 mM Trp) and showed an inverse relationship between Trp concentration and cell size (Fig. S6a-b). Since both *ftsZ*-depletion strains could be maintained indefinitely in standard growth medium (Hv-Cab) in the absence of added Trp, we sought to determine whether the *ftsZ* genes were completely dispensable by employing a direct selection procedure to delete each gene. The Δ*ftsZ1* and Δ*ftsZ2* strains were viable and we also made a double deletion (Δ*ftsZ1* Δ*ftsZ2*) and then confirmed the deletions by PCR and genome sequencing. Mid-log cultures had biomass doubling times of 3.08±0.06 h (H26 wild type), 3.58±0.27 h (Δ*ftsZ1*), 3.56±0.23 h (Δ*ftsZ2*) and 4.20±0.18 h (Δ*ftsZ1* Δ*ftsZ2*) (mean±95% CI, Fig. S7a). Dilution and agar plating of mid-log samples (OD_600_ = 0.2) showed that the three Δ*ftsZ* strains had 20-25-fold fewer colony forming units (cfu) compared to the wild-type (Fig. 2a), consistent with a larger cell size and/or a reduced fraction of viable cells. The Δ*ftsZ* strains displayed very heterogeneously sized and misshapen cells, and a significant quantity of cellular debris (Fig. 2b-c), very similar to the respective *p*.*tna-ftsZ* strains grown continuously without Trp (Fig. S6a). The Δ*ftsZ2* and Δ*ftsZ1* Δ*ftsZ2* strains showed more severe cell-division defects compared to Δ*ftsZ1*, indicating that FtsZ2 has a limited capacity to support division in the absence of FtsZ1 (Fig. 2b-c). The equally low cfu count of Δ*ftsZ1* therefore suggests FtsZ1 may have additional functions that contribute to cell viability. Furthermore, the relatively moderate FtsZ1-depletion cell-division phenotype (Fig. 1a) appears to be largely caused by differing roles of FtsZ1 and FtsZ2 rather than the differences in depletion rate (Fig. 1b).

**Figure 2.**
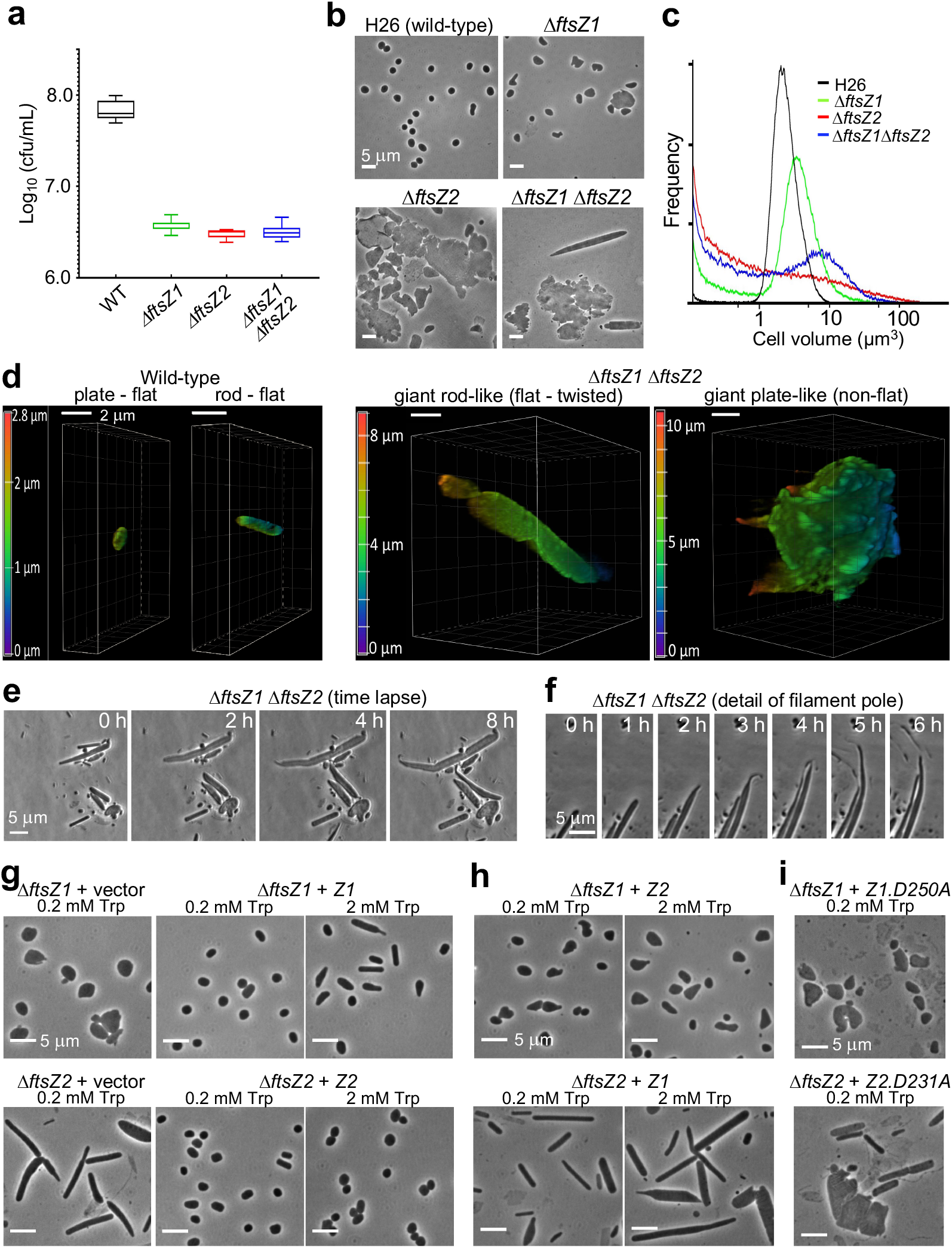
FtsZ1 and FtsZ2 are dispensable for survival but required for normal cell division. (a) Box-and-whisker plots of agar colony counts (cfu/mL) of *H. volcanii* wild-type (H26), Δ*ftsZ1* (ID76), Δ*ftsZ2* (ID77), and Δ*ftsZ1* Δ*ftsZ2* (ID112), sampled during mid-log growth (OD_600_ = 0.2) in Hv-Cab medium (+ 50 µg/mL uracil) (n = 4 separate cultures). (b) Phase-contrast images and (c) Coulter cell-volume frequency distributions (normalized to total count). (d) Confocal microscopy 3D-reconstructed images of cells stained with Mitotracker Orange and embedded in low melting point agarose. The reconstructions are shown with a ∼45° rotation around the x-axis. The colour scale indicates the z-depth. Similar morphologies were seen with other *ftsZ* mutants and FM1-43 membrane staining (Fig. S7b). (e) Live cell time-lapse images of the Δ*ftsZ1* Δ*ftsZ2* strain growing on an agarose gel media pad, and (f) showing polar tubulation and budding-like processes. (g-i) Phase-contrast microscopy of *H. volcanii* strains containing expression plasmids, as indicated, sampled in steady mid-log growth in Hv-Cab medium with the indicated concentrations of Trp.

The genome sequence data for H98 and the three knock-out strains showed several small sequence variations of no known or expected consequences (Table S5). We therefore sought to verify the attribution of the above phenotypes specifically to *ftsZ1* and/or *ftsZ2* by reintroducing these genes under control of the *p*.*tna* promoter on a plasmid (based on pTA962) into the corresponding deletion mutants. In all three cases this successfully complemented the division defects in the presence of 0.2 mM Trp induction or greater (Fig. 2g, S8, S9). At relatively high concentrations of Trp (1 or 2 mM), the strains expressing *ftsZ1* showed some rods or spindle-like cell shapes (investigated further below).

The apparent correlation between the appearance of greatly enlarged cells and debris (< 1 µm^3^, Fig. 2c) suggested that the largest cells produce cell fragments or disintegrate to form the debris. Interestingly, *ΔftsZ1 ΔftsZ2* also showed a greater frequency of highly elongated filamentous cells (31%), compared to the plate-like cell shapes seen in mid-log cultures of the wild-type and two single *ΔftsZ* strains (< 1% filaments) (Fig. 2b, S7c). *H. volcanii* shapes can be readily influenced by several conditions or stresses^24^ (see Supplementary Results and Discussion) and the two major categories of giant cells appear to be the result of division defects in the common plate and rods morphotypes, respectively. The filamentous Δ*ftsZ1* Δ*ftsZ2* cells showed a similar flattened shape profile cells as the wild-type rod- and plate-shaped cells, which are normally ∼0.5 µm thick (Fig. 2d, Video S4). However, the giant plate-like cells had lost much of the flatness of the wild-type and displayed multi-lobed ramified 3D morphologies (Fig. 2d, S7b, Video S4), underscoring the remarkable plasticity of the *H. volcanii* envelope in the giant cells. This could lead to abiotic division or cell disintegration/fragmentation in a turbulent environment, and account for some of the highly irregular morphologies observed when cells are transferred from liquid to an agarose gel surface (*e*.*g*., Fig. 1a, 2b).

To further assess the capacity of cells to survive without *ftsZ*, including potential propagation via alternative biotic processes, mid-log samples were transferred to soft-agarose media pads to image cells during growth (Fig. 2e, S7d, Videos S5-S8). Under these conditions, the mutants grew substantially larger (over ∼12 h), indicating that cells were structurally stabilized by the gel compared to the prior liquid culture (Fig. 2c). Nevertheless, while some Δ*ftsZ1* cells grew very large (Video S5), others divided, often very acentrally or with a crooked division plane (Video S6). Division was not seen in Δ*ftsZ2* cells, which expanded to extremely large sizes, and rare blebbing was seen (Video S7). Interestingly, the filaments common in Δ*ftsZ1* Δ*ftsZ2* showed substantial polar narrowing and extension (tubulation), followed by fission to produce much smaller fragments (Fig. 2e-f, Video S8). These samples also contained many small particles, which may be the product of the polar tubulation-and-fission or fragmentation in prior culture. The fragmentation, budding, tubulation-and-fission, and, in the case of Δ*ftsZ1*, occasional inaccurate division, appear to result in particles, at least some of which contain DNA (see Fig. S5a), that produce enough viable cells for culture propagation in the absence of normal cell division.

### Overexpression of ftsZ1 or ftsZ2 cause contrasting phenotypes, and cannot properly compensate for loss of the other

If FtsZ1 and FtsZ2 have different roles in division, then overexpression of each could exacerbate aspects of each protein’s specific activities and potentially lead to different effects. Furthermore, if overexpression of *ftsZ2* in the *ΔftsZ1* strain, and vice versa, corrects the division defects (*i*.*e*., cross-complementation), then this would indicate similar or overlapping functions of the two proteins. FtsZ1 overproduction in the wild-type background caused enlarged cells, indicating inefficient or mis-regulated division (Fig 3). It also caused a change in cell shape, where cells frequently displayed a rod-like appearance or a noticeable taper at one or both poles (Fig. 3c). In contrast, FtsZ2 overproduction resulted in smaller cells than the wild type, indicating that FtsZ2 overproduction stimulates division, with no specific influence on cell shape (Fig. 3).

**Figure 3.**
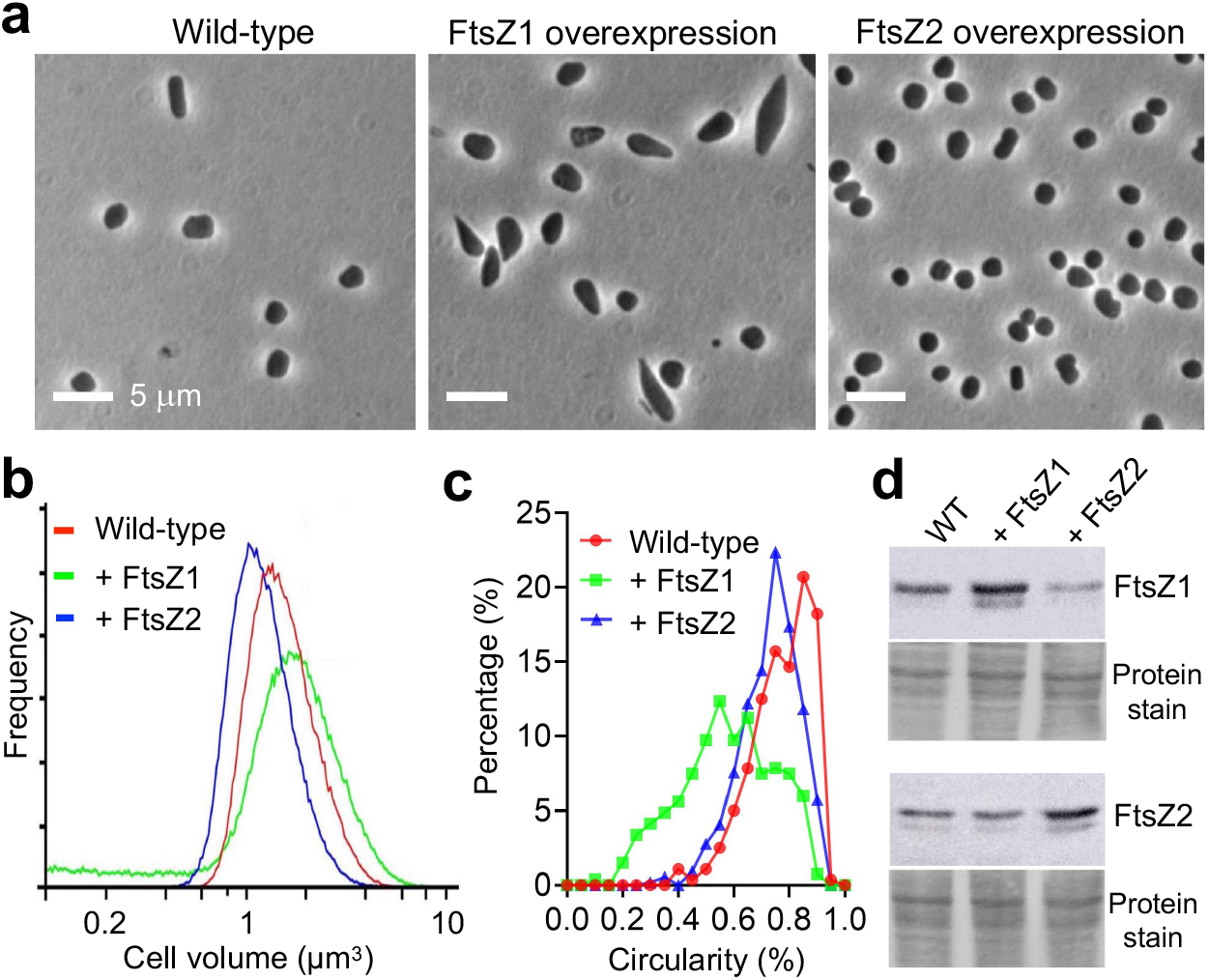
Overproduction of FtsZ1 or FtsZ2 differentially affects cell division and shape. (a) Phase contrast micrographs of *H. volcanii* wild-type (H98 + pTA962), ID25 (FtsZ1 overproduction, H98 + pTA962-*ftsZ1*) and ID26 (FtsZ2 overproduction strain, H98 + pTA962-*ftsZ2*) during steady mid-log cultures in Hv-Cab + 2 mM Trp. Scale bars, 5 µm. (b) Coulter cell-volume frequency distributions (normalized to total count) of the strains in panel (a). (c) Frequency distributions of the circularity of cell outlines (WT, n = 280; + FtsZ1, n = 267; + FtsZ2, n = 534). (d) Western blots of whole cell lysates of the wild-type, FtsZ1 overproduction and FtsZ2 overproduction strains, probed with FtsZ1 or FtsZ2 antisera, as indicated. Total protein pre-staining of each membrane (with Ponceau S) is shown as a loading control.

FtsZ1 overproduction provided no detectable recovery to the Δ*ftsZ2* division defect (Fig. 2h, S10), but caused increased debris and some striking spindle-like cell shapes with rather pointy poles or various angular morphologies, as well as very long filamentous cells of approximately normal rod-cell width. The spindly cells therefore appear to be associated with FtsZ1 overproduction in the wild-type, Δ*ftsZ1* and Δ*ftsZ2* backgrounds (Fig. 3a, 2g-h, S11). FtsZ2 overproduction caused weak complementation of the *ΔftsZ1* division defect (Fig. 2g-h, S8, S10), indicating partial compensation for the missing FtsZ1 or a capacity to take on some of the role(s) of FtsZ1 in division. These findings show that FtsZ1 and FtsZ2 have differing functions, and that FtsZ2 has an essential active role in the constriction mechanism, whereas FtsZ1 has a supporting role. Consistent with this, as *ftsZ1* mutant cultures slowed growth and entered stationary phase, cell size recovered to nearly normal, but this did not occur in *ftsZ2* mutants (Fig. S12). The stationary phase results and effects of *ftsZ* mutants on cell shape are further discussed in the Supplementary Results and Discussion.

### Predicted GTPase active site (T7 loop) residues are essential for FtsZ1 and FtsZ2 functions

FtsZ1 and FtsZ2 both contain the conserved GTP binding domain and catalytic residues in the T7 loop (see Fig. S1d) that are needed for GTP hydrolysis and correct function of FtsZ/tubulin polymers^31,32^. Bacterial FtsZ studies have showed that point mutations of the T7 catalytic residues block GTPase-dependent filament disassembly, thus forming hyper-stable filaments that can co-assemble with the wild-type protein and severely disrupt function^33-35^. Overproduction of an equivalent aspartate-to-alanine point mutant of FtsZ1 (D250A, substitution in the T7 loop) inhibited division in *H. volcanii*^20^. To determine whether both FtsZ1 and FtsZ2 require their predicted GTPase catalytic residues for their functions in division, we expressed *ftsZ1*.*D250A* and the equivalent *ftsZ2*.*D231A* mutant from plasmids in their corresponding Δ*ftsZ* backgrounds. These mutants completely failed to complement the division defects (Fig. 2i). The Δ*ftsZ1* + *ftsZ1*.*D250A* combination clearly exacerbated the Δ*ftsZ1* defect (Fig. 2b-c, S13a). Similarly, Δ*ftsZ2* + *ftsZ2*.*D231A* displayed a mix of giant plates and cell debris, but also showed filaments at the early-mid stages of culture (Fig. 2i, S13b). The additional phenotypes associated with expression of the two point-mutants suggested that they actively interfere with division or the cell envelope. Indeed, even moderate expression of *ftsZ1*.*D250A* or *ftsZ2*.*D231A* caused strong division defects in the wild-type background (Fig. S14). We therefore sought to investigate the subcellular localization and assembly of the wild-type and mutant proteins.

### Subcellular localization of FtsZ1 and FtsZ2

Genes encoding the fluorescent proteins GFP and mCherry (mCh) were fused to the C-termini of FtsZ1 and FtsZ2. FtsZ1-GFP or FtsZ1-mCh produced from plasmids (with 0.2 mM Trp) partially complemented the cell division defect of the Δ*ftsZ1* strain (Fig. S15a); many cells showed normal size and shape, with a fluorescent band at midcell or the division constriction, whereas others were larger and occasionally misshapen with more complex fluorescent structures near midcell. FtsZ2-GFP failed to complement the Δ*ftsZ2* division defect, but the highly filamentous cells showed sporadic FtsZ2-GFP foci or rings that were seen ∼3-4 µm from a pole (Fig. S15b), suggestive of the influence of a division site positioning mechanism^28^ that is sensitive to pole proximity.

Functional interference from fluorescent proteins is very common in FtsZ and tubulin^36-38^. Since the FtsZ fusion proteins were not fully functional in division as the sole copy, but did localize, we reasoned that they could be useful localization markers and not substantially perturb cell division in the presence of the wild-type FtsZ (as seen in bacteria). Indeed, low-level induction of FtsZ1 or FtsZ2 tagged with mCherry or GFP in a wild-type background did not disturb normal cell size and shape. FtsZ1-mCh localized as a band at midcell or the division constriction in almost all cells (Fig. 4a), with no evidence of perturbed division at 0.2 mM and 1 mM Trp (Fig. S15c). Similarly, FtsZ2-GFP localized at midcell or the division constriction in almost all cells, which appeared normal or slightly enlarged with 0.2 mM Trp (Fig. 4b, S15d). With 1 mM Trp, cells producing FtsZ2-GFP were substantially enlarged and some had aberrant FtsZ2-GFP clusters near the centre, consistent with their detachment from the envelope (Fig. 4b, S15d).

**Figure 4.**
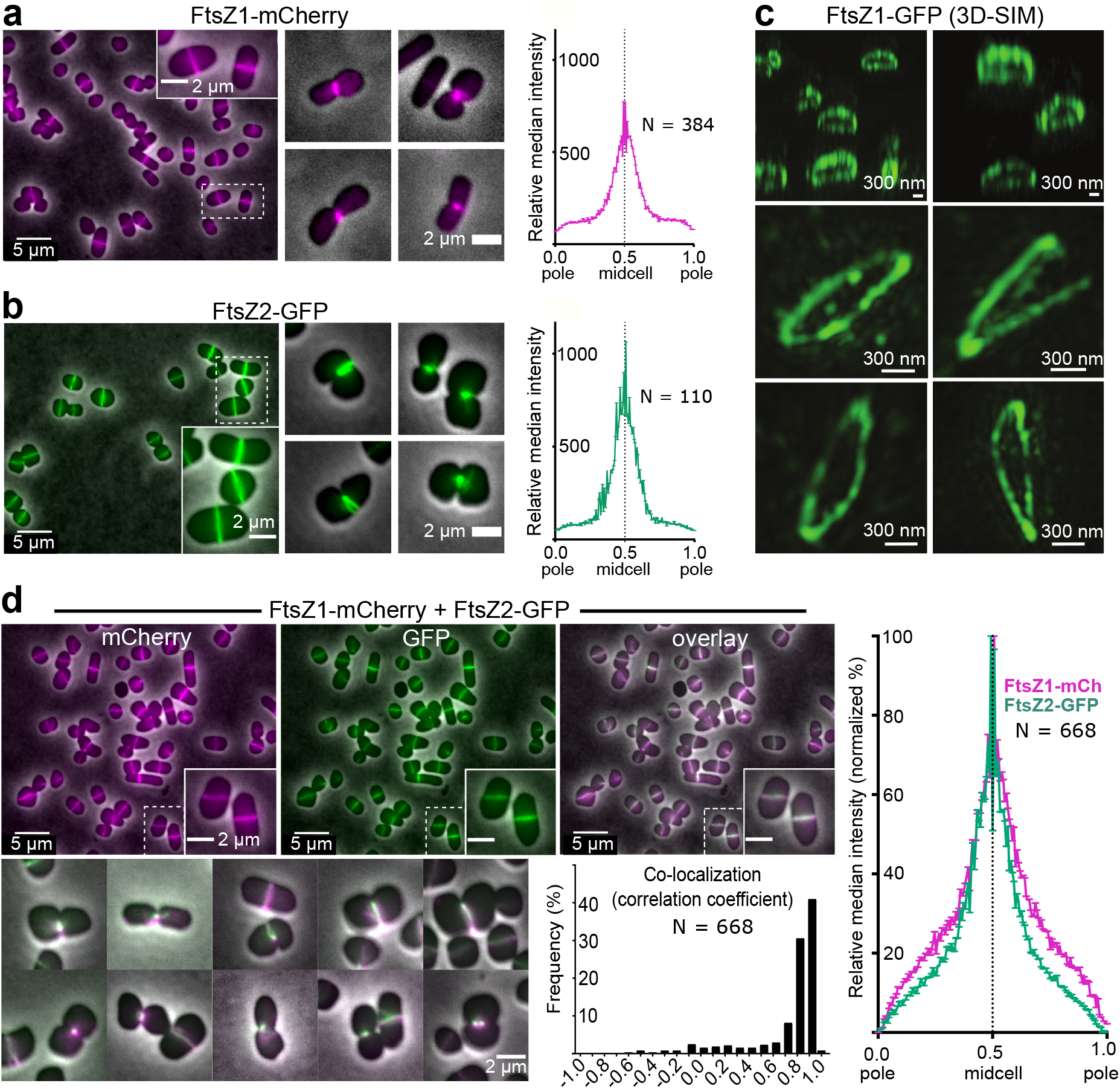
Midcell localization of FtsZ1-mCh and FtsZ2-GFP. Mid-log cultures of (a) *H. volcanii* ID49 (WT + FtsZ1-mCh), (b) ID17 (WT + FtsZ2-GFP) and (d) ID67 (WT + FtsZ1-mCh + FtsZ2-GFP dual localization), cultured with 0.2 mM Trp, were imaged by phase-contrast and fluorescence microscopy. The mCh (magenta) and GFP (green) fluorescence was quantified along the long axis of cells, by plotting the relative (normalized) median intensity perpendicular to the long axis. (c) 3D-SIM images of *H. volcanii* ID16 (WT + FtsZ1-GFP) sampled during mid-log growth with 0.2 mM Trp. The upper two panels show a ‘side-on’ view (∼70° tilt of the xy view), and the lower four panels show a ∼20° xy tilt. (d) Co-localization of FtsZ1-mCh and FtsZ2-GFP fluorescence in cells, including selected examples of dividing cells (lower left). Correlation coefficients were plotted as a frequency distribution (80% of the cells had a correlation coefficient greater or equal to 0.7). The ring containing FtsZ1 and FtsZ2 closes down with division constrictions that range between unilateral to partially and equally bilateral.

We confirmed the expected flattened ring-like structure of the midcell bands by imaging FtsZ1-GFP in the wild-type background with 3D structured-illumination microscopy (Fig. 4c, upper two images, ∼70° xy-tilt). This also revealed patchy, discontinuous rings or short helical structures (Fig. 4c, lower four images, ∼20° xy tilt), similar to the appearance of bacterial Z-rings^39,40^. Time-lapse imaging showed that the uneven fluorescence intensity of FtsZ1-GFP around the ring was dynamic during the cell cycle (Video S9, S10) and confirmed that prompt re-assembly of new rings occurs after division in orientations consistent with predictions from cell morphology^28^.

When produced together, FtsZ1-mCh and FtsZ2-GFP generally colocalized as a midcell band in essentially all cells (Fig. 4d, S15e), indicating that both proteins assemble into a midcell band very soon after the previous division and are maintained at midcell during cell growth. Co-localization was strong, with a correlation coefficient of >0.8 in the majority of cells but appeared noticeably imperfect in some (Fig. 4d). The two proteins contracted together during division, which varied from bilateral to unilateral (Fig. 4) and remained generally co-localized during reassembly over several cell cycles (Fig. 4d, Video S11).

### Differing interdependency of FtsZ1 and FtsZ2 localization

The differing functions of FtsZ1 and FtsZ2 in division, and their co-localization at midcell, prompted us to investigate the dependency of FtsZ1 and FtsZ2 on each other for cellular localization and assembly, by visualizing each FtsZ-FP in the alternate Δ*ftsZ* and T7-loop mutant backgrounds. The T7 mutations were expected to produce aberrant hyper-stable polymers and could help to resolve individual functions in divisome assembly or constriction.

In the absence of *ftsZ2*, FtsZ1-mCh displayed one or more clear rings (Fig. 5a) in the predominantly filamentous cells (62%, Fig. S16a). The localizations were spaced on average every ∼5 µm, and some appeared less condensed than normal rings—as a short helicoid or irregular structure (Fig. 5a, S17). Giant plate-like cells were also present (5%), which contained clusters of FtsZ1-mCh (similar to that seen in Fig. 5b). In the absence of *ftsZ1*, FtsZ2-GFP formed patches, foci or short filaments, but no rings (0.2 mM Trp, Fig. 5a). This strain showed a much more severe division defect than Δ*ftsZ1* and Δ*ftsZ1* + unlabelled FtsZ2 (Fig. 2g-h). To minimize this strong inhibitory effect of the FtsZ2-GFP, we used lower induction levels (0 or 50 µM Trp) and indeed observed some cells with poorly formed midcell structures, which appeared as more normal rings in dividing cells (Fig. 5a, S18). Since FtsZ2-GFP induced with 0.2 mM Trp had only a very minor effect on division when FtsZ1 is present (Fig. 4b, S13d-e), and at higher levels (1 mM Trp) it inhibited division and appeared to inhibit envelope association (Fig. S15d), FtsZ1 must counteract the destabilizing effect of FtsZ2-GFP on normal FtsZ2 assembly. Taken together, these results strongly suggest that FtsZ1 can assemble independently of FtsZ2 and promotes and stabilizes proper FtsZ2 assembly for division at the midcell envelope.

**Figure 5.**
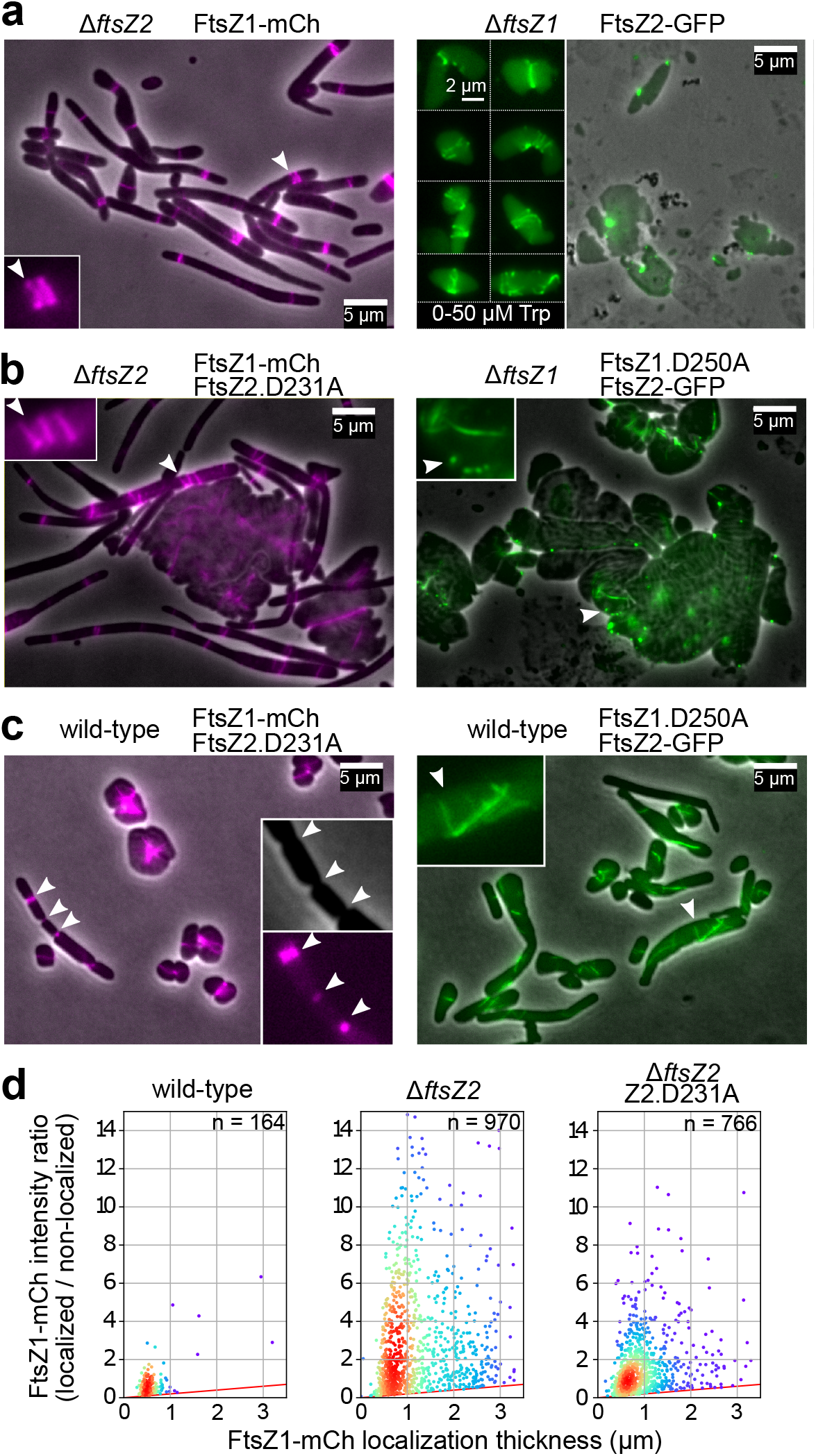
Localization interdependency and the effect of FtsZ mutations on FtsZ1-mCh and FtsZ2-GFP localization. Fluorescence and phase-contrast microscopy of mid-log *H. volcanii* strains (wild-type or Δ*ftsZ* backgrounds) carrying plasmid copies of the indicated *ftsZ* mutants and fusion proteins. Strains were grown with 0.2 mM Trp unless otherwise specified. Insets show either a magnified view at the arrowhead in the main panels, or represent another culture grown with the indicated concentration of Trp. (a) Localization of each FtsZ in the absence of the other FtsZ. (b) Localization of each FtsZ in the presence of the T7-loop mutant as the sole copy of the other. (c) Localization of each FtsZ in the wild-type genomic background expressing the T7-loop mutant of the other. Inset scale bars are 5 µm unless indicated otherwise. (d) Plots of FtsZ1-mCh localization relative intensity (per µm across the cell, as a ratio over the mean cellular background FtsZ1-mCh fluorescence) *versus* localization thickness (µm) in the indicated strain backgrounds. Data-points are coloured with a proximity heatmap. Representative images of the wild-type background can be seen in Fig 4a. See also Fig. S17.

When FtsZ2.D231A was produced as the sole copy of FtsZ2 (in Δ*ftsZ2*) with FtsZ1-mCh, highly elongated cells were observed (Fig. 5b, S16b), similar to the unlabelled strain (Fig. 2i), and FtsZ1-mCh localizations again appeared as rings, broader zones or helicoid structures (Fig. 5b). Measurements of the FtsZ1-mCh localization intensity (compared to cellular background) and individual localization thickness in the filaments (Fig. 5d) confirmed the lack of condensation of some of the FtsZ1 localizations in the FtsZ2 mutants and revealed that a substantially higher proportion of the total FtsZ1-mCh was localized in the absence of wild-type FtsZ2. These findings suggest that FtsZ2 contributes to proper divisome condensation or maturation and may limit the incorporation of FtsZ1 into the ring. In the less frequent giant plates (Fig. 5b), FtsZ1-mCh showed separate filaments or diffuse clusters or only rarely as an apparent ring spanning the cell, likely due to a lack of spatial cues for positioning the division machinery in the giant malformed cells^28^ and a low probability of assembling a ring of such abnormal size. When FtsZ1.D250A was produced as the sole copy of FtsZ1 (Fig. 5b), the cells formed giant plates as expected, and FtsZ2-GFP displayed patches or foci and some filaments, but no rings. Overall, the moderate influence of the sole T7-mutants on the other FtsZ’s localization pattern, compared to the respective knockout background (Fig. 5a), suggests that the sole-copy T7-mutant proteins do not efficiently homo-polymerize or do not efficiently interact (or interfere) with the other FtsZ.

Highly informative results were obtained with the wild-type background. In this setting, production of FtsZ2.D231A with FtsZ1-mCh partly inhibited division during constriction— the enlarged cells often displayed one or more incomplete constrictions with FtsZ1-mCh rings or occasional irregular structures present at those sites (Fig. 5c left). This strongly suggests that FtsZ2.D231A mixes with wild-type FtsZ2 to inhibit GTPase-dependent constriction, and that therefore the primary function of FtsZ2 and its GTPase activity is division constriction. In contrast, production of FtsZ1.D250A with FtsZ2-GFP in the wild-type background caused obvious aberrant filaments containing FtsZ2-GFP that were orientated seemingly at random or with the longer cell axis (Fig. 5c right). Clearly, the presence of wild-type FtsZ1 in this strain allows the assembly of aberrant filaments caused by FtsZ1.D250A that recruit FtsZ2-GFP (compare Fig. 5a-c right), leading us to conclude that FtsZ2’s subcellular localization is strongly influenced by FtsZ1. Furthermore, the absence of both midcell localization and partial constrictions (Fig. 5c right) indicates that the primary function of FtsZ1 and its GTPase activity is the proper assembly of the division machinery.

To our surprise, FP-tagging of the T7-mutants—in order to simultaneously visualize both the T7-mutant of one FtsZ and the wild-type copy of the other—suppressed their dominant-inhibitory phenotypes (Fig. S19a-b). However, the tagged T7 mutants still partly co-localized with the alternate wild-type FtsZ, and these additional experiments further confirmed the relative independence of FtsZ1 filaments and the capacity of FtsZ2 to provide some structural feedback in ring formation (see Supplementary Results and Discussion, and Fig. S19).

## Discussion

We have shown that archaeal FtsZ1 and FtsZ2, from distinct families within the tubulin superfamily of cytoskeletal/cytomotive proteins, both localize to the division ring but have different mechanistic roles. FtsZ1 assembles largely independently of FtsZ2 and directs the assembly, stabilization and correct localization of FtsZ2, whereas FtsZ2 is critical during constriction, including divisome activation or structure (see Fig. 6).

**Figure 6.**
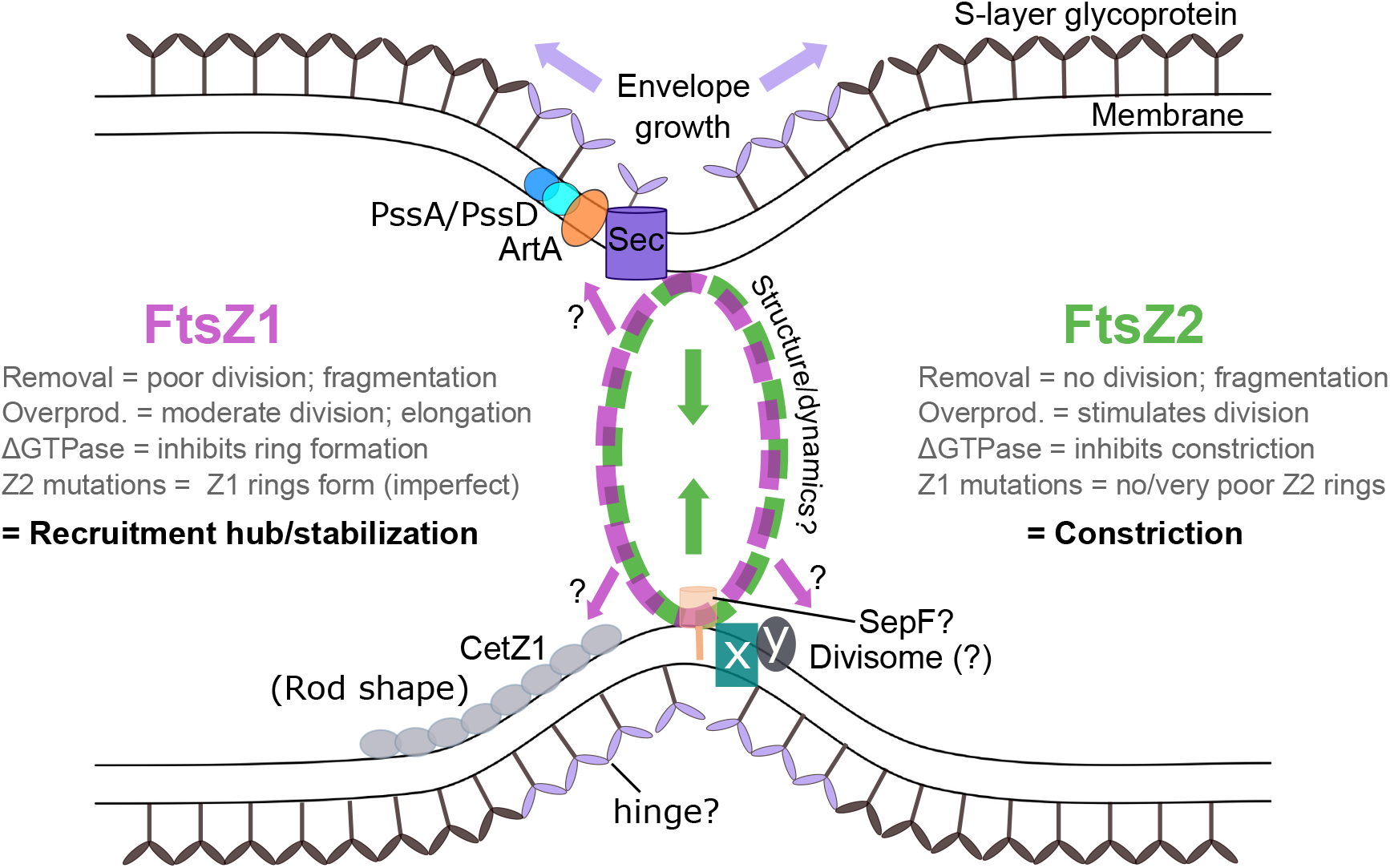
Model of FtsZ1 and FtsZ2 function at the archaeal division site. Schematic models of FtsZ1 (magenta) and FtsZ2 (green) localization and functions (arrows) are based on our results (summarized here). Most archaea, including *H. volcanii*, have an envelope composed of a lipid membrane and an S-layer glycoprotein semi-crystalline array on the surface. Our results raise questions about the ultrastructure and dynamics of the two FtsZ proteins in the division ring, the identity of an unknown number of other putative divisome components (x, y) including a SepF homolog, and how the ring could functionally associate with the envelope growth (Sec/PssA/PssD/ArtA) and cell shape regulation (CetZ1) machineries, which appear to be spatially and functionally linked to midcell^20,41^.

FtsZ1 was also capable of stimulating the formation of rod and tapered morphologies from the common plate/disc morphotype. *H. volcanii* rods also form during CetZ1 overproduction^20^, and CetZ1 can also localize at midcell or as filaments along the cell edge perpendicular to the division plane during its key function in rod development. Our findings suggest a possible linkage between FtsZ1 and CetZ1 function in promoting cell elongation (Fig. 6). Furthermore, nascent S-layer glycoprotein (SLG) and the PssA/PssD/ArtA enzymes required for its lipid modification and proper membrane anchoring were recently found to localize at midcell, and blocking this pathway caused rod formation but did not affect division^41^. These findings suggest a model in which FtsZ1 creates a scaffold or platform at midcell that acts as a hub or landmark to recruit and stabilize the division, shape and envelope biosynthesis machineries.

Despite these important functions in division and shape, *H. volcanii* lacking both FtsZs could be cultured indefinitely with standard solid and liquid media. FtsZ is thought to be essential for division and survival in almost all walled bacteria^25^, but under specific conditions of wall removal (‘L-forms’) or excess envelope—causing cell ramification—they can produce viable fragments and propagate without FtsZ^42,43^. *H. volcanii* Δ*ftsZ* strains appear similar (Fig. 2). Most archaea have a lipid membrane and a flexible glycoprotein S-layer but no cell wall as in peptidoglycan-enclosed bacteria^44^. Considering the morphological plasticity of wild-type *H. volcanii*^24^ and the Δ*ftsZ* strains, a turbulent liquid environment may be expected to promote physical fragmentation or disintegration of the giant cells, from which any particles with a complete genome may be viable and contribute to propagation without proper cytokinesis. We imagine that such ‘abiotic division’ could have helped sustain very early life on earth. Our results also suggested that budding and polar tubulation-fission might contribute to survival, and similar non-FtsZ modes of proliferation can occur in wall-less bacteria^45-48^. *H. volcanii*, and potentially other wall-less archaea, should therefore be suitable for studying primitive or alternative division modes and the normal cellular functions underlying them.

While the two FtsZs are conserved in a majority of surveyed archaea (Table S3), six of the 41 complete genomes encode only one FtsZ each. Remarkably, we noticed that the same six species, from the classes Methanobacteria and Methanopyri, are the only ones known to have a pseudomurein wall analogous to bacterial peptidoglycan (murein) (Table S3)^49,50^. Furthermore, these classes only encode FtsZ1 homologs. The archaeal dual FtsZ system therefore appears to have evolved after an early paralogous gene duplication, and then FtsZ2 was lost when pseudomurein evolved in those lineages.

In bacteria, division appears to be primarily driven by the FtsZ-directed ingrowth of the peptidoglycan cell wall^3,4^ and possibly an additional constriction force on the cytoplasmic membrane^5,6^. Electron microscopy of Methanobacteria and Methanopyri show a clear septal wall during division very similar in appearance to bacteria^51-53^. Archaea without pseudomurein show a division furrow instead^54,55^, and with only a single flexible S-layer (a 2D semi-crystalline protein array), the archaeal envelope is almost certainly not capable of providing the same directionality or structural persistence that remodelling of the covalent peptidoglycan mesh would in bacterial septal ingrowth^56^. Our experimental results and the clear phylogenetic association strongly suggest that FtsZ2 is required for constriction where there is no covalent wall ingrowth to impose directionality to division. We therefore predict that a division complex containing FtsZ2 at its core provides an intrinsic constriction force on the flexible archaeal envelope. With the low or no turgor pressure in wall-less archaea, especially halophilic archaea^57^, this force might be sufficient to complete division^7^.

Given the functional differentiation of two FtsZs in archaeal cell division, and the corresponding lack of a cell wall, we propose that *H. volcanii* provides a powerful model for mechanistic studies of cell division to contrast against the well-studied bacterial models with peptidoglycan. We anticipate that this will contribute to establishing the underlying and primordial principles of FtsZ-based division in all domains of life and help to unravel some of the mysteries of early cellular evolution and division.

## Methods

### Identification, phylogeny and analysis of FtsZ sequences and structure

Thirteen known FtsZ protein sequences from bacteria and plants were aligned with MUSCLE^58^ for use as the search set to identify homologs in a diverse collection of 60 archaeal genomes (Table S3) by searching their respective Reference Proteome or UniProt databases using JackHMMER^59^. All significant hits were selected, excluding several redundancies and low-quality sequences, and they were then aligned, together with the search set, using MUSCLE. The identification of homologs belonging to either the archaeal FtsZ1, FtsZ2, CetZ, or ‘other’ categories was done based on the position of individual sequences in a Maximum Likelihood phylogenetic tree, obtained using MEGA (ver. 7.0.26)^60^ with 100 bootstrap replicates, based on the alignment containing only those sites with >25% occupancy. *H. volcanii* FtsZ1 (HVO_0717), FtsZ2 (HVO_0581) and CetZ1 (HVO_2204) were taken as the reference sequences for identifying the three families.

To determine the percent sequence identities for each of the four domains, as defined for *H. volcanii* FtsZ1 and FtsZ2 in Fig. S1b, the sequence alignment generated above was arranged into protein families, including only those proteins classifying as FtsZ1, FtsZ2 or CetZ, and was then separated into individual domain alignments using Jalview (ver. 2.9.0b2)^61^. Each set was individually realigned with MUSCLE and used as input for generating a ClustalX percent identity matrix^62^ that was used for determining means. The crystal structure of *M. jannaschii* FtsZ1^63^ (PDB code: 1FSZ) was displayed with PyMol (ver. 1.7).

### Growth of H. volcanii

Most experimental *H. volcanii* cultures were grown in Hv-Cab medium^20^ or Hv-YPC medium for some genetic modification procedures^64^. Where necessary as auxotrophic requirements, media were supplemented with uracil (10 µg/ml or 50 µg/ml) for *ΔpyrE2* strains, or thymidine and hypoxanthine (40 µg/ml each) for *ΔhdrB* strains. Cultures were incubated at 45°C with rotary shaking (200 rpm) and were generally maintained in continuous logarithmic growth (OD_600_ < 0.8) for at least 2 days prior to sampling for analysis of mid-log cultures, unless otherwise indicated. To control gene expression via the *p*.*tna* promoter, the indicated concentration of L-tryptophan (Trp) (Sigma), was included in these cultures.

### Genomic modification

The strains used in this study and plasmids used to construct them as appropriate are listed in Tables S1 and S2, respectively. In order to generate strains allowing tryptophan-controlled expression of *ftsZ1* or *ftsZ2* for depletion studies (Fig. S2), two respective plasmids were first constructed that can recombine with each corresponding genomic *ftsZ* locus using two-step homologous recombination^64^ thereby substituting the normal transcriptional regulation of *ftsZ* with the specific tryptophan-inducible *p*.*tna* promoter. First, the upstream and downstream flanks for homologous recombination on either side of each *ftsZ* gene’s start codon were PCR amplified from *H. volcanii* DS2 genomic DNA (upstream flank) and the corresponding pIDJL40-*ftsZ* plasmid (downstream flank, containing the L11e transcription terminator and *p*.*tna-ftsZ* cassette to be inserted), by using the primers given in Table S2. The upstream and downstream fragments in each case were joined, by overlap-extension PCR, and the products digested with HindIII and BamHI (sites located in the end primers; Fig. S1 and Table S2) and ligated to pTA131 (at HindIII-BamHI), giving pIDJL74 (*p*.*tna-ftsZ1*) and pIDJL75 (*p*.*tna-ftsZ2*). For *ftsZ1*, the sequence of the chosen upstream flank would result in a 39 bp deletion of genomic DNA immediately upstream of the *ftsZ1* start codon during the replacement. For *ftsZ2*, which is located as the second gene in a predicted operon (downstream of HVO_582, with a 2 bp gap), no genomic DNA was deleted when inserting the *p*.*tna* construct upstream of the *ftsZ2* ORF. To allow direct selection of the *p*.*tna-ftsZ* genomic replacements, the BamHI fragment from pTA1185 (T. Allers, personal communication), containing *p*.*fdx-hdrB*^64^ was inserted at the BglII site in pIDJL74 and pIDJL75 (*i*.*e*., between the upstream and downstream fragments, described above). Clones containing the *p*.*fdx-hdrB* co-directional with the downstream *p*.*tna-ftsZ* cassette were selected and named pIDJL96 (*ftsZ1*) and pIDJL97 (*ftsZ2*) (Table S2).

Demethylated pIDJL96 and pIDJL97 (obtained via passage through *E. coli* C2925) were used to transform *H. volcanii* H98 separately, selecting isolated colonies that grew on agar medium (without uracil), which were expected to contain the plasmid integrated by single-crossover (“pop-in”)^64^ between the upstream or downstream flank and the corresponding genomic *ftsZ* locus. After growth of single colonies in liquid media (minus uracil), cells were plated onto Hv-Ca agar containing 10 µg/mL uracil, 50 µg/mL 5-fluoroorotic acid (FOA) and 0.5 mM Trp, to select for recombinational excision of these non-replicative plasmids (“pop-out”)^64^ that following gene conversion left the cassette containing *p*.*fdx-hdrB* and *p*.*tna-ftsZ* as the only copy of *ftsZ*; single colonies were streaked onto the same medium, and then arising colonies were screened by allele-specific PCR to verify the expected chromosomal structure, giving strains ID56 (*p*.*tna-ftsZ1*) and ID57 (*p*.*tna-ftsZ2*) (Table S1).

To construct plasmids for deletion of each *ftsZ* gene, PCR fragments of downstream flanking DNA for *ftsZ*1 and *ftsZ2* were amplified (primers shown in Table S2). The products were digested with BglII and BamHI and then ligated respectively to the large fragments of pIDJL74 or pIDJL75 digested with BglII, BamHI and alkaline phosphatase. Isolated clones containing the appropriate co-orientation of up- and down-stream flanks were selected and named pIDJL128 (*ftsZ1* flanks) and pIDJL129 (*ftsZ2* flanks) (Table S2). These plasmids were then digested with BglII (*i*.*e*. cutting between the flanks) and then the *p*.*fdx-hdrB* BamHI fragment from pTA1185 was ligated to them, generating pIDJL142 (for *ftsZ1* deletion) and pIDJL143 (for *ftsZ2* deletion).

To delete *ftsZ* and/or *ftsZ2* in *H. volcanii*, plasmids pIDJL142 and pIDJL143, respectively, were demethylated as described above, and then used to transform *H. volcanii* H98 using the two-step procedure noted above^64^, yielding strains ID76 (i.e. (H98) *ΔftsZ1-p*.*fdx-hdrB*), and ID77 (i.e. (H98) *ΔftsZ2-p*.*fdx-hdrB*). Then, to construct the double-knock-out strain, *H. volcanii* ID77 was transformed with demethylated pIDJL128 (Table S2) using the two-step method described above, generating *H. volcanii* ID112. The expected mutations were verified by allele-specific PCR and genome sequencing, as described below, to detect the presence of the expected *ftsZ*-deletion allele and the absence of the wild-type allele(s). In addition, the expected lack of the corresponding proteins was verified with western blotting using specific FtsZ1 or FtsZ2 antibodies (Fig. 1, S8d).

### Genome sequencing and variant calling

Whole genome sequencing libraries for strains H98, ID76, ID77 and ID112 were prepared using the Illumina Nextera Flex DNA kit with custom index primers. Sequencing fragment libraries were then pooled and size-selected using the SPRI-Select magnetic beads (Beckman Coulter) and the pooled library was then quality checked and quantified with a Agilent Bioanalyzer 2100 using the DNA HS kit (Agilent). Sequencing of was performed using an Illumina MiSeq with a 300-cycle micro flow cell with V2 chemistry. Sequence data were de-multiplexed using Phylosift^65^ and trimmed using Trimmomatic with default settings^66^. A draft H98 genome was assembled using the A5 miseq assembler (version 20150522)^67^, and then the trimmed reads for the three Δ*ftsZ* strains were aligned to the draft H98 genome using bwa^68^. Variant calling was done using bcftools mpileup and call^69^. Any single nucleotide polymorphisms (SNPs) that had a quality score of less than 20 or were located within contigs of less than 1000 bp were discarded. Non-synonymous variants in coding regions or any variants in intergenic regions were then identified in ID76, ID77 and ID112 compared to the H98 draft reference genome (Table S5).

### Construction of plasmids for gene expression in H. volcanii

All plasmids used are listed in Table S2. Plasmids pTA962^70^ or pIDJL40 (containing GFP)^20^ were used as the basis to construct plasmids for controlled expression of the *ftsZ* genes or modified versions. To construct pTA962-*ftsZ1*, the *ftsZ1* ORF (HVO_0717) was amplified from *H. volcanii* DS70 genomic DNA using primers *ftsZ1-f* and *ftsZ-r* (Table S2), and then the product was cloned between the NdeI and BamHI sites of pTA962. pTA962-*ftsZ2* was constructed in the same way, except with primers *ftsZ2-f* and *ftsZ2-r* to amplify the *ftsZ2* ORF (HVO_0581) (Table S2). The *ftsZ1*.*D250A* mutation was constructed previously^20^, and the *ftsZ2*.*D231A* mutation was similarly made by overlap extension PCR using the mutational primers and primers *ftsZ2-f* and *ftsZ2-r* (Table S2) and the product cloned between NdeI and BamHI in pTA962.

The wild-type and mutant *ftsZ* ORFs were also amplified without their stop codons (‘NS’ primers; Table S2) and then cloned between the NdeI and BamHI sites of pIDJL40, to create *ftsZ-gfp* fusions (linker encoding GS, derived from the BamHI site). Then, to construct *ftsZ1-mCh* and *ftsZ2-mCh* fusions (pIDJL114 and pIDJL115, respectively), the GFP fragments in pIDJL40-*ftsZ1* and pIDJL40*-ftsZ2* were replaced by a BamHI-NotI fragment encoding a linker (GSAGSAAGSGEF, including the initial BamHI site and ending EcoRI site) and mCh (Table S2), as described previously^20^. To construct plasmids for expression of the mutant *ftsZ* genes tagged with mCh, NdeI + BamHI fragments (containing the point-mutant *ftsZ* genes without their stop codons) were ligated to the large fragment of pIDJL117 (*i*.*e*. expression of *cetZ1-mCh*) digested with NdeI and BamHI, thus replacing the *cetZ1* with the mutant *ftsZ* (Table S2). The expected DNA sequences of all cloned PCR products and point mutations in this study were verified by Sanger sequencing.

For dual expression of the various *ftsZ* genes, PvuII-fragments from the above plasmids (containing a *p*.*tna-ftsZ* gene fusion) were ligated into HindIII-cut (Klenow blunt-ended) plasmids containing the other *ftsZ* gene for the required combination, creating plasmids with two independent copies of the different *p*.*tna-ftsZ* fusions; the combinations made are given in Table S2, where the second-listed gene in the description refers to the one transferred secondarily in the PvuII fragment. All plasmids were demethylated by passage through *E. coli* C2925 and re-purified prior to transfer to *H. volcanii* by PEG-mediated spheroplast transformation^64^.

### Growth curves (microtiter plate) and colony (cfu) counting

Cultures maintained in log growth for at least 2 days were diluted to an OD_600_ of 0.005 with fresh medium, and then 150 µL volumes were dispensed into a BD Falcon 96-well culture plate per well. Absorbance (OD_600_) of the samples over time was measured with a Tecan Spark 10M microplate spectrophotometer at 42°C with an orbital shaking (amplitude 2.5 mm, 216 rpm), with readings every 30 min for 56 h. Representative data from one of four independent experiments were then plotted, displaying standard error bars for technical replicates of the OD_600_ at each timepoint. Colony forming units (cfu) were determined by taking mid-log samples of each strain at OD_600_ = 0.2, and then plating 100 µL volumes of serial dilutions onto Hv-Cab (+ 50 µg/mL uracil) agar, and colonies were then counted after 4-5 days incubation in a sealed bag at 45°C.

### Microscopy

For transmitted light and epifluorescence microscopy, a 1 or 2 μL sample of culture was placed on a 1% agarose pad containing 18% buffered saltwater (BSW) on a glass slide at room temperature, and a #1.5 glass coverslip was placed on top. Images were acquired using a 1.4 NA oil immersion objective with phase-contrast or differential-interference contrast transmission optics, or by epifluorescence with GFP (Ex: 475/28 nm; Em: 525/48 nm) or mCh (Ex: 632/22 nm; Em: 676/34 nm) filter sets. 3D structured-illumination microscopy (3D-SIM) was performed as previously described^20^.

For live-cell time-lapse imaging, the ‘submerged-sandwich’ technique for growth of cells on agarose pads^20^ or a microfluidics platform were used. For agarose pads, 1-2 μL of mid-log culture was placed on a ∼1 mm thick gel pad (0.3% w/v agarose, containing the complete medium for the required conditions), that had been prepared on an 8 mm diameter #1.5 circular glass coverslip (WPI). The coverslip-pad-sample assembly was then lifted with forceps and placed, inverted, onto the base of a 35 mm glass-based (#1.5) FluoroDish (WPI). Three millilitres of pre-warmed liquid medium were then gently applied to cover the pad assembly on the base, and the lid was applied to avoid evaporation. For microfluidics, A CellASIC ONIX system was used with bacterial microfluidic plates (B04A, EMD Millipore). The flow cells were first equilibrated with 1 mg/mL Bovine Serum Albumin (BSA) in phosphate-buffered saline followed by 18% BSW at constant flow pressure of 5 psi. Cells were loaded into the chamber and perfused with the indicated medium and pressure. Time-lapse imaging was performed at 42°C or 45°C on a Nikon Ti-E microscope with a NA 1.45 oil-immersion phase-contrast objective, maintaining focus automatically during time-lapse.

For 3D confocal imaging, live cells were stained by addition of cytoplasmic vital stain Mitotracker Orange (1 µM)^71^ to mid-log cultures ∼18 h (45°C with shaking) prior to washing the cells three times and resuspending in fresh medium (without stain), or by addition of membrane dye FM1-43 (5 µg/mL) to culture samples at 45°C for 15 min. The stained cells were then gently mixed with an equal volume of molten (45°C) 1% low-melting-point agarose in fresh growth medium, and the mixture was placed onto the base of a FluoroDish and the environment was maintained humidified with a wet tissue during imaging at 37 or 45°C. A Nikon A1 laser-scanning confocal microscope was used with a 488 nm laser z-step of 0.2 μm. Each 3D confocal series was deconvolved by NIS-elements software (Nikon) using blind deconvolution (20 iterations). The 3D volume visualization was rendered using depth-coded alpha blending to display the depth information, followed by the blind deconvolution process for 20 iterations. The 3D video animations were created by rotation around Y-axis with a display rate of 20 frames per second.

### Image analysis

To determine cell circularity, phase-contrast images were first smoothed using a Gaussian filter followed by a rolling-ball background subtraction in FIJI^72^. Individual objects (cells) were then identified by thresholding and filling any holes. Clearly touching cells were manually separated. The minimum cell area was 0.2 μm^2^ and objects overlapping the image edge were excluded. The cell area was obtained from the *analyze particles* function, and cell circularity was calculated by determining each cell’s bounding disc coverage (the percentage area of the cell within the minimal circle that completely contains the cell outline^20^) with a custom script in FIJI. Cells and particles were classified as filaments (cell area > 7.5 μm^2^, Circularity < 0.3), giant plates (Cell area > 7.5 μm^2^, Circularity > 0.3), wild-type-like cells (any shape, with cell area between 1.5 μm^2^ and 7.5 μm^2^), or cellular debris (< 1.5 μm^2^), as indicated in the graphs.

FtsZ1 and FtsZ2 medial localization along the cell long axis (Fig. 4) was then done with FIJI^72^ and MicrobeJ^73^. Fluorescent images were pre-treated with the background subtraction filter (ballsize = 8) in FIJI. Cell outlines were detected in MicrobeJ by phase-contrast image segmentation using the Otsu method, with medial axis mode, area >0.98, and exclude-on-edges options, and then manually corrected where needed. FtsZ foci were detected using the ‘Foci’ and ‘Smoothed’ modes with a tolerance of 160, a Z-score of 8, and area >0.005. Raw pixel intensities and medial profiles were recorded. Data were output using the MicrobeJ XStatProfile Plot function with the parameters: Stat: Median, Y axis: MEDIAL.intensity.ch1 (and 2 for colocalization), bin#:150. The correlation coefficients for individual cells for FtsZ1 and FtsZ2 fluorescence were obtained using the MicrobeJ statistics functions, Stat.: count, Data: EXPERIMENT.count.total. Split horizontal: INTENSITY.ch2.correlation.ch1.

To determine FtsZ1-mCh localization frequency, intensity and thickness (Fig. 5, S17), cell outlines (regions of interest, ROIs) were first obtained from the phase-contrast channel in FIJI, by combining automated detection and manual curation. The long medial axis for each cell was then identified and the fluorescence channel data (after image/sensor background subtraction) were then quantified by averaging the intensity on the transverse axis to create a longitudinal intensity profile. Gaussian peaks were fitted to the significant localizations and a spline fit to the cellular Z1-mCh fluorescence background. The localization thickness was taken as the width of the fitted Gaussian peaks at half height (µm), and the intensity was taken as the integrated peak area (per µm across the cell).

### Coulter and flow cytometry

For Coulter cytometry to obtain cell volume distributions, culture samples were diluted (1:100 or 1:1000) with 0.2-µm-filtered 18% BSW and were analysed with a Multisizer M4 Coulter cytometer (Beckman-Coulter) equipped with a 20 or 30 µm aperture tube, running and calibrated with 2 µm latex beads in 18% BSW as the electrolyte. Runs were completed in the volumetric mode (100 μL), with a current of 600 μA and gain of 4. For flow cytometry, mid-log culture samples were diluted 1:10 with a solution of 5 µM SYTOX Green in 18% BSW and incubated at room temperature for ∼30 min before log-scale data acquisition (triggered by a side-scatter threshold) with an LSR-II flow cytometer (BD) with 18% BSW as sheath fluid. Cell sampling size normally ranged between 30,000 to 100,000 events per sample.

### Western blotting

Rabbit antisera for detection of *H. volcanii* FtsZ1 and FtsZ2 were generated with a synthetic peptide antigen derived from the sequence of the C-terminal region of FtsZ1, (QAHAEERLEDIDYVE, Cambridge Research Biochemicals, UK), used at 1:1000 dilution of serum. For FtsZ2, antibodies were raised against a peptide containing an N-terminally derived sequence (ERQTQSSLEDSDDQFGDPR, Thermo Fisher, USA) (1:500 dilution of serum), and one derived from the C-terminal region (SDGGRDEVEKNNGLDVIR, Thermo Fisher, USA) (1:4000 dilution of affinity purified antibody stock of 2.95 mg/ml). The FtsZ2 antibodies were used either as diluted serum or affinity purified protein (selecting for binding for the target peptide), as follows. N-term (serum): Fig. 1b, S6c, S8d, and C-term (affinity purified): Fig. 3d.

*H. volcanii* cell pellets (carefully separated from any excess liquid medium) were resuspended in SDS–PAGE sample buffer and then the samples were heated (95°C, 5 min) and vortexed. Samples were separated by SDS-PAGE, and then electroblotted (Bio-Rad) on a nitrocellulose membrane (Protran, Whatman), and probed with the rabbit polyclonal primary antibodies described above, followed by secondary antibody (donkey anti-rabbit IgG-HRP, AbCam 16284, 1:5000 dilution), with standard techniques. Bands were detected by enhanced chemiluminescence reagents (Thermo Fisher) with an Amersham Imager 600 system.

### Experimental replication and data and materials availability

In addition to the preliminary experiments carried out throughout this study, the data displayed are representative of at least two biological replicate experiments, and the displayed microscopy images and accompanying descriptions were selected to fairly represent the whole sample dataset, which was quantitatively analysed as described and represented in full. The data and biological materials that support the findings of this study are available from the corresponding author upon reasonable request. Genome sequence data has been deposited at NCBI under BioProject PRJNA681931.

## Acknowledgments

This study was supported by the Australian Research Council (FT160100010 to IGD) and the UK Biotechnology and Biological Sciences Research Council (BBSRC) and Medical Research Council (U105184326 to JL). For technical support, we thank Lynne Turnbull, Michael Johnson and Louise Cole from the UTS Microbial Imaging Facility (MIF), and Kay Anantanawat and Michael Liu from the UTS DNA sequencing facility.

## Author Contributions

Designed research: ID, SI, YL, JL. Preliminary data and strain construction: ID, SI, YL. Contributed/analysed data: Fig. 1: YL, Fig. 2: YL (b-i), SI (a), Fig. 3: YL, Fig. 4: SI (a, b, d), ID (c), Fig. 5: SI (a-d), YL/ID (a), CE (d), Fig. 6: SI/YL/ID. Supplementary Figures: S1: ID, S2: YL/ID, S3: YL, S4: YL, S5: ID, S6: YL, S7: YL/CE, S8: YL, S9: YL, S10: YL, S12: YL, S13: YL, S14: SI, S15: SI, S16: SI/CE, S17: CE/SI/ID, S18: YL/SI, S19: SI. Supplementary Tables: S1: ID/SI, S2: ID/SI, S3: ID, S4: ID, S5: YL, S6: SI/YL. Supplementary Videos: S1-S8: YL, S9-S10: ID/YL, S11: SI. Interpreted results, wrote and reviewed the manuscript and figures: ID, YL/SI, JL, CE. Management, funding acquisition and supervision: ID, JL.

## Declaration of Interests

The authors declare no competing interests.

## Supplementary Information

Supplementary videos, and a file containing supplementary results and discussion, tables, figures, video legends and references are available with the online version of this article. Genome sequence data has been deposited at NCBI under BioProject PRJNA681931.

## Notes

### Competing Interest Statement

The authors have declared no competing interest.

### Summary of Updates

Shorter, revised version. Some new data included.

