## Supplementary Information file for "Two FtsZ proteins orchestrate archaeal cell division through distinct functions in ring assembly and constriction"

**Contents:**

**Supplementary Results and Discussion**

**1.** Characteristics of two distinct clades of FtsZ in archaea p2
**2.** Domain arrangement and structure of FtsZ1 and FtsZ2 p2
**3.** Different consequences of *ftsZ1* and *ftsZ2* mutations in stationary phase p3
**4.** Different cell shapes of *H. volcanii* in various *ftsZ1* and *ftsZ2* mutant backgrounds p3
**5.** FP-tagging of the T7-mutants p4

**Supplementary Figures**

**Figure S1**. Molecular phylogeny and comparison of archaeal FtsZ1 and FtsZ2 families p5
**Figure S2.** Construction of *p.tna-ftsZ* depletion strains p6
**Figure S3.** Depletion of FtsZ1 and FtsZ2 in microfluidic chambers p7
**Figure S4.** Growth curves and cell size distributions during depletion of FtsZ1 and FtsZ2. p8
**Figure S5**. Cellular DNA content during depletion of FtsZ1 and FtsZ2 p9
**Figure S6**. Inducer (Trp) concentration-dependence of cell size in FtsZ-depletion strains p10
**Figure S7**. Phenotypes of Δ*ftsZ* strains p11
**Figure S8**. Complementation of the single Δ*ftsZ* strains p12
**Figure S9**. Complementation of the Δ*ftsZ1* Δ*ftsZ2* strain p13
**Figure S10**. FtsZ overproduction fails to properly complement knockout of the alternate *ftsZ* p14
**Figure S11**. Analysis of cell shape during *ftsZ*1/2 overexpression in various *ftsZ* mutant backgrounds p15
**Figure S12**. Comparison of *ftsZ* mutant cellular phenotypes in mid-log and stationary phases p16
**Figure S13**. Effects of GTPase active-site (T7) mutants during the growth cycle p17
**Figure S14**. Dominant-inhibitory phenotypes of T7 mutants *ftsZ1.D250A* and *ftsZ2.D231A* p18
**Figure S15**. Analysis of the function of FtsZ1 and FtsZ2 fluorescent fusions p19
**Figure S16**. Cell shape analyses for the localization interdependency studies. p20
**Figure S17**. FtsZ1-mCherry ring frequency and thickness in FtsZ2 mutant strains. p21
**Figure S18**. Localization of FtsZ2-GFP in the absence of *ftsZ1* p22
**Figure S19**. Localization studies of FtsZ T7-loop mutants, *ftsZ1.D250A* and *ftsZ2.D231A* p23

**Supplementary Tables**

**Table S1**. Strains used in this study p24
**Table S2**. Plasmids and oligonucleotides used in this study p25
**Table S3**. Number of tubulin superfamily sequences identified in the indicated archaea p26
**Table S4.** Percent identity amongst domains of the archaeal FtsZ1, FtsZ2 and CetZ protein families p27
**Table S5.** Small sequence variants in *H. volcanii* H98, Δ*ftsZ1*, Δ*ftsZ2*, and Δ*ftsZ1*Δ*ftsZ2*. p28

**Supplementary Video Legends**

**Video S1**. Time-lapse microscopy of FtsZ1 and FtsZ2 depletion and restoration p29
**Video S2**. Time-lapse microscopy of division/fragmentation of FtsZ1-depleted cells p29
**Video S3**. Time-lapse microscopy of fragmentation of FtsZ2-depleted cells p29
**Video S4**. 3D imaging of *H. volcanii* wild-type and Δ*ftsZ1* Δ*ftsZ2* strains p29
**Video S5**. Time-lapse microscopy of Δ*ftsZ1* – expansion of giant plates on agarose p29
**Video S6**. Time-lapse microscopy of Δ*ftsZ1* – occasional division/fragmentation p29
**Video S7**. Time-lapse microscopy of Δ*ftsZ2* – expansion of giant cells p29
**Video S8**. Time-lapse microscopy of Δ*ftsZ1* Δ*ftsZ2* – expansion of giant plates and polar tubulation of filaments p29
**Video S9**. Time-lapse microscopy of FtsZ1-GFP during multiple rounds of division p29
**Video S10**. Time-lapse microscopy of FtsZ1-GFP shows dynamic behavior in midcell rings p29
**Video S11**. FtsZ1 and FtsZ2 dynamically co-localize at midcell during division p29

**Supplementary References (p30)**

### Supplementary Results and Discussion

#### 1. Characteristics of two distinct clades of FtsZ in archaea

To identify features of the two archaeal FtsZ families and their distribution in well-known and recently discovered archaea, we surveyed the Archaea domain for the presence of FtsZ homologs in 60 complete or draft genomes of a diverse selection of species. This identified 149 non-redundant sequences from the broader tubulin superfamily, *i.e.,* FtsZ, CetZ, tubulin, or other non-canonical relatives (Table S3). Forty-nine of the surveyed genomes (including incomplete genomes) encode at least one FtsZ, 36 have at least two FtsZ proteins, and 33 have at least one from each of the FtsZ1 and FtsZ2 families (Table S3).

The five complete genomes analysed from the archaeal phylum Methanobacteria and the one from Methanopyri have only one (FtsZ1). Some non-canonical deeply branching FtsZ-like sequences present in Thaumarchaeota, Korarchaeota and others of unknown function, as well as the CetZ family (involved in cell shape), were also identified. The Crenarchaeota and Thaumarchaeota lacked specific members of the FtsZ families. Thaumarchaeota contain several non-canonical tubulin-superfamily proteins that are unlikely to function in division[1], but as expected most Crenarchaeota do not contain any[2-4]. FtsZ was also not found in other species from *Candidatus* phyla Marsarchaeota and Verstraetearchaeota in the TACK superphylum, although FtsZ was identified in some other TACK groups such as Bathyarchaeota (Table S3).

Interestingly, the unusual archaeon *Thermoplasma acidophilum* encodes two different, deep branching FtsZ sequences but no clear FtsZ2 (Fig. S1a, Table S3). *Thermoplasma* have a unique glycolipid envelope with no pseudomurein or S-layer[5], raising the question of whether the unusual FtsZ pair in *Thermoplasma* evolved independently to play a role analogous to the common FtsZ1/2 system described here, or whether an alternative or additional cytoskeletal system exists for division[6]. Other sporadic examples of multiple FtsZ or FtsZ-like proteins can be seen in archaea (Table S3) and bacteria, although the additional FtsZ proteins appear not to function in division or have as yet unknown functions[7,8]. Two plant FtsZ subfamilies within the major bacteria/plant family (Fig. S1a) are involved in division of plant plastids, and these FtsZ proteins show functional differentiation during division of these multi-layered organelles derived from cyanobacteria[9], raising the possibility that the plant and archaeal multi-FtsZ systems share some analogous activities related to division of a relatively flexible, multi-layered envelope.

#### 2. Domain arrangement and structure of FtsZ1 and FtsZ2

*Sequence similarity by domain*

*H. volcanii* FtsZ1 (HVO_0717) and FtsZ2 (HVO_0581) have the same overall domain arrangement as bacterial FtsZ. In the core fold of FtsZ, containing the GTP-binding and polymerization domains, the two *H. volcanii* FtsZ proteins each share high average sequence identity with other members of their respective families across the full diversity of archaea surveyed (Fig. S1b, percentages within each). Comparison of domains between the two *H. volcanii* proteins (Fig. S1b, %ID) showed lower sequence identity (46% overall), indicating conserved differences between the families. This was also seen by comparing average percent identities amongst all proteins belonging to the archaeal FtsZ1, FtsZ2, and CetZ families (Table S4). Closer inspection of the sequences revealed conserved differences between the families, described below.

*N-terminal tails*

An N-terminal extension (or 'N-tail') to the core fold is present in both FtsZ1 and FtsZ2 (Fig. S1b), confirming this region as a characteristic feature of the broader FtsZ family, compared to CetZ and tubulin that generally lack the N-tail. FtsZ1 and FtsZ2 have a conserved amphipathic ~9 amino-acid (aa) sequence of unknown 3D structure at the very N-terminus that appears to have very limited similarity to the highly variable bacterial N-terminal extension[10]. In FtsZ1, the consensus from our alignment was $M_{78}(D/K)_{52}S_{54}I_{41}V_{44}E_{32}D_{44}A_{61}I_{41}$ (the percentage occupancy of the consensus amino acid at each position is shown as subscript), and in FtsZ2 the consensus was $M_{66}Q_{34}D_{31}I_{43}V_{46}E_{23}E_{26}A_{80}L_{40}$. Inspection of amino acid substitutions in these motifs revealed that the charged or aliphatic character in each position was very strongly conserved.

After the initial N-terminal 9-amino-acid motif, FtsZ1 proteins then have an apparent spacer region (0-43 aa long, mean=21, SD=9, n=41) in the central portion of the N-tail that shows poor sequence conservation but is rich in charged amino acids and contains several proline and glycine residues, which are common in unstructured regions. This spacer is slightly longer on average than the corresponding charged region in FtsZ2

(0-18 aa long, mean=10, SD=5, n=35). At the C-terminal end of the N-tail, both archaeal FtsZ families show differing conserved motifs that are more conserved amongst FtsZ1 proteins ($D_{83}E_{41}E_{88}L_{88}E_{24}E_{27}V_{29}L_{63}E_{34}D_{22}L_{41}K_{34}$) than in FtsZ2 proteins ($D_{37}D_{20}D_{40}E_{49}F_{57}G_{69}$). This region forms an alpha helix (H0) in the previously determined crystal structure of *Methanocaldococcus jannaschii* FtsZ1 (Fig. S1d)[10]. Overall, the bacterial FtsZ N-tails appear to vary more than the archaeal families and lack the two archaeal sequence motifs.

*Core domains*

In the core domains, conserved differences between FtsZ1 and FtsZ2 were located in the vicinity of loops T4, T5, T6 and T7 in the N-terminal domain, which contribute to the GTP-binding interface located between subunits in FtsZ polymers (Fig. S1c-e)[10,11]. In some regions, bacterial FtsZ showed conserved matches specifically to FtsZ1 and in other regions to FtsZ2, whereas some other sequence features were apparently specific to FtsZ1 or FtsZ2 (*e.g.,* parts of the T7 loop region). Interestingly, the conserved T6-H6 amino acid triplet of FtsZ2 proteins ($DNN_{184-186}$ in HvFtsZ2) matched the equivalent region in eukaryotic tubulin and archaeal CetZ but differs from the triplet in the bacterial FtsZ and archaeal FtsZ1 families (Fig. S1c).

*C-terminal tails*

The C-tail also varies in average length and sequence between the families (Fig. S1b). An initial quitter charged variable region contains glycine and proline residues suggesting it may be another unstructured spacer (FtsZ1: 9-58 aa, mean=18, SD=10, n=41; FtsZ2: 10-78 aa, mean=37, SD=18, n=35), as seen in bacterial FtsZ (13-114 aa, mean=51, SD=24, n=25). A ~7 aa consensus motif is present at the end of the C-tail of archaeal FtsZ1 ($L_{27}G_{49}I_{76}D_{63}F_{51}V_{51}E_{24}$) and FtsZ2 ($L_{37}G_{71}I_{43}D_{69}V_{37}I_{49}R_{31}$) (Fig. S1c), which differ compared to the ~10 aa bacterial FtsZ motif (consensus DDLDIPAFLR) that plays a critical role in binding other division proteins[12].

**3. Different consequences of *ftsZ1* and *ftsZ2* mutations in stationary phase**

The differing functions of FtsZ1 and FtsZ2 were also apparent when cultures of the knock-out, complementation and overexpression strains were compared in mid-log and stationary phases. Wild-type (H98 + pTA962) cells became smaller in stationary phase compared to mid-log, and this trend was also observed in the strains containing FtsZ2, *i.e.,* Δ*ftsZ1*, Δ*ftsZ1* + *ftsZ1*, Δ*ftsZ1* + *ftsZ2*, and both overexpression strains (Fig. S12), in which almost wild-type cell sizes were attained in stationary phase. In contrast, the strains without FtsZ2 showed poor recovery from their cell division defects in stationary phase. These findings suggest that FtsZ2 confers a partial ability to divide and recover more normal cell sizes as the cell growth rate slows in stationary phase, whereas cells without FtsZ2 have a much stronger block to division that is maintained even as cells slow or stop growth in stationary phase.

**4. Different cell shapes of *H. volcanii* in various *ftsZ1* and *ftsZ2* mutant backgrounds**

There are several conditions known to cause *H. volcanii* cell shape changes, *i.e.*, transitions between elongated/rod cells and the plate morphotype, including during the development of motile rods (on 0.3% agar or at early stages of liquid culture), overexpression of cytoskeletal protein CetZ1, trace element starvation and other growth media conditions[13-15]. Cell elongation in at least some of these conditions was strongly dependent on the presence of a plasmid (based on the natural *H. volcanii* DS2 plasmid pHV2), modified to confer prototrophy in auxotrophic mutants[14]. It is currently unclear whether the shape effects are caused by plasmid replication/maintenance functions and/or its effect on auxotrophic/metabolic status.

A comparison of the cell morphology of *ftsZ* mutant strains in this study revealed several conditions that affected cell morphology. We observed that the Δ*ftsZ2* genotype is associated with highly filamentous cells and fewer giant plates in plasmid-containing (pTA962) strains compared to equivalent plasmid-free strains, which were mostly giant plates and debris (compare examples in Fig. 2b-c and 2g). Other strains based on the Δ*ftsZ2* background carrying a plasmid also showed filaments (Fig. 2h-i, S10a, S11c, S12c, S13b, S15b, S16a-b, S17, S19), except for the complementation strain Δ*ftsZ2* + FtsZ2 with 0.2 mM Trp or greater (Fig. S8). This suggests that the lack of FtsZ2 allows or promotes plasmid-dependent cell elongation (resulting in highly filamentous cells, owing to the division defect), which is likely to be related to one or more of the growth/media conditions listed above that cause plasmid-dependent rod formation in the wild type[13-15].

Interestingly, the Δ*ftsZ1* strain did not form elongated or filamentous cells with or without pTA962 (Fig 2b and 2g) and neither did the other strains based on the Δ*ftsZ1* background carrying a plasmid (Fig. S10a, S12b, S13a, S15a, S16a-b, S19). Instead, various backgrounds showed rod/elongated cells when FtsZ1 was overproduced, some of which displayed a noticeable taper or tubular narrowing at cell poles (Fig. 2h, 3, S8a-b, S9a, S11, S12a). These results suggest that the cell elongation seen in Δ*ftsZ2* + plasmid strains might require FtsZ1, and that the normal function of FtsZ1 influences cell elongation. This is apparently via influencing or recruiting the cell envelope machinery (Fig. 6), but how FtsZ1 does this and influences the structure of cell poles, are unknown and should be of interest in future.

In the double knock-out (Δ*ftsZ1* Δ*ftsZ2*), filamentous cells (~31% of the total detected) were observed even in the absence of pTA962 (Fig. 2, S7c), highlighting the multi-factorial nature of cell shape determination in *H. volcanii*. These cells also displayed tubulation-fission events at cell poles (Fig. 2d-e, Video S8). It is unknown whether the formation of these cell structures is due to the specific combination of Δ*ftsZ1* Δ*ftsZ2* or potentially related to additional mutations present (Table S5). While the division (cell size) defect of this strain was corrected by expression of the two *ftsZ* genes on a plasmid (Fig. S9), it remains to be seen whether this also fully reversed the apparent gain-of-function filamentation and tubulation-fission phenotypes; some cells exhibited obvious polar tubules (Fig. S9a, 2 mM Trp), which could be attributed to the overproduced FtsZ1 and/or additional mutations. The results overall emphasize the complex nature of cell shape determination and highlight the importance of careful interpretation of results of cell division or other experiments in which cell shape is also affected, directly or indirectly, through mutations.

**5. FP-tagging of the T7-mutants suppresses their dominant-inhibitory phenotypes yet reveals co-localization with the wild-type FtsZ proteins and independent filaments of FtsZ1**

Based on the results shown in Fig. 5, we tagged the T7 mutants in order to simultaneously visualize both the T7-mutant of one FtsZ and the wild-type copy of the other. The tagged T7 mutants failed to complement their respective knockout strains, as expected, and localized weakly as foci or short filaments (Fig. S19a-b). Surprisingly, we found that the tags almost completely suppressed the strong dominant-inhibitory effects of the T7 mutations in the wild-type background; the cell size had returned almost to normal, yet the proteins still localized (Fig. S19c-d). Indeed, FtsZ2.D231A-GFP caused a substantially milder division defect than FtsZ2-GFP (1 mM Trp; Fig. S15d, S19d). Similarly, FtsZ1.D250A-mCh caused minimal division defects and showed midcell localization, and some aberrant localizations away from midcell (Fig. 5c right). At 1 mM Trp, there were more noticeable cell deformations and protrusions, and small extracellular fluorescent particles (Fig. S19c right). Generally, these results indicated that FtsZ1.D250A-mCh was more independent of midcell than the more subservient FtsZ2.D231A-GFP. The tagged T7-mutants therefore appear to form mixed filaments with the wild-type FtsZ proteins that retain a capacity to localize without substantially disrupting the activity of the division machinery.

The two alternate combinations of the tagged T7-mutant and wild-type proteins were then co-expressed. In Δ*ftsZ2*, the presence of GFP on FtsZ2.D231A-GFP caused additional disruption to the apparent condensation FtsZ1-mCh helicoidal structures (compare Fig. S19e left to Fig. 5a-b). The FtsZ2.D231A-GFP formed foci that co-localized with patches of FtsZ1-mCh at irregular intervals in filaments (Fig. S19e). Similar results were seen in the Δ*ftsZ1* Δ*ftsZ2* background, though with further irregularities in FtsZ1-mCh structures (Fig. S19i middle). In the Δ*ftsZ1* background, FtsZ1.D250A-mCh and FtsZ2-GFP co-localized in patches around the edges of giant plates (S19f; compare to Fig. 5b right). In addition, there were independent FtsZ1.D250A-mCh edge-patches. In the Δ*ftsZ1* Δ*ftsZ2* background, FtsZ1.D250A-mCh was similarly edge-associated and thus minimally affected by the additional loss of FtsZ2 (Fig. S19i right), but FtsZ2-GFP was diffuse with a few foci elsewhere, consistent with the requirement for the wild-type FtsZ proteins for normal FtsZ2-GFP localization (Fig. S15). These results illustrate that FtsZ2 is not required for FtsZ1 assembly but has a moderate influence on the structures or condensation of FtsZ1 assemblies, whereas FtsZ2 assembly and structures strongly follow those of FtsZ1.

In the wild-type background, FtsZ1-mCh and FtsZ2.D231A-GFP co-localized at midcell and the cells appeared of normal size and shape (Fig. S19g), consistent with the minimally disruptive behaviour of the two proteins individually (Fig. 4a, S19d). Only at 1 mM Trp did we see some mild aberrant localization (Fig. S19g left inset), although cell size and shape were still not substantially affected (Fig. S19g right). On the other hand, FtsZ1.D250A-mCh with FtsZ2-GFP generated a moderate cell division defect, as expected with FtsZ2-

GFP (Fig. S19h; severe at 1 mM Trp, inset). The FtsZ1.D250A-mCh and FtsZ2-GFP displayed remarkable co-localization with as helical or extended filaments, but notably contained numerous independent filaments containing FtsZ1.D250A-mCh. Comparison of Fig. 5c, S19d and S19h suggests that the filaments containing FtsZ2-GFP are caused by their association with long aberrant filaments containing FtsZ1.D250A. Taken together with a comparison of Fig. 5a, 5b and S19f (left panels), the combined data strongly support the view that: (1) FtsZ1 localization is largely independent of FtsZ2, (2) FtsZ2 assembly, positioning and stabilization are heavily dependent on the presence, localization and correct functioning of FtsZ1, and (3) FtsZ1-ring condensation (via a helicoid intermediate) is in turn promoted by feedback from the presence and correct functioning of FtsZ2.

#### Supplementary Figures

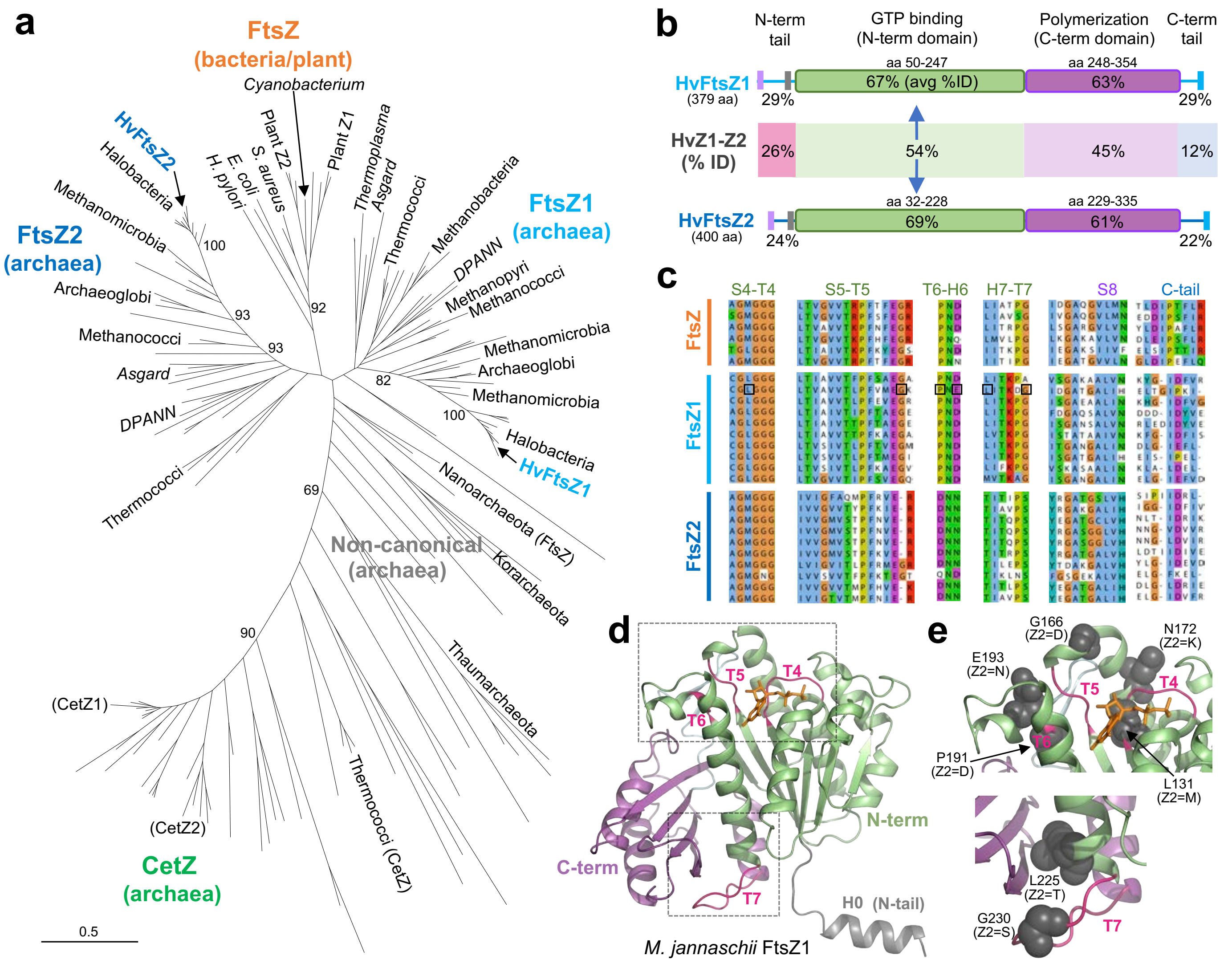

**Figure S1. Molecular phylogeny and comparison of archaeal FtsZ1 and FtsZ2 families.**

**(a)** Phylogenetic tree of the identified archaeal tubulin superfamily proteins, and the bacterial/plant sequences used to identify them. Bootstrap support is shown for selected branches (%). **(b)** Domain organization and percent sequence identities for FtsZ1 and FtsZ2. The percentages over the domains (green and purple boxes) indicate the average sequence identity in that region for each *H. volcanii* FtsZ compared to all of the other members of the same family that were identified across the Archaea domain. The region between FtsZ1 and FtsZ2 represents the percent identity in the region between the *H. volcanii* FtsZ1 and FtsZ2 (%ID). The approximate location of conserved sequence motifs within the tail regions are indicated by vertical bars, coloured to indicate similarities between the two. **(c)** Aligned sequence regions containing conserved differences between the bacterial/plant FtsZ and the archaeal FtsZ1 and FtsZ2 families, labelled with the secondary structural elements [11]. Boxed residues indicate conserved sites that are displayed in panel **(e)**. **(d)** Crystal structure of FtsZ1 from *Methanocaldococcus jannaschii* (PDB: 1FSZ)[10], with selected loops (T4-T7) involved in nucleotide binding and hydrolysis shown in pink. GDP is shown in orange, and the main domains are coloured as in panel **(b)**. Boxed regions are expanded in panel (**e**), which displays some conserved residues that characteristically differ between the FtsZ1 and FtsZ2 families (grey space-filling models, with FtsZ2 consensus residues in parentheses) and cluster around the nucleotide-dependent polymerization surfaces.

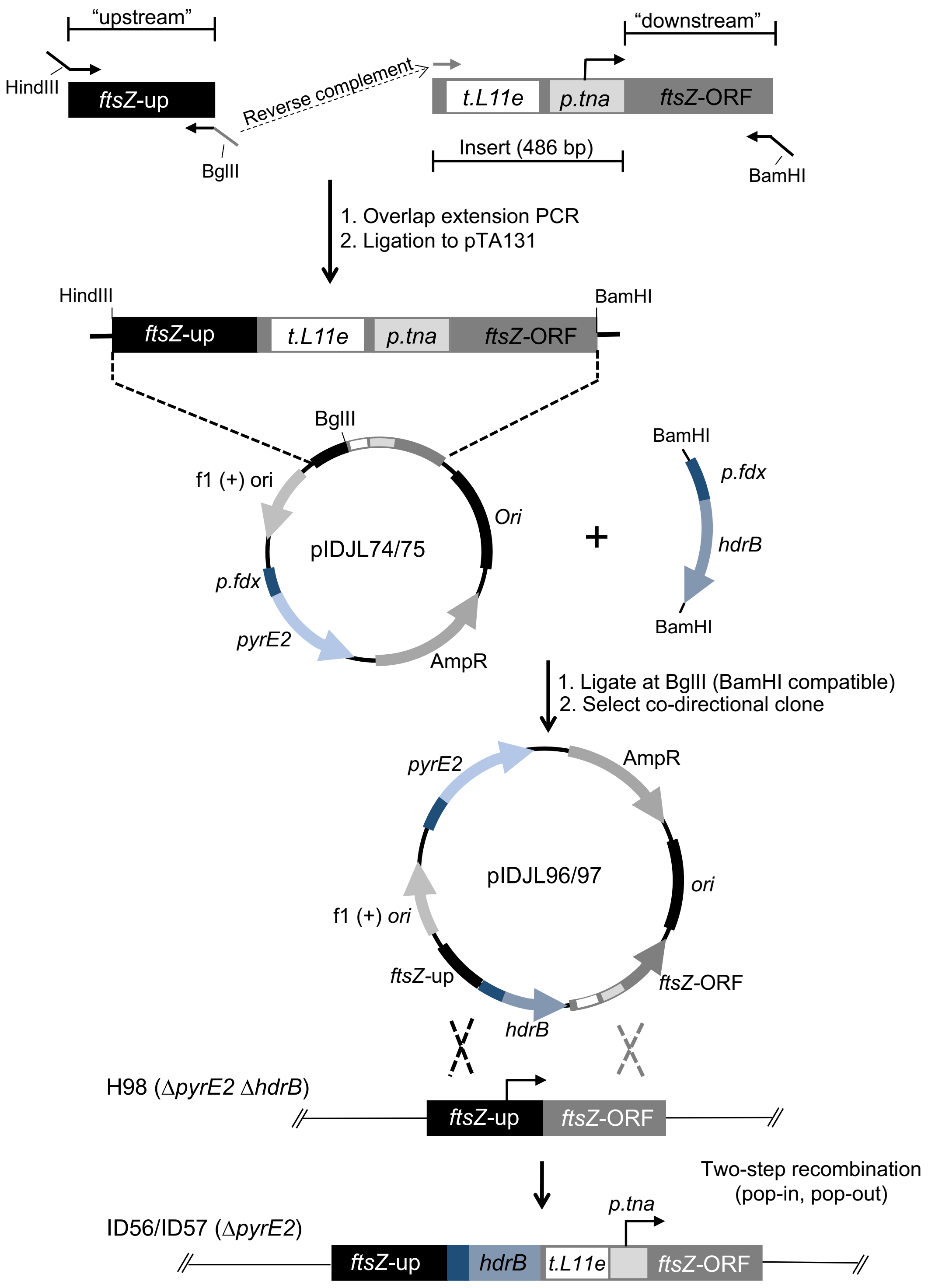

**Figure S2. Construction of *ftsZ*-depletion strains.**

The chromosomal *ftsZ* genes were individually placed under control of the tryptophan-regulated promoter, *p.tna*. Flanks for homologous recombination (upstream and downstream) were amplified and spliced (top), giving a product that included the t.L11e transcription terminator and *p.tna* promoter configured to drive *ftsZ* expression. This was cloned into pTA131 (at HindIII and BamHI), to give pIDJL74/75, and the cloned sequences were confirmed. The *hdrB* marker from pTA1185 was then inserted at the BglII site of pIDJL74/75, and a clone with *hdrB* co-oriented with *ftsZ* in each case was selected and used for transformation of *H. volcanii* (H98), applying the two-step procedure for genomic DNA modification that yields the genomic structure shown at the bottom, allowing Trp-control of *ftsZ1* (strain ID56) or *ftsZ2* (ID57) expression. Note that in *H. volcanii* ID56 the *p.tna* cassette replaces the predicted *ftsZ1* promoter. In *H. volcanii* ID57, the *p.tna* cassette is instead inserted between *ftsZ2* and the upstream gene (HVO_0582) (which show a 2 bp gap between ORFs), resulting in no loss of genomic DNA.

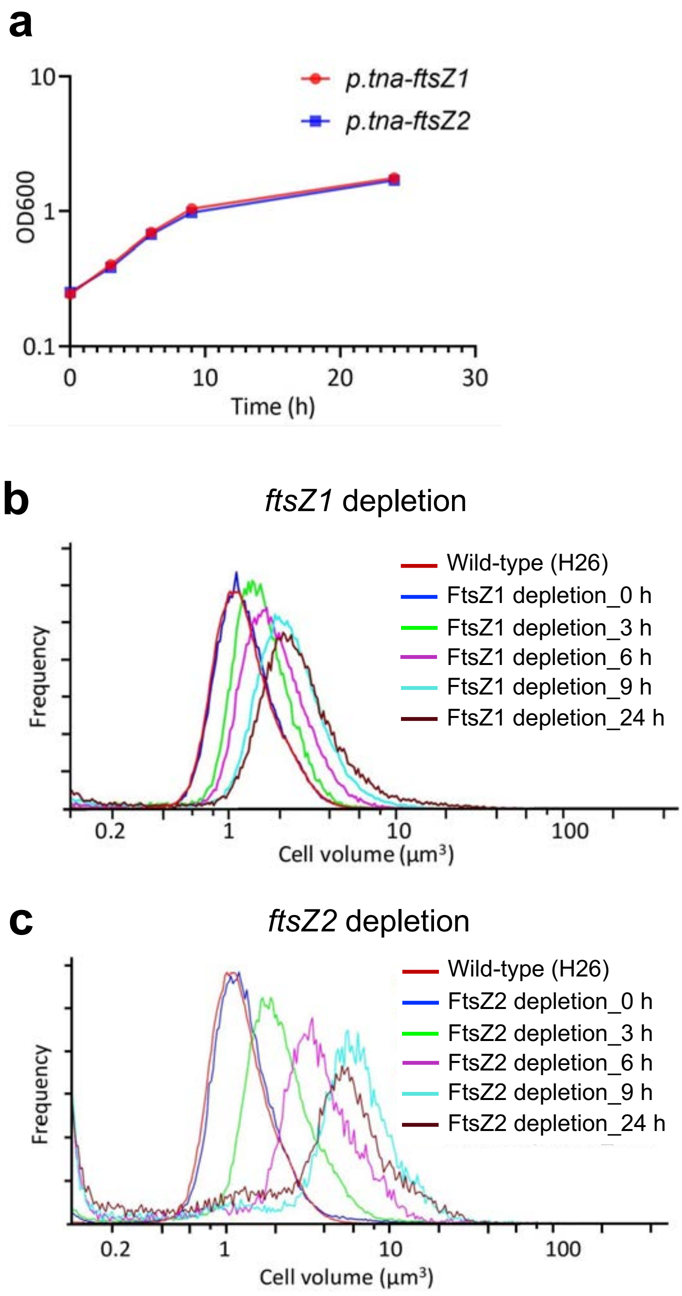

**Figure S3. Growth curves and cell size distributions during depletion of FtsZ1 and FtsZ2.**

Each *ftsZ* gene was under the control of the *p.tna* inducible promoter in *H. volcanii* ID56 (*p.tna-ftsZ1*) and ID57 (*p.tna-ftsZ2*), and samples were withdrawn at the indicated timepoints over 24 h after removal of Trp from mid-log cultures. **(a)** Growth curves ($OD_{600}$) and Coulter cell-volume distributions of strains *p.tna-ftsZ1* **(b)** and *p.tna-ftsZ2* **(c)**. Samples were from the same cultures as shown in Fig. 1 (main article). The same dataset for the wild-type H26 control is shown in both graphs as a reference.

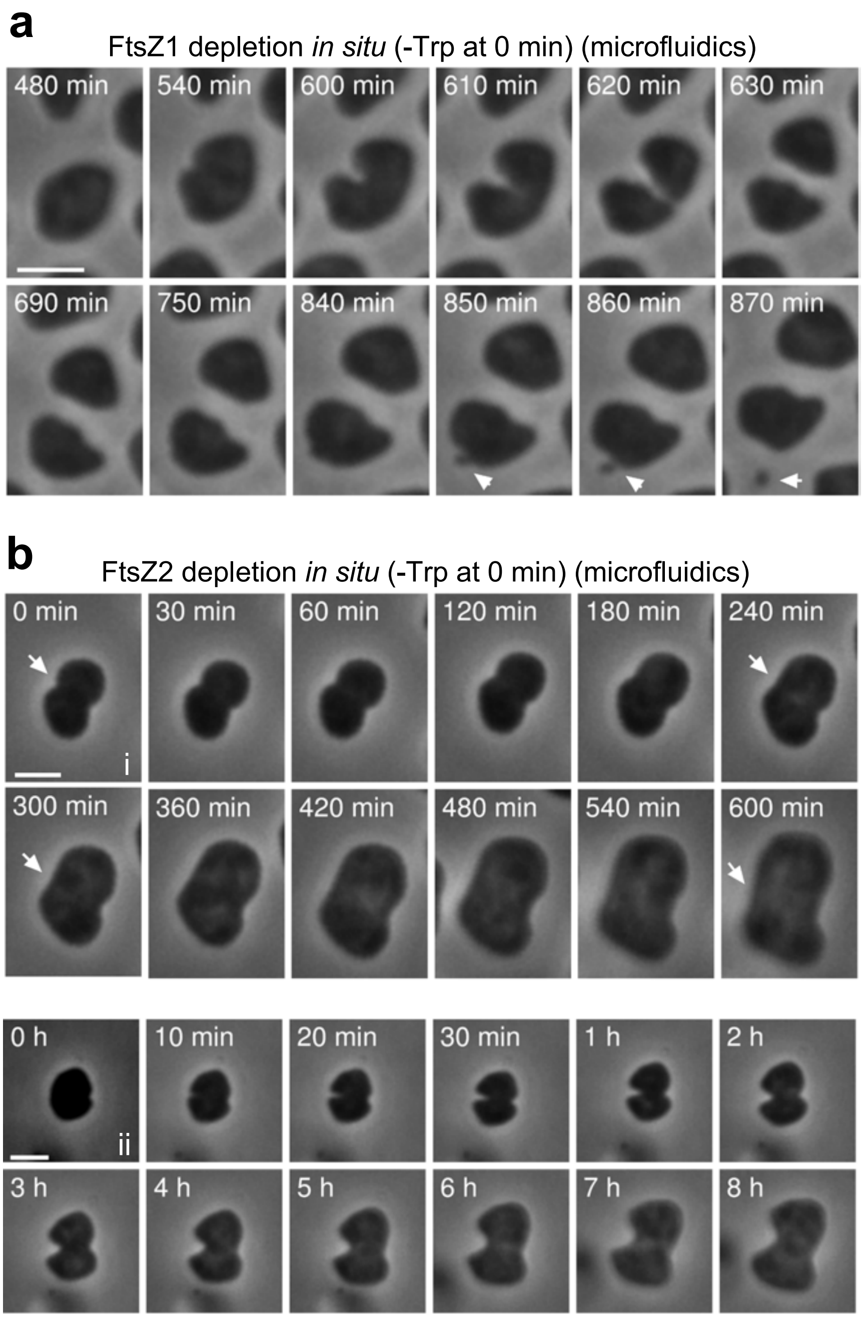

**Figure S4. Partial division phenotypes during depletion of FtsZ1 or FtsZ2.**

**(a)** *H. volcanii* ID56 (*p.tna-ftsZ1*) was cultured in Hv-Cab + 2 mM Trp, and then loaded into a microfluidics platform and cultured with a flow of Hv-Cab (without Trp) over 15 h (0.5 p.s.i) to deplete FtsZ1. Shown is one cell that was identified to divide (unilaterally), even after ~9 h of depletion, and then one cell exhibited a budding-like process (arrows). Scale bar, 2 µm. **(b)** *H. volcanii* ID57 (*p.tna-ftsZ2*) was pre-cultured in Hv-Cab + 2 mM Trp, and then loaded into a microfluidics platform and cultured with a flow of Hv-Cab + 2 mM Trp for 3 h, followed by Hv-Cab (no Trp) for 10 h (2 p.s.i) to deplete FtsZ2. The zero timepoint represents the start of medium flow without Trp. During the early stage of depletion of FtsZ2, partial constrictions were sometimes observed, as seen in these two examples, but these never completed division and the constriction eventually reversed over several hours (see arrows). Cells, however, retained some apparent 'memory' of the initial constriction often manifesting in a somewhat bilobed shape. Scale bar, 2 µm.

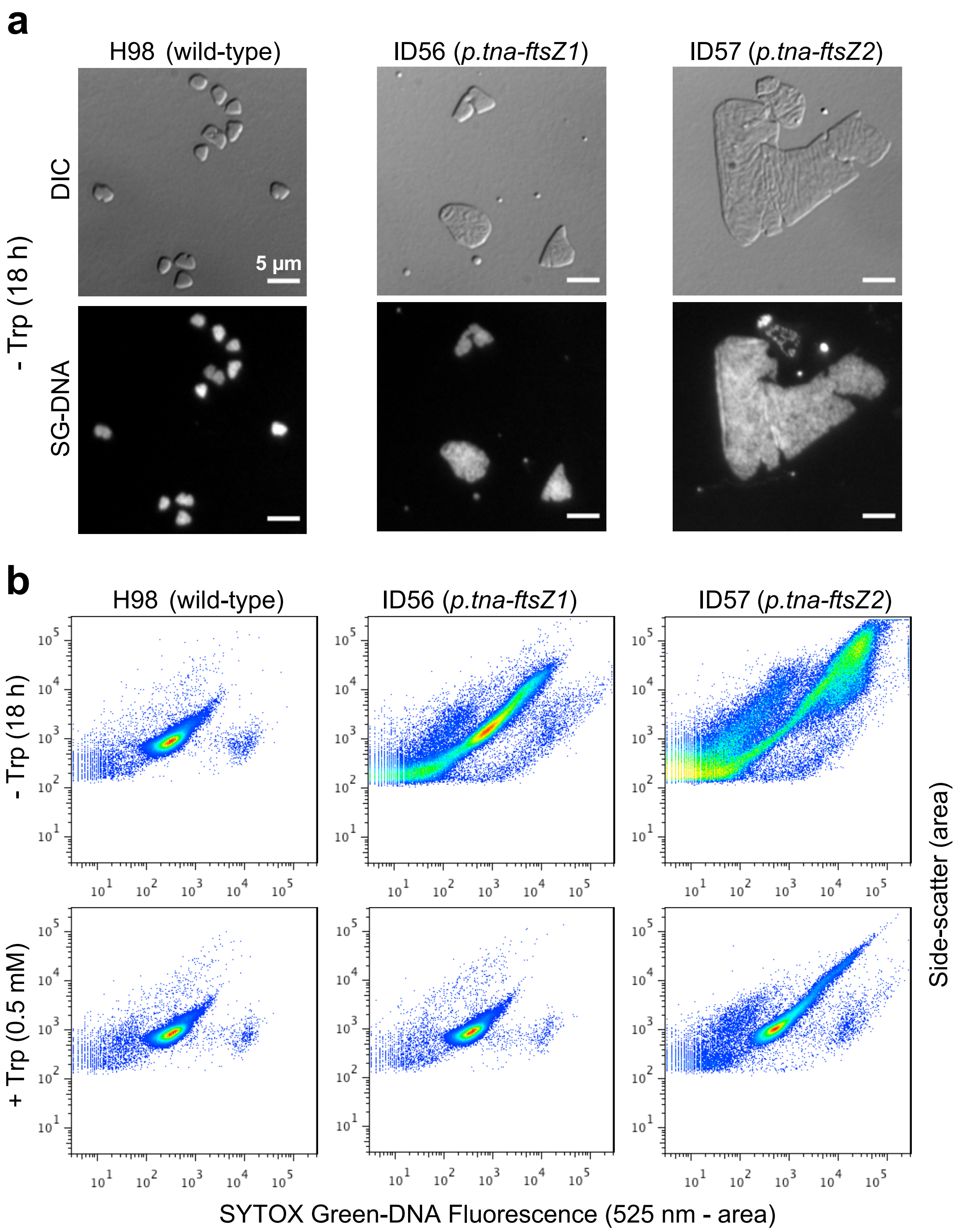

**Figure S5. Cellular DNA content during depletion of FtsZ1 and FtsZ2.**

**(a)** SYTOX Green (SG) DNA staining of cells sampled from cultures 18 h after resuspension of mid-log cells in media without Trp. Stained cells were placed on an agarose pad and visualized by differential-interference contrast (DIC) and fluorescence microscopy (lower panels). Scale bars are 5 µm. **(b)** Flow cytometry analyses of cells sampled as per panel (a) (upper three panels), displaying side-scatter (as a proxy for cell size) versus SYTOX Green (SG)-DNA fluorescence. The lower three panels represent cultures treated in the same way, except 0.5 mM Trp was included in the medium. The individual datapoints represent the area under the curve of each event detected; events were detected by a threshold of the side-scatter signal. After 18 h of *ftsZ1* or *ftsZ2* depletion, many very large cells with correspondingly high DNA content were observed, consistent with the images shown in panel (a). This indicates that DNA synthesis continues in proportion to the increase in cell volume during inhibition of cell division caused by depletion of FtsZ1 or FtsZ2.

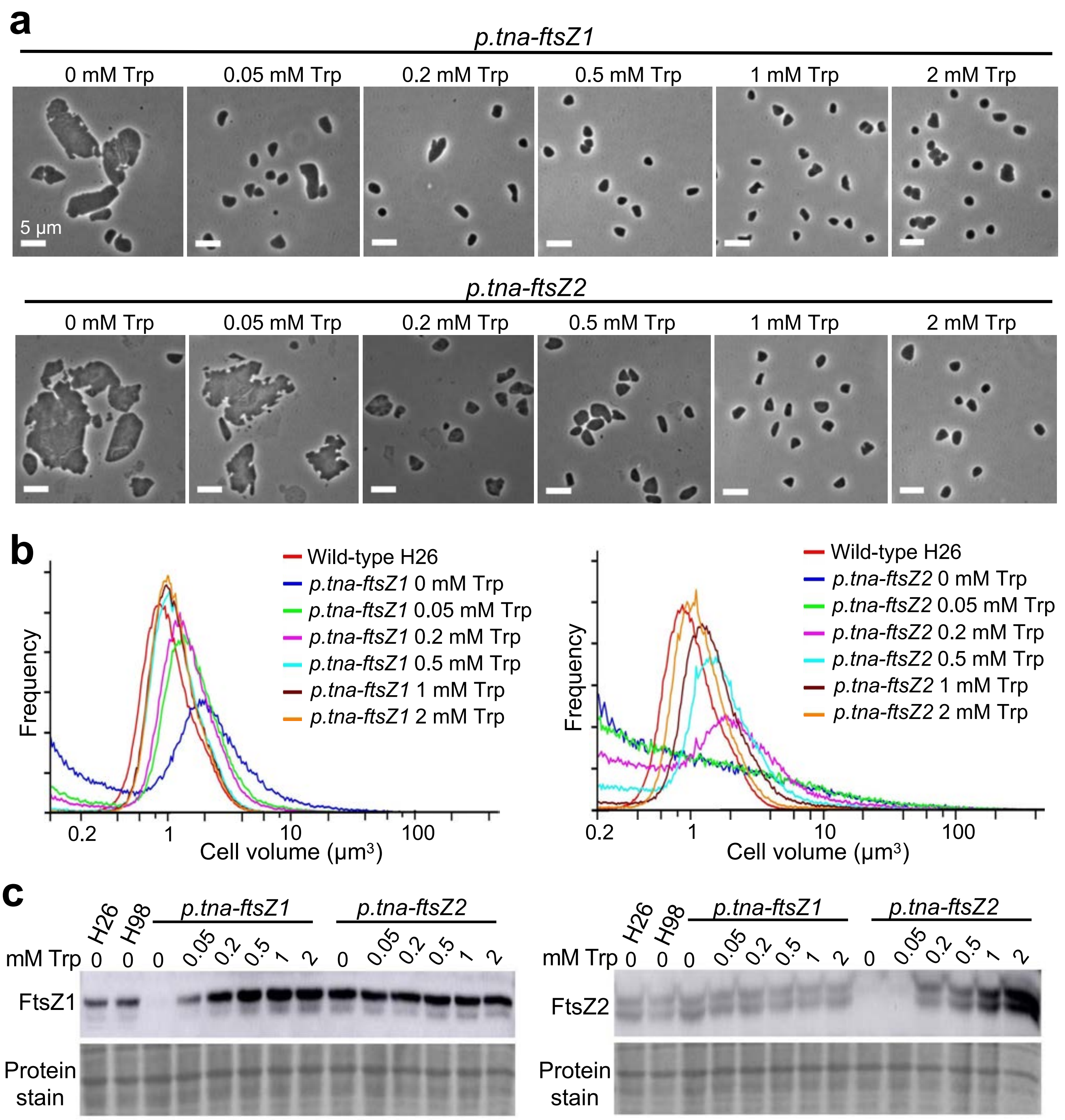

**Figure S6. Inducer (Trp) concentration-dependence of cell size in *ftsZ*-depletion strains.**

**(a)** Phase-contrast microscopy of *H. volcanii* ID56 (*p.tna-ftsZ1)* and ID57 (*p.tna-ftsZ2*) sampled during steady mid-log growth with the indicated concentrations of Trp. **(b)** Coulter cell-volume distributions of *p.tna-ftsZ1* and *p.tna-ftsZ2* cultures, sampled as in panel (a). The same dataset for the wild-type H26 control is shown in both graphs as a reference. **(c)** Western blots of *p.tna-ftsZ1* and *p.tna-ftsZ2* strains, sampled as described above, probed with antibodies raised against synthetic peptides based on unique sequences within FtsZ1 and FtsZ2. The levels of FtsZ1 or FtsZ2 in their corresponding *p.tna-ftsZ1* and *p.tna-ftsZ2* strains increase in response to increasing concentrations of Trp in the medium. The recovery of cell sizes and normal protein levels occurred at lower concentrations of Trp for *p.tna-ftsZ1* compared to *p.tna-ftsZ2*; cells appeared the same as wild-type size and shape in the presence of at least 0.5 mM Trp for *p.tna-ftsZ1*, and 2 mM Trp for *p.tna-ftsZ2*.

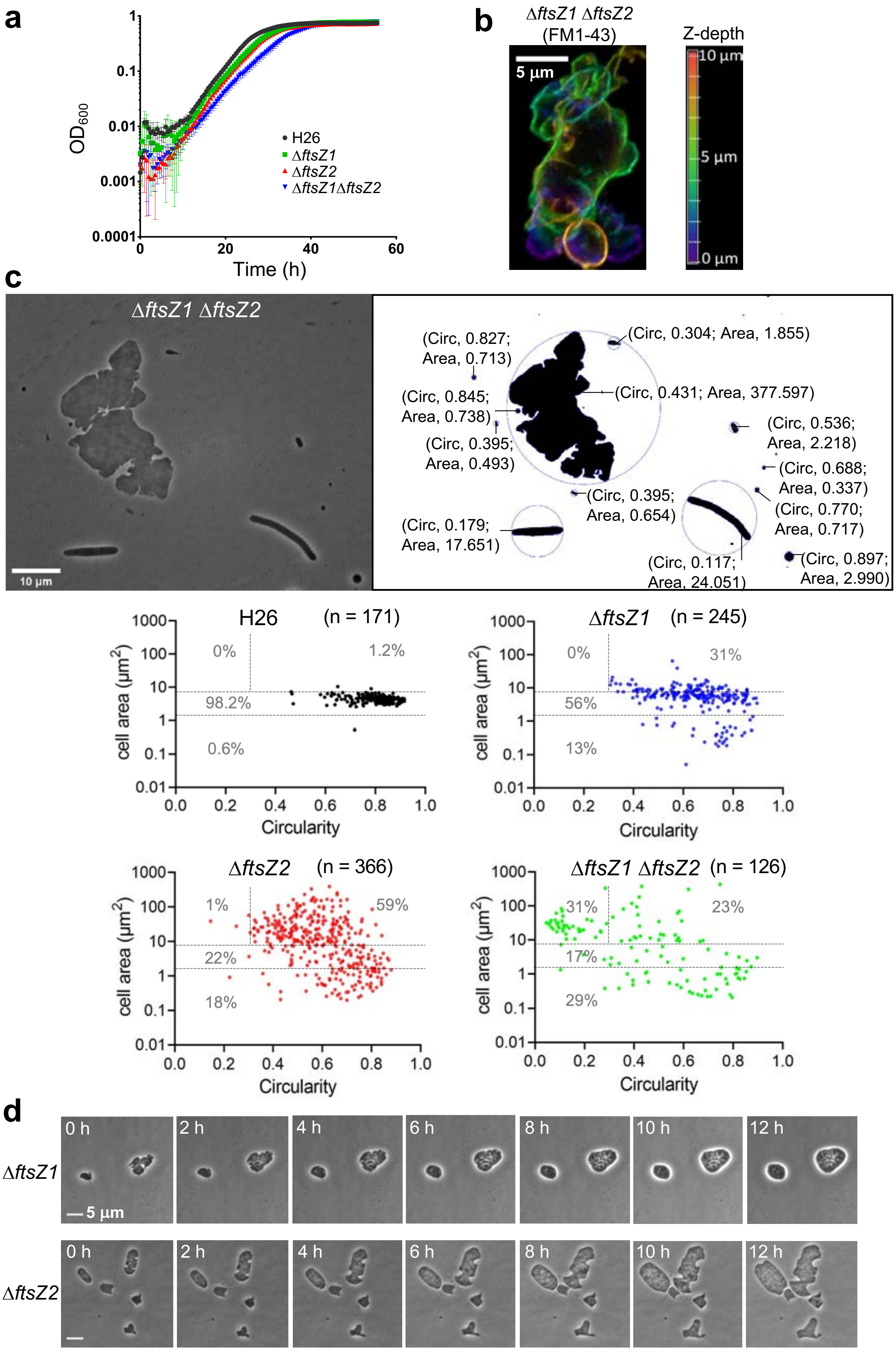

**Figure S7. Phenotypes of Δ*ftsZ* strains.**

**(a)** Microtiter plate monitoring of growth ($OD_{600}$) of *H. volcanii* wild type (H26), Δ*ftsZ1* (ID76), Δ*ftsZ2* (ID77), and Δ*ftsZ1* Δ*ftsZ2* (ID112). **(b)** Confocal 3D images (xy-view) with a false-colour z-depth cue of Δ*ftsZ1* Δ*ftsZ2* stained with FM1-43 membrane dye. **(c)** An example image of Δ*ftsZ1* Δ*ftsZ2* (ID112) (left) and the corresponding threshold image for cell size and shape analysis. Cell circularity was calculated here as the fractional area of the minimal circle that completely encircles the cell. Area values are in $\mu m^2$. The cell area versus circularity scatter plots show the percentage of cells/particles (datapoints) classified as filaments (cell area > 7.5 $\mu m^2$, Circularity < 0.3), giant plates (Cell area > 7.5 $\mu m^2$, Circularity > 0.3), wild-type-like cells (any shape, with cell area between 1.5 $\mu m^2$ and 7.5 $\mu m^2$), or cellular debris (< 1.5 $\mu m^2$) for the indicated strains (sampled at $OD_{600}$ = 0.2). **(d)** Live cell *in situ* time-lapse microscopy image series (see supplementary Video S5-S7).

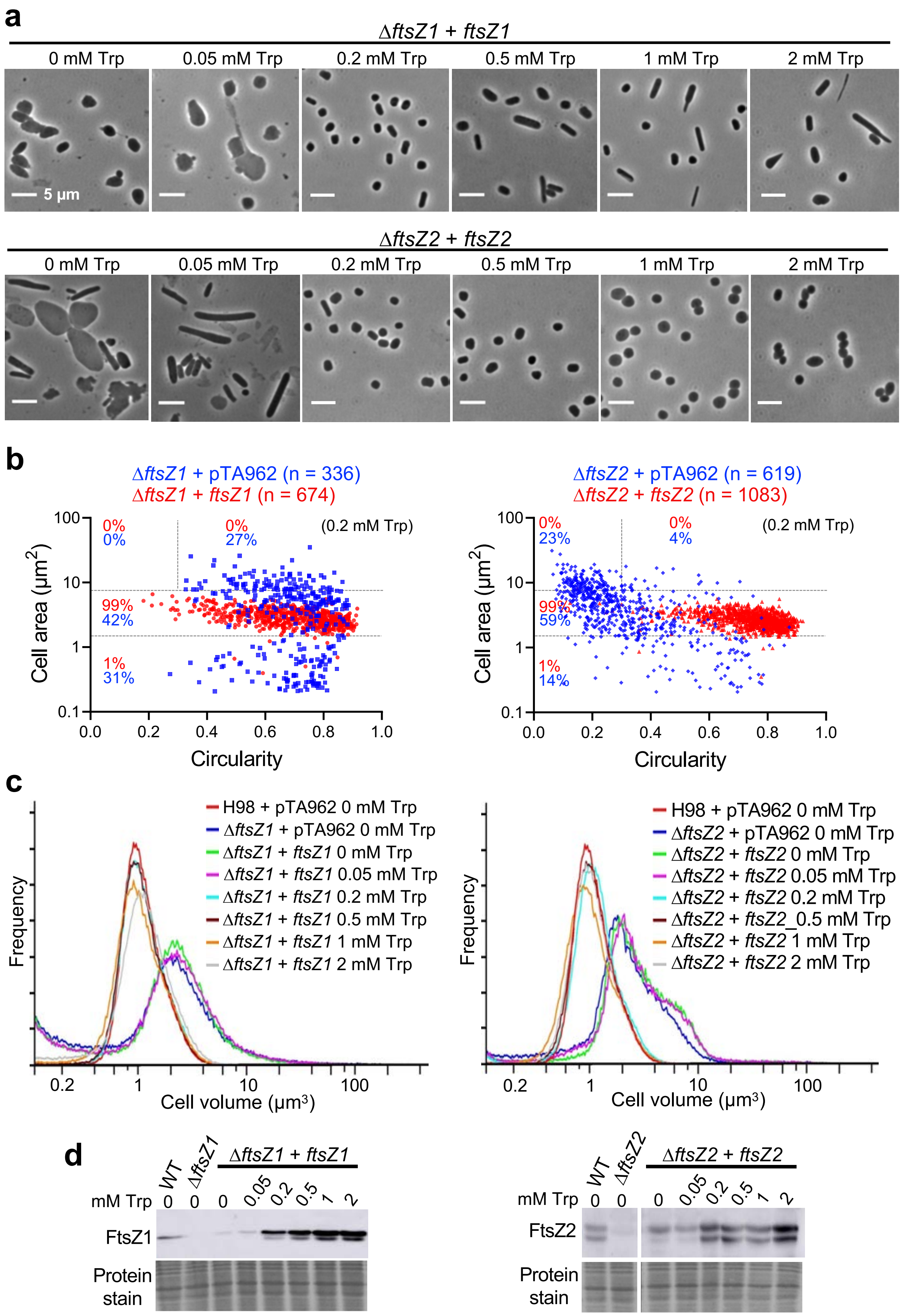

**Figure S8. Complementation of single Δ*ftsZ* strains.**

**(a)** Phase-contrast images of *H. volcanii* FtsZ1 self-complementation strain, ID86 (Δ*ftsZ1* + pTA962-*ftsZ1*), and FtsZ2 self-complementation strain, ID92 (Δ*ftsZ2* + pTA962-*ftsZ2*), sampled during mid-log growth with the indicated concentrations of Trp. **(b)** Cell shape quantification of the indicated midlog strains (cultured with 0.2 mM Trp, sampled at $OD_{600}$ = 0.3). **(c)** Coulter cell volume analyses of midlog cultures of the *ftsZ1* and *ftsZ2* complementation strains. The same dataset for the wild-type control is shown in both graphs as a reference. **(d)** Western blot analyses of FtsZ1 and FtsZ2 levels.

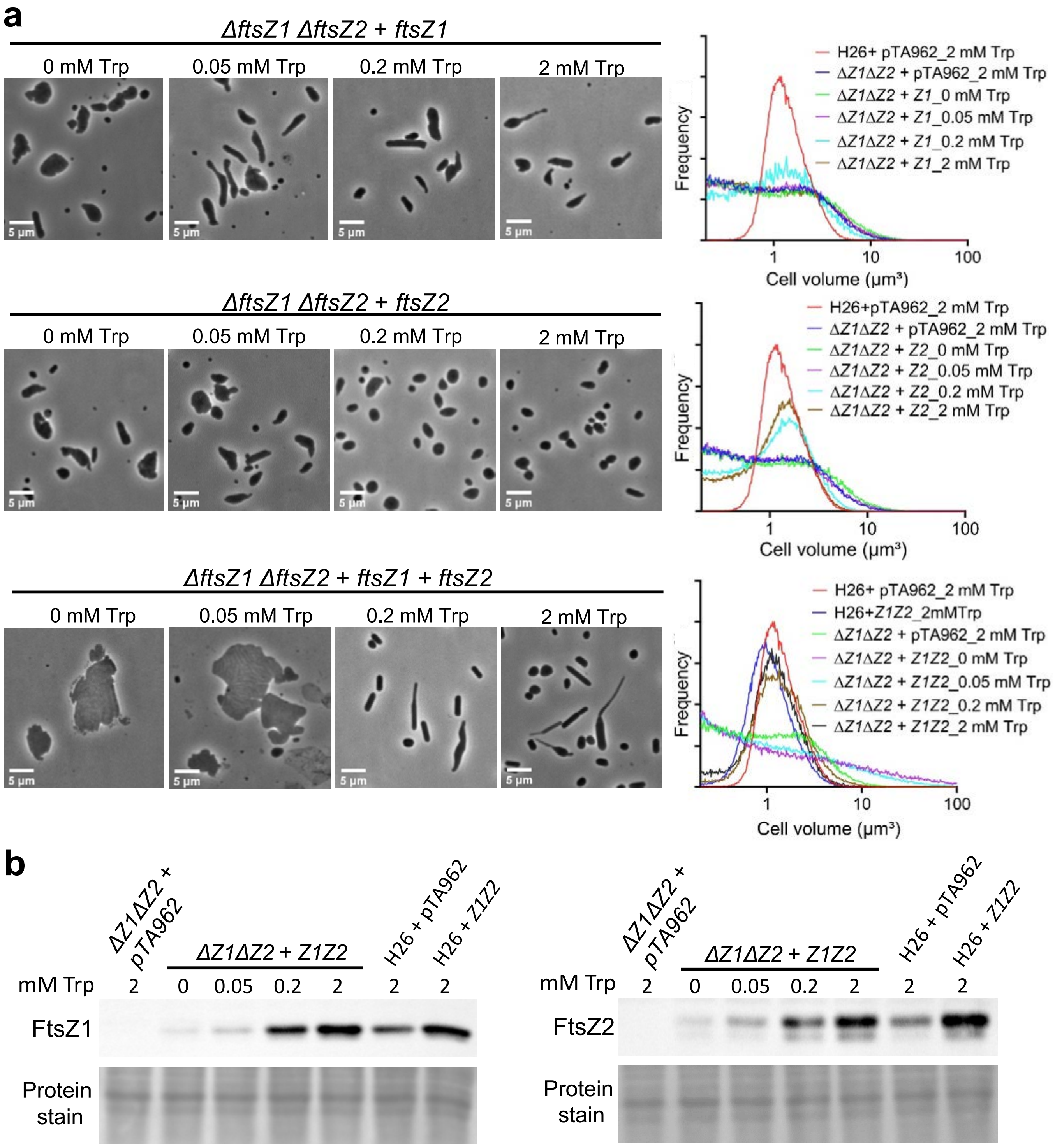

**Figure S9. Complementation of Δ*ftsZ1* Δ*ftsZ2*.**

**(a)** Phase-contrast images (left) and Coulter cytometry (right) of strains based on *H. volcanii* ID112 (Δ*ftsZ1* Δ*ftsZ2*), plus pTA962-based plasmids expressing the indicated *ftsZ* genes, sampled during mid-log growth with the indicated concentrations of Trp. The same dataset for the wild-type control (H26 + pTA962) is shown in all graphs as a reference. **(b)** Corresponding western blot analyses of FtsZ1 and FtsZ2 protein levels in total cell extracts of the indicated strains.

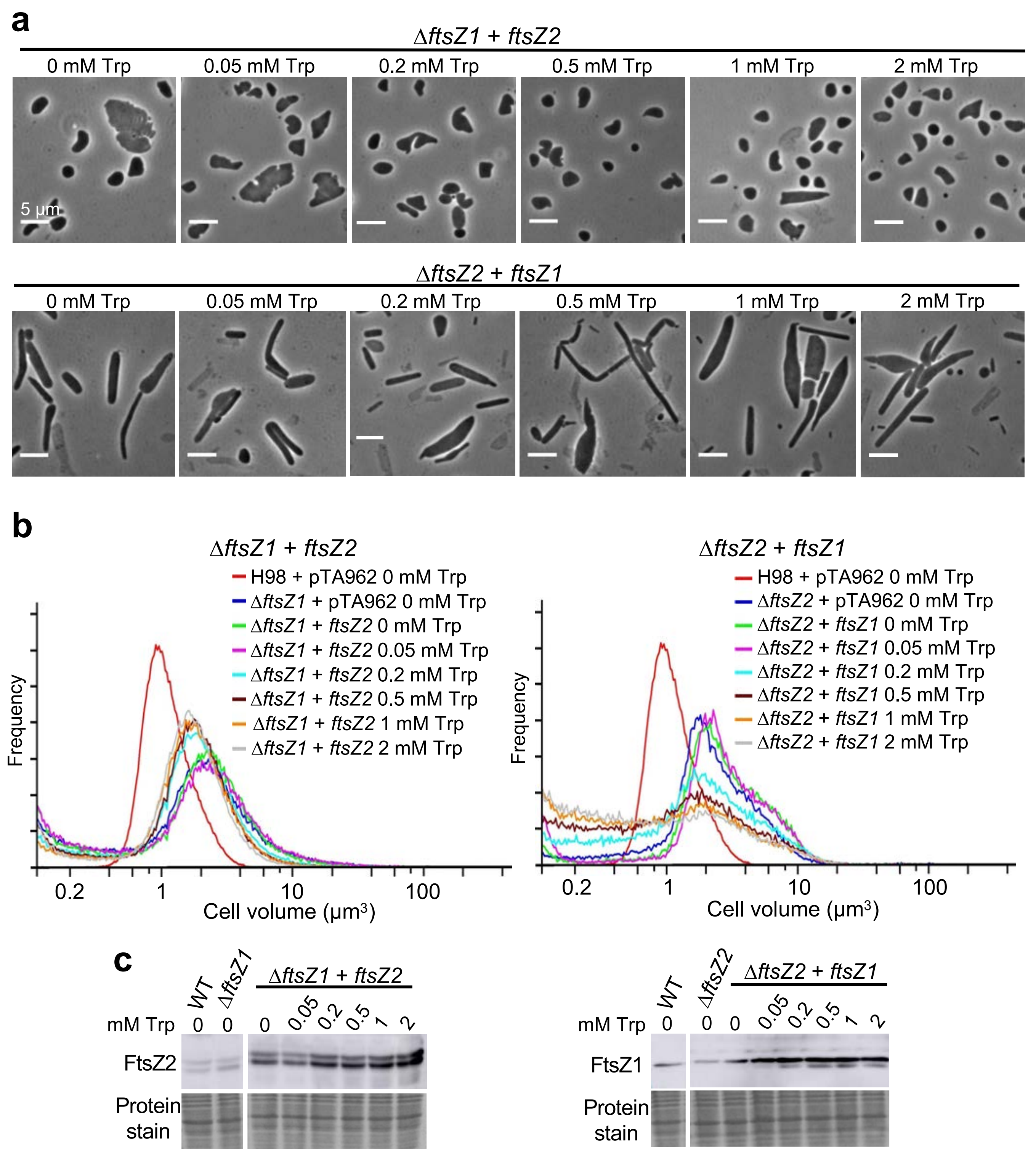

**Figure S10. FtsZ overproduction fails to properly complement strains carrying a knockout of the other FtsZ.**

**(a)** Phase-contrast images of midlog samples of the cross-complementation strains, Δ*ftsZ1* + pTA962-*ftsZ2*) and Δ*ftsZ2* + pTA962-*ftsZ1*, in the indicated concentrations of Trp. **(b)** Coulter cell volume analysis of the cross-complementation strains. The same dataset for the wild-type control is shown in all graphs as a reference. Expression of *ftsZ2* appears to partially complement the *ftsZ1* knockout defect at ≥ 0.2 mM Trp, whereas *ftsZ1* did not show any noticeable rescue of the *ftsZ2* knockout in all the tested concentrations of Trp. The *ftsZ2* knockout with *ftsZ1* overexpression (right) shows severe cell size defect at all tested concentrations of Trp, and elevated cell debris (< 0.5 µm$^3$) at Trp concentrations ≥ 0.2 mM, suggesting that the defect becomes more severe with increasing FtsZ1 levels in this strain. **(c)** Corresponding western blot analyses of the cross-complementation strains.

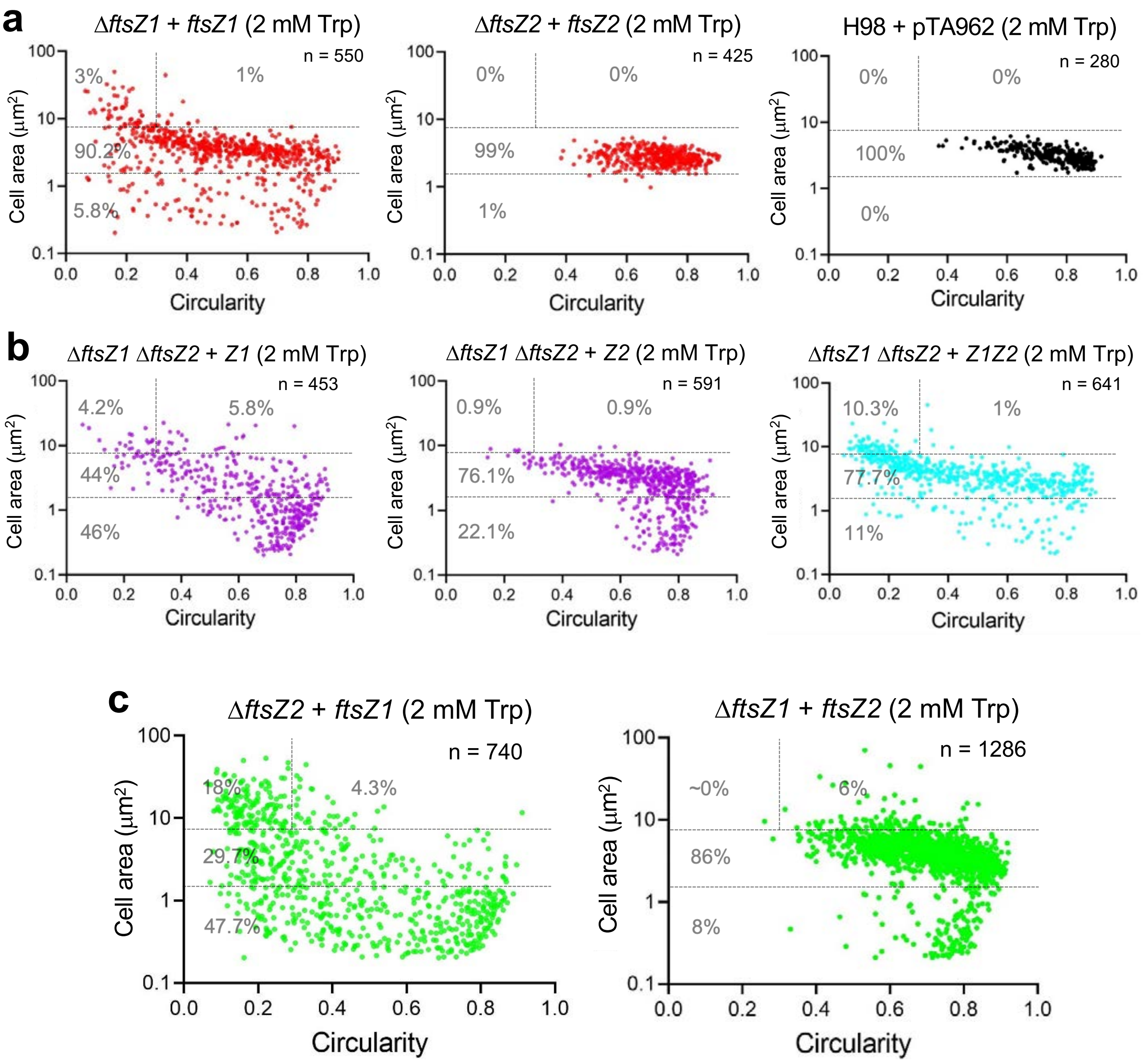

**Figure S11. Cell morphology analysis of *ftsZ1/2* overexpression in various *ftsZ* mutant backgrounds.**

Cell shape quantification scatterplots were generated from analysis of phase-contrast images of the indicated midlog strains, cultured with high-level induction of *ftsZ* expression (2 mM Trp). **(a)** *ftsZ* complementation strains and wild-type control (H98 + pTA962). **(b)** double-*ftsZ* knockout complementation strains. **(c)** Cross-complementation strains. Overexpression of *ftsZ1* increases cell elongation (decreased circularity) in all backgrounds, whereas *ftsZ2* overexpression reduces cell size (area). Reference knockout strains are shown in Fig. S8b.

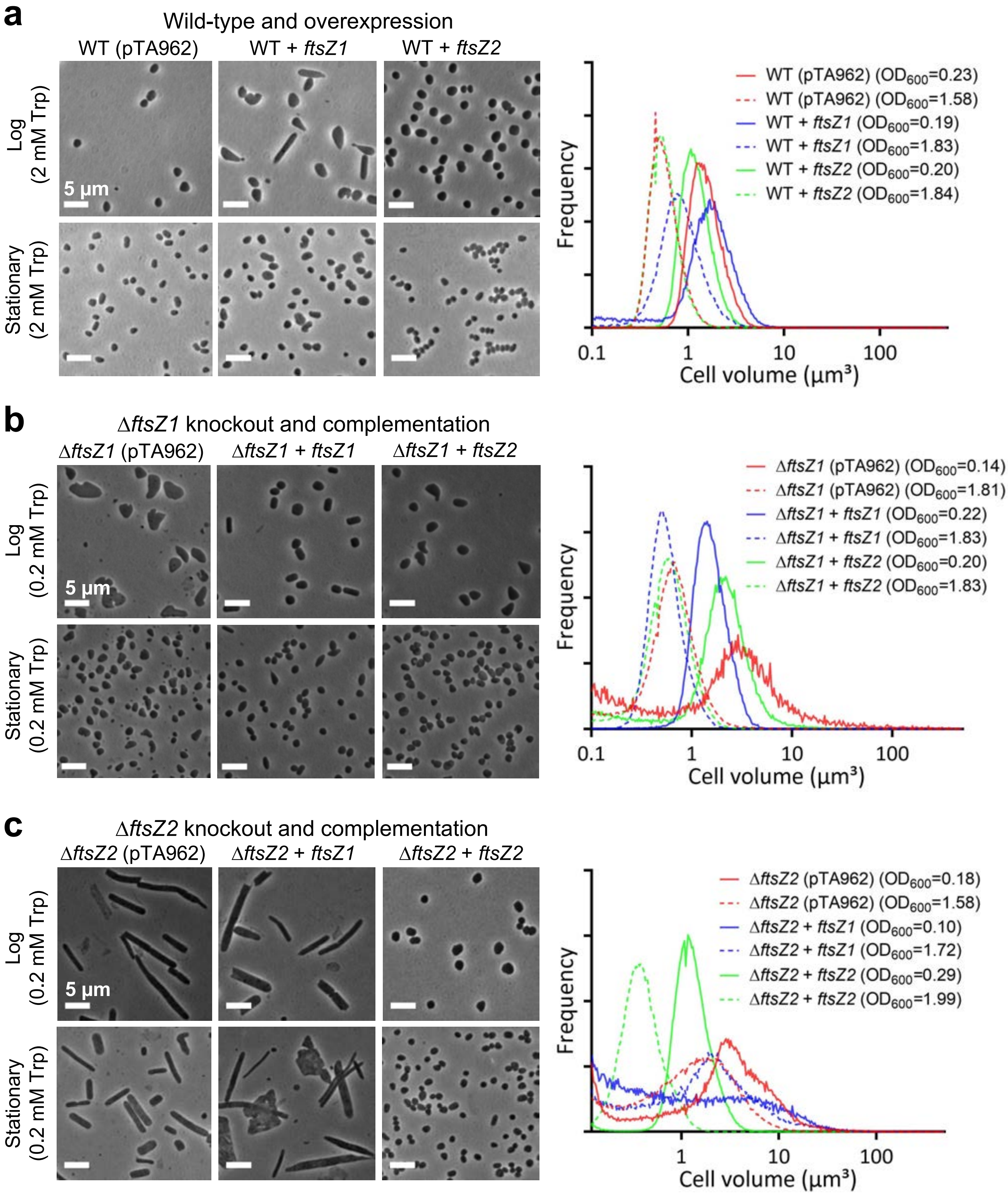

**Figure S12. Comparison of *ftsZ* mutant cellular phenotypes in log and stationary phases.**

Phase-contrast images (left) and Coulter cytometry distributions (right) of the wild-type and overexpression strains **(a)** and the indicated *ftsZ* knockout and complementation strains **(b-c)**, all grown in Hv-Cab with the indicated concentration of Trp and sampled at mid-log and stationary phases. The $OD_{600}$ of the cultures at sampling is shown in the graphs, where mid-log samples are shown in solid colour lines and stationary phase samples in dashed lines. Compared to mid-log cells, all the strains except the strains without a copy of *ftsZ2*, tended towards the wild-type size (smaller) and regular plate morphology in stationary phase. The Δ*ftsZ2* strains were somewhat smaller in stationary phase, but maintained greatly enlarged giant plate and elongated cells, suggesting a poor recovery as cell growth slows in stationary phase. All scale bars are 5 µm.

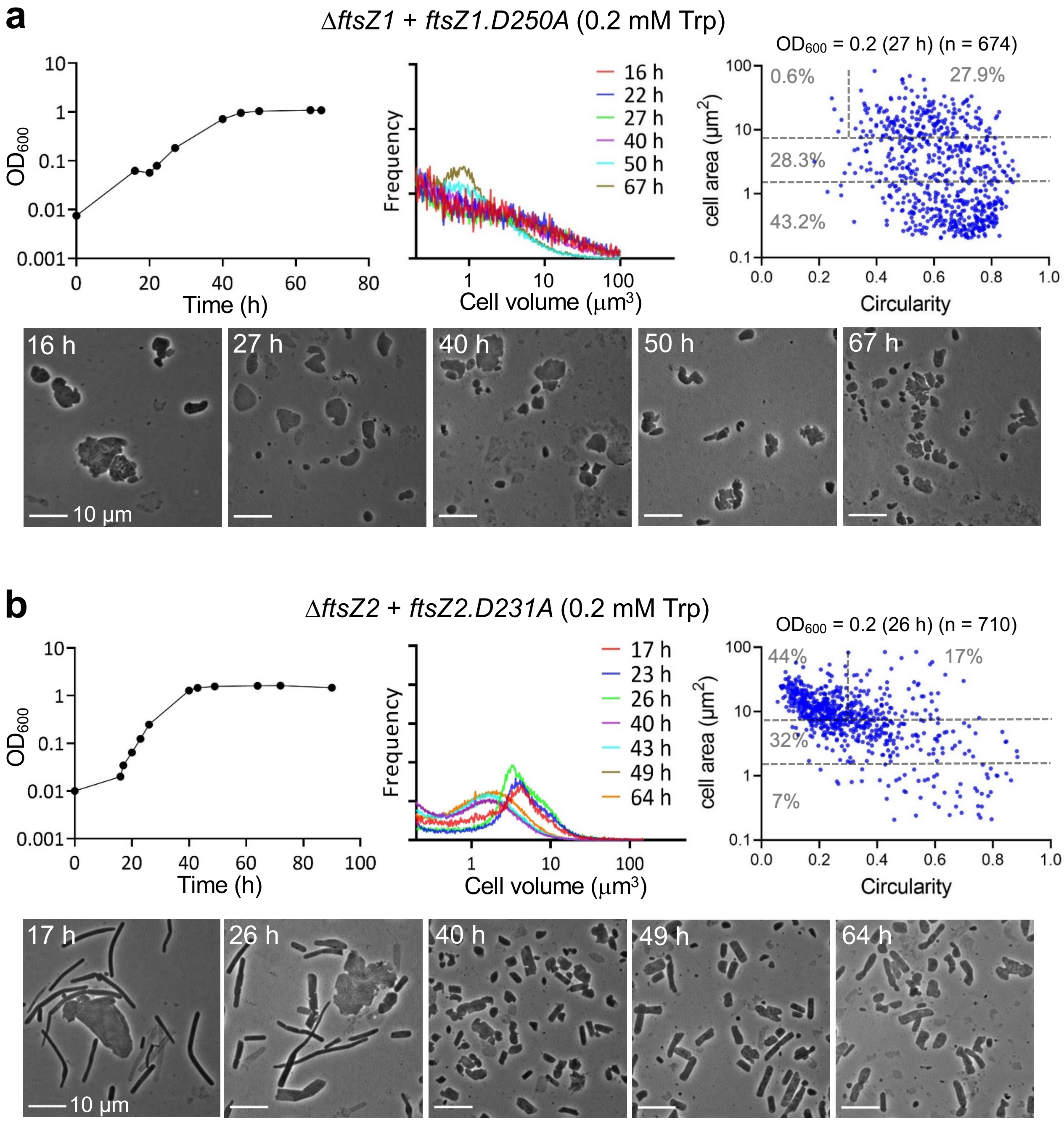

**Figure S13**. **Effects of GTPase active-site (T7) mutants during the growth cycle.**

**(a)** Growth curves, Coulter cell volume distributions and phase-contrast microscopy images of samples from cultures at the indicated timepoints of the Δ*ftsZ1* + pTA962-*ftsZ1.D250A* strain grown with 0.2 mM Trp. **(b)** The same experiment, except with Δ*ftsZ2* + pTA962-*ftsZ2.D231A*. The largest cells early in the culture generally give way to greater levels of debris at later stages.

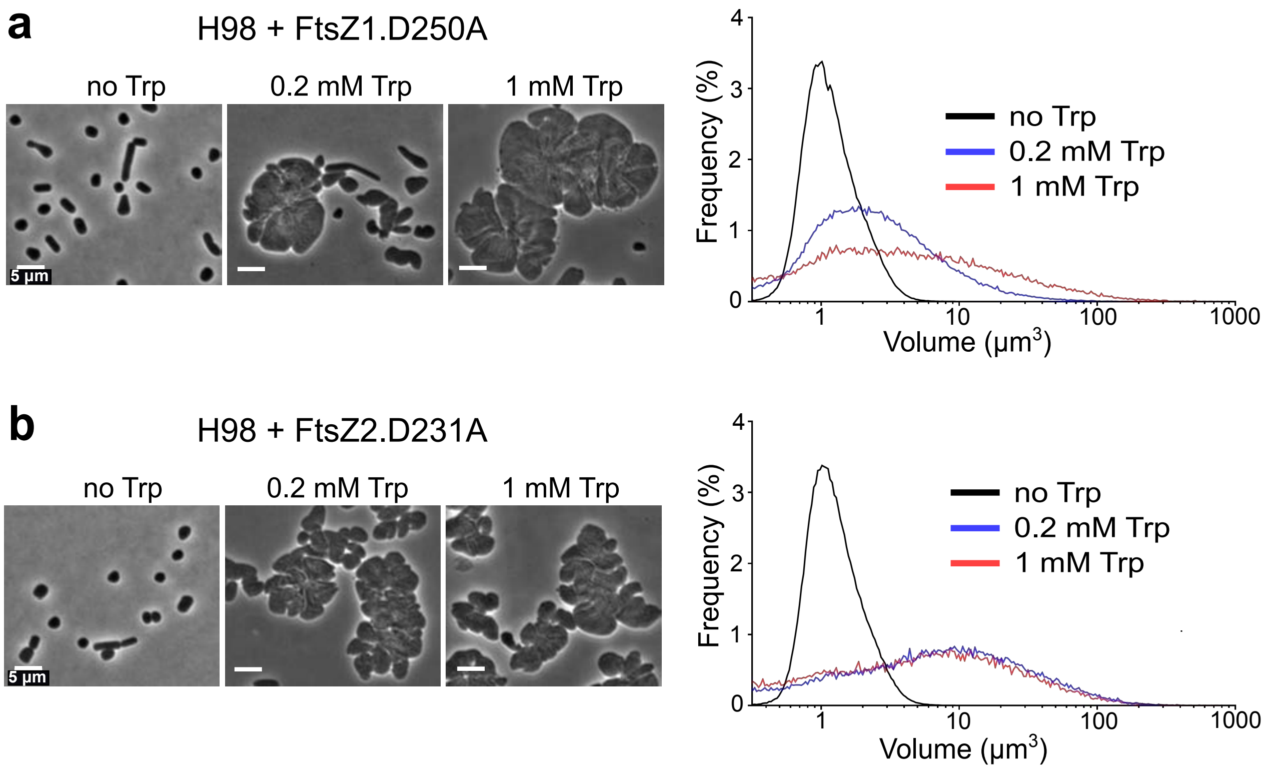

**Figure S14. Dominant-inhibitory phenotypes of T7 mutants *ftsZ1.D250A* and *ftsZ2.D231A*.**

Plasmids based on pTA962 for expression of *ftsZ1.D250A* and *ftsZ.D231A* point mutants were transferred to *H. volcanii* (H98), giving strains ID104 and ID105, respectively (Table S1). **(a)** Phase-contrast microscopy (left) and Coulter cytometry (right) analyses of mid-log cultures expressing *ftsZ1.D250A* with the indicated concentrations of Trp **(b)** Results for the equivalent experiment with *ftsZ2.D231A*. The cell division defects observed with FtsZ1.D250A increase as the inducer concentration increased, whereas a similar strong cell division defect was observed for FtsZ2.D231A at both 0.2 mM and 1 mM Trp.

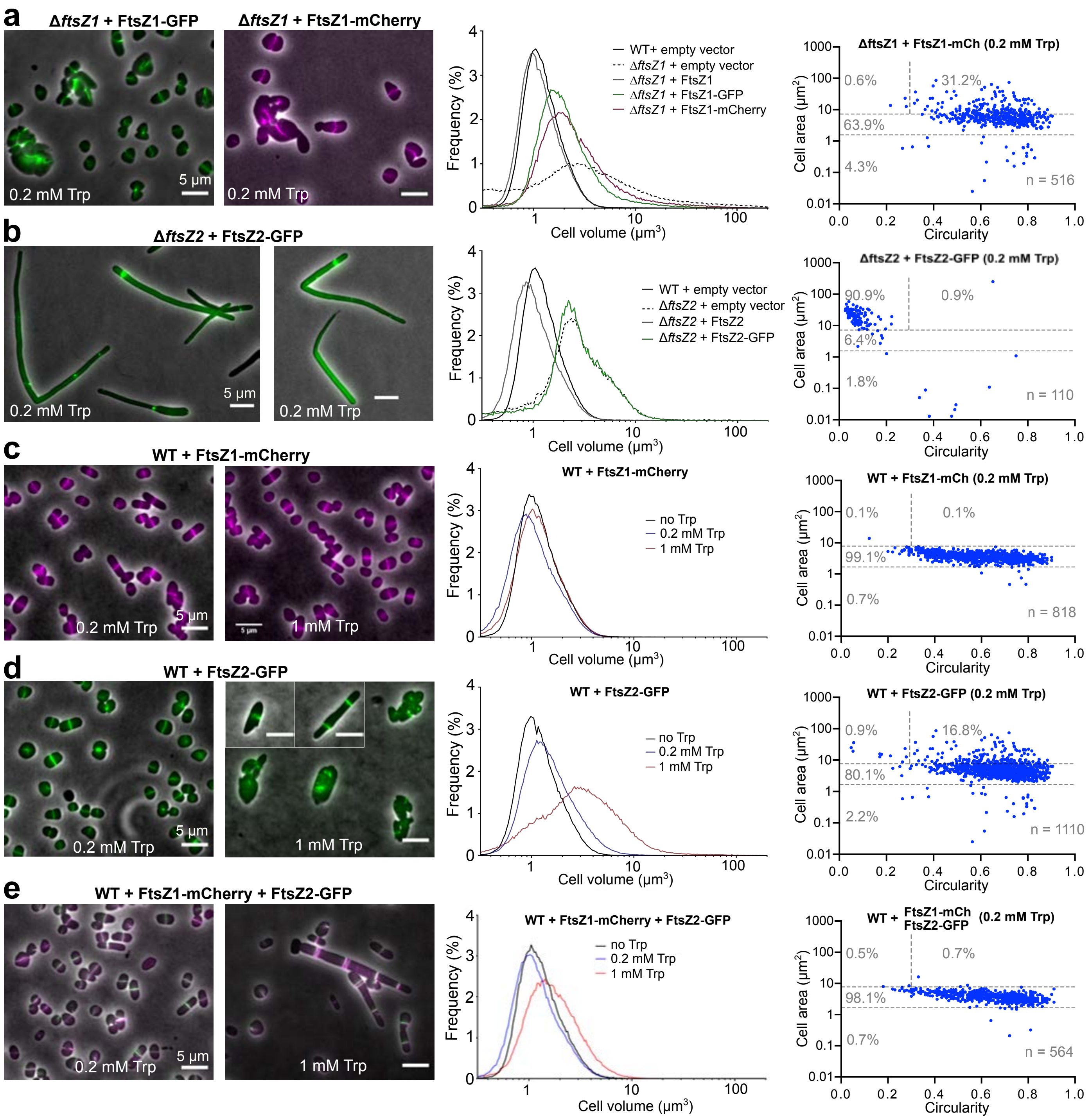

**Figure S15. FtsZ1 and FtsZ2 fluorescent fusions are not fully functional as sole copies but at moderate concentrations have minimal impact on cell division in the wild type.**

**(a-b)** FtsZ1 and FtsZ2 fluorescent fusion proteins were functionally tested in their respective Δ*ftsZ1* or Δ*ftsZ2* backgrounds by phase-contrast and fluorescence microscopy (left) and by Coulter cytometry (right) for cell size. FtsZ1-GFP and FtsZ1-mCherry (0.2 mM Trp) partially complement Δ*ftsZ1*. **(b)**. FtsZ2-GFP was unable to complement the Δ*ftsZ2* background, whereas the untagged protein achieves full complementation (0.2 mM Trp). The same dataset for the wild-type control is shown in both graphs as a reference. **(c-e)** When expressed in wild-type cells, FtsZ1-mCherry or FtsZ2-GFP, or both, cause minimal effects on cell size and shape at a moderate level of expression (0.2 mM Trp), and show sharp midcell bands. These proteins are therefore useful localization markers for division, although detailed analyses of FtsZ subcellular ultrastructure and dynamics await the development of functional complete labelling. Scale bars, 5 µm.

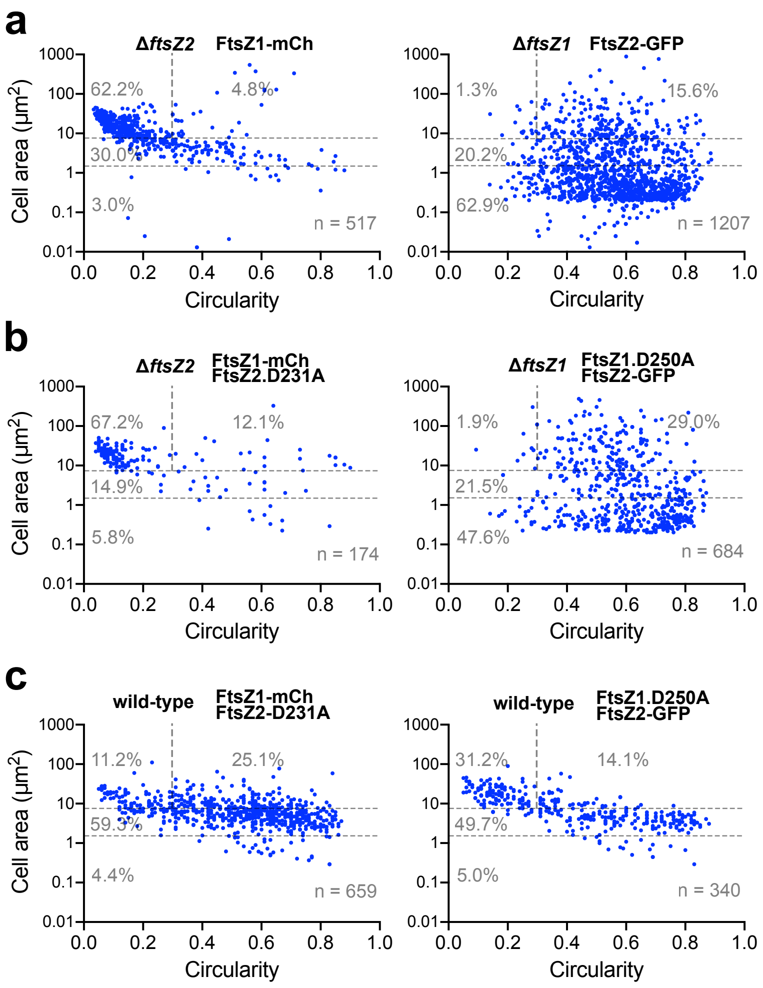

**Figure S16. Cell shape analyses for the FtsZ localization interdependency studies.**

Cell area and shape (circularity) were determined for individual cells, as per Fig. 5 (0.2 mM Trp), and data were combined from two replicate experiments in for each plot. The plots are labelled with the strain's relevant genomic background (left) and the *ftsZ* variant(s) expressed on the plasmid (right).

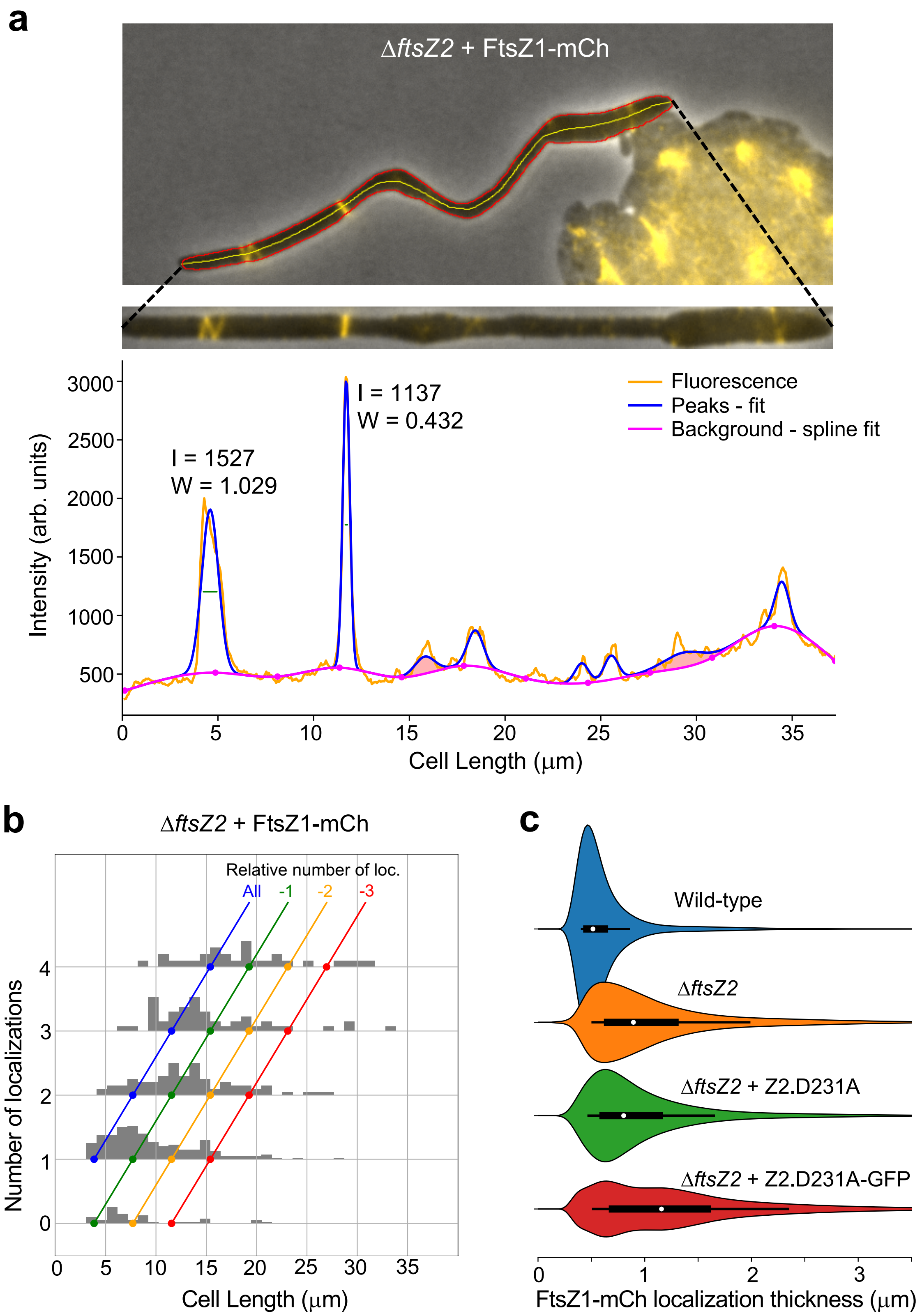

**Figure S17. FtsZ1-mCherry localization in *ftsZ2*-mutant strains.**

**(a)** Demonstration of the automated image analysis procedure for determining FtsZ localization parameters. Cell outlines were obtained (red), and the fluorescence (FtsZ1-mCherry in yellow) was quantified by averaging the intensity on the transverse axis to create a longitudinal intensity profile. Gaussian peaks were fitted to the significant localizations and a spline fit to the background. The localization thickness (W) was taken as the width of the fitted Gaussian peaks at half height (μm), and the intensity (I) was taken as the integrated peak area (per μm across the cell). See *Methods* for further details. **(b)** Violin plots of the thickness of FtsZ1-mCh localization in the indicated strain backgrounds; the median is indicated by a white dot, the thick bar is the interquartile range, and thin bar is the $9^{th}$-$91^{st}$ percentile range. Explanation of the experiment using the Δ*ftsZ2* + FtsZ2.D231A-GFP + FtsZ1-mCh strain is given in the Supplementary results and discussion and Fig. S19.

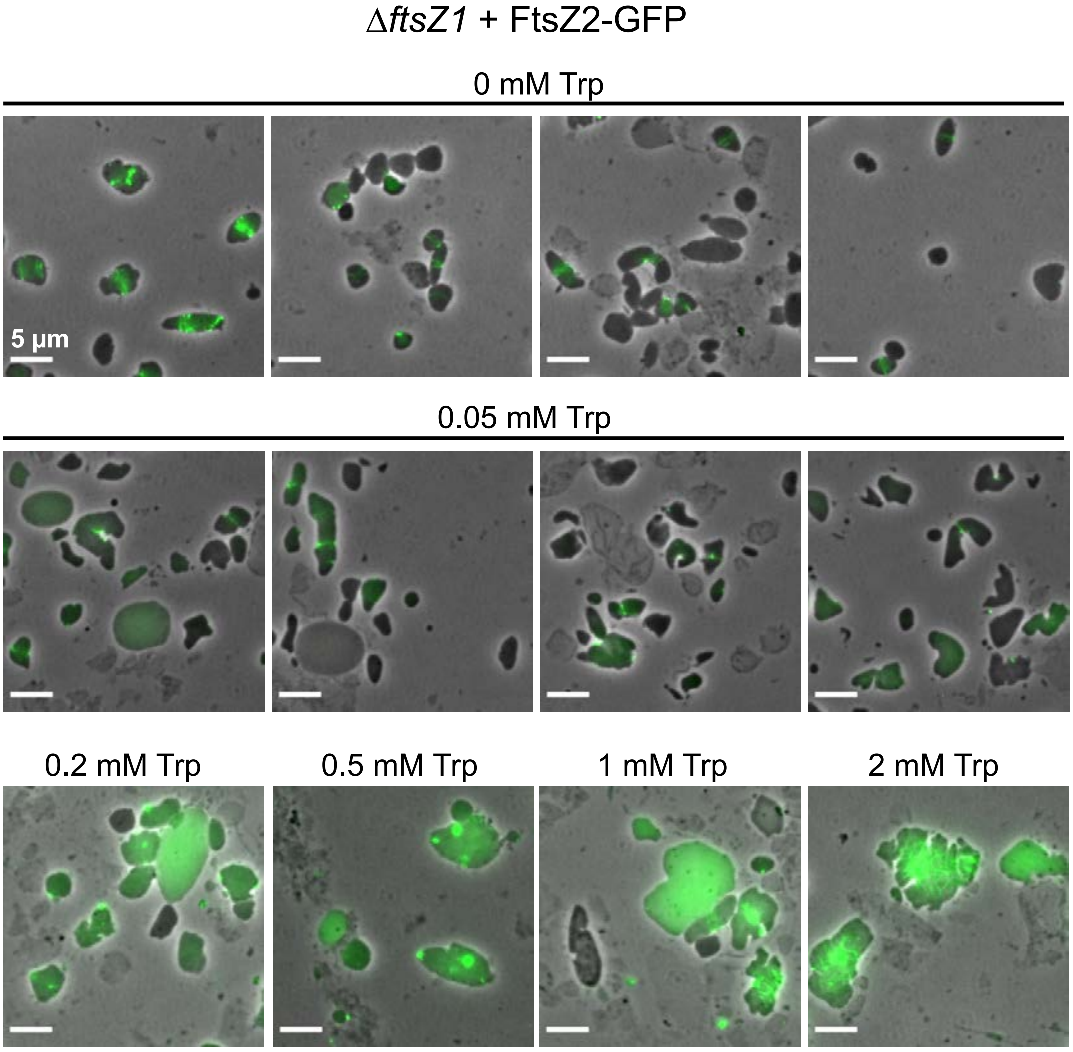

**Figure S18. Localization of FtsZ2-GFP in the absence of *ftsZ1*.**

*H. volcanii* ID90 (Δ*ftsZ1* + *ftsZ2*-GFP) was grown with the indicated concentrations of Trp and sampled during mid-log growth for fluorescence and phase-contrast microscopy. Without Trp induction or at the lowest concentration of Trp tested (0.05 mM), FtsZ2-GFP was produced at a low level in some cells and showed occasionally a diffuse or poorly structured ring. Infrequently, a ring that appeared more normal was observed at apparent division constrictions. At the higher concentrations of Trp, the elevated levels of FtsZ2-GFP were seen diffusely in the cell or formed bright foci or patches at areas including irregular indentations at the cell edges, but no rings. At the highest Trp concentration more intense large masses of localized FtsZ2-GFP appeared generally towards the center of the giant plate cells. Elevated debris and 'ghost' cell envelopes were more frequently seen at the high Trp concentrations, which are associated with a severe division defect. All scale bars are 5 µm.

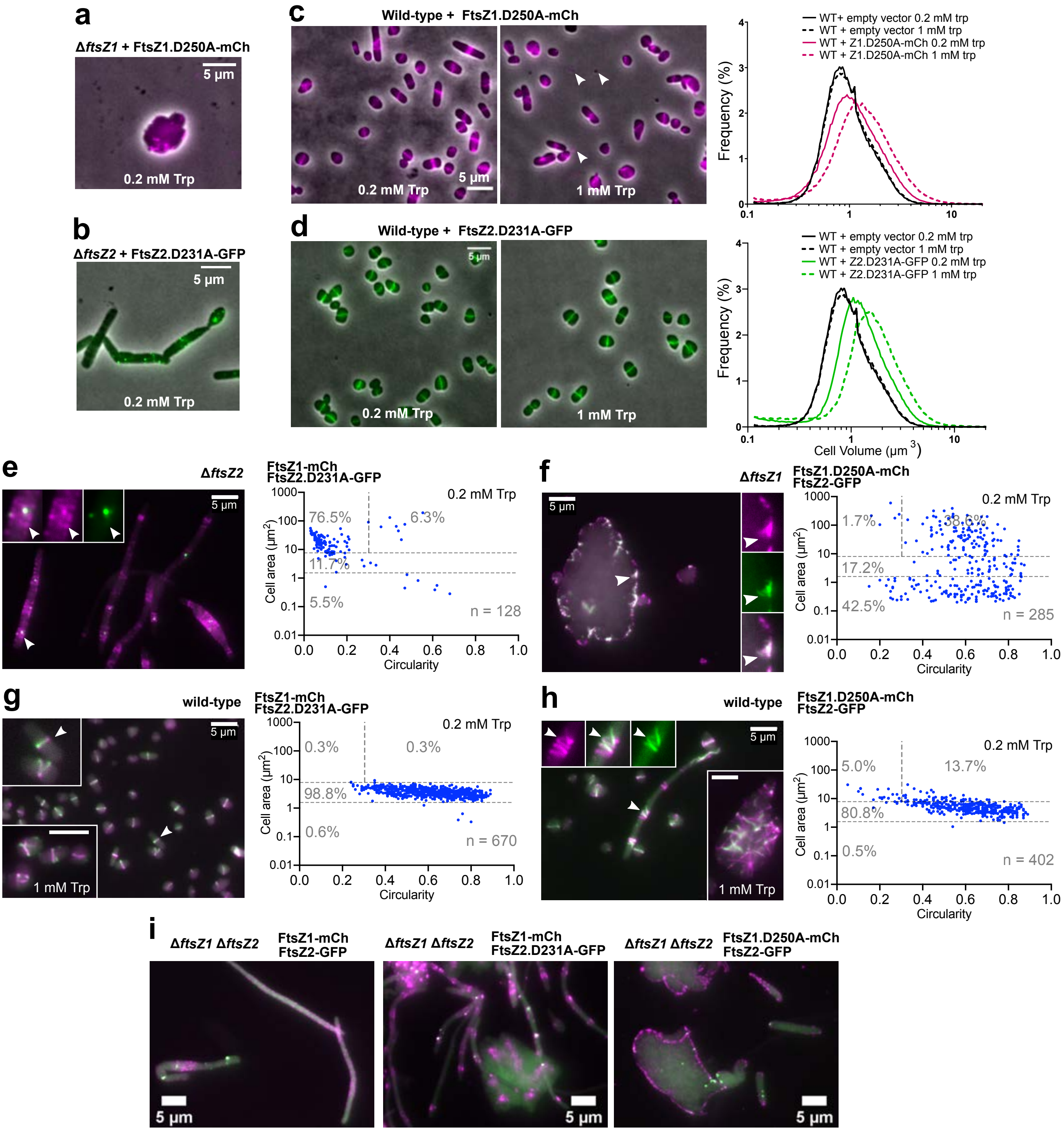

**Figure S19. Localization studies of FtsZ T7-loop mutants, *ftsZ1.D250A-mCh* and *ftsZ2.D231A-GFP*.**

**(a-b)** The FP-tagged T7 mutants fail to complement their respective Δ*ftsZ* strain. **(c-d)** Suppression of the severe dominant-inhibitory effects of the T7-mutants (see Fig. S14) by the fluorescent tags, shown by phase-contrast and fluorescence microscopy (left) and Coulter cytometry (right). FtsZ1.D250A-mCh showed aberrant localization and cellular distortions and envelope protrusions associated with the fluorescent filaments; small fluorescent free particles are also evident (arrowheads). Yet the cell size distribution was only subtly affected. Similar results were obtained with FtsZ1.D250A-GFP (*H. volcanii* ID153) (data not shown). FtsZ2.D231A-GFP shows similar localization to wild-type (FtsZ2-GFP), with only a moderate increase in cell size increase observed at 1 mM Trp; note that FtsZ2-GFP had a much stronger influence (Fig. S15d). **(e-h)** Microscopy and cell size/shape analysis plots for the indicated strains (0.2 mM Trp), containing one tagged wild-type protein and the alternate tagged the T7-loop mutant (colocalization appears white). In panels **(g)** and **(h)**, the 1 mM Trp data is shown in the lower insets; the morphology percentages were: **(g)** 95.4% wild-type-like, 3.2% giant plates, 0.9% filaments, and 0.5% debris (n = 439), and **(h)** 40.7% wild-type-like, 25.5% giant plates, 29.1% filaments, and 5.1% debris (n = 196). **(i)** Co-expression (0.2 mM Trp) of both fusions in Δ*ftsZ1* Δ*ftsZ2* (left) confirms their co-dependence on the wild-type proteins. (Middle and Right) Localization of the two alternate combinations of wild-type and T7-mutant *ftsZ* in the Δ*ftsZ1* Δ*ftsZ2* background (compare to corresponding panels e-h).

### Supplementary Tables

**Table S1**. Strains used in this study.

| Strain | Genotype | Description | Source |
|---|---|---|---|
| | | ***E. coli*** | |
| DH5α | *fhuA2 Δ(argF-lacZ)U169 phoA glnV44 Φ80 Δ(lacZ)M15 gyrA96 recA1 relA1 endA1 thi-1 hsdR17* | General cloning strain for plasmid construction | Invitrogen |
| C2925 | *ara-14 leuB6 fhuA31 lacY1 tsx78 glnV44 galK2 galT22 mcrA dcm-6 hisG4 rfbD1 R(zgb210::Tn10)* $Tet^S$ *endA1 rspL136* ($Str^R$) *dam13::Tn9* ($Cam^R$) *xylA-5 mtl-1 thi-1 mcrB1 hsdR2* | DNA methylation-deficient strain for preparation of demethylated plasmids for *H. volcanii* transformation | New England Biolabs |
| | | ***H. volcanii*** | |
| DS70 | (DS2) ΔpHV2 | Wild-type *H. volcanii* DS2 cured of pHV2 | T. Allers |
| H26 | (DS70) Δ*pyrE2* | Auxotroph (uracil) | T. Allers |
| H98 | (DS70) Δ*pyrE2* Δ*hdrB* | Auxotroph (uracil, hypoxanthine and thymidine) | T. Allers |
| ID41 | (H98) pTA962 | (wild type for *ftsZ*) carrying pTA962 | [13] |
| ID56 | (H98) *p.fdx-hdrB p.tna-ftsZ1* | Trp-regulated expression of genomic *ftsZ1* (HVO_0717) | This study |
| ID57 | (H98) *p.fdx-hdrB p.tna-ftsZ2* | Trp-regulated expression of genomic *ftsZ2* (HVO_0581) | This study |
| ID76 | (H98) *p.fdx-hdrB* Δ*ftsZ1* | Deletion of *ftsZ1* | This study |
| ID77 | (H98) *p.fdx-hdrB* Δ*ftsZ2* | Deletion of *ftsZ2* | This study |
| ID25 | (H98) pTA962-*ftsZ1* | Carries plasmid for expression of *ftsZ1* (*p.tna-ftsZ1*) | This study |
| ID26 | (H98) pTA962-*ftsZ2* | Carries plasmid for expression of *ftsZ2* (*p.tna-ftsZ2*) | This study |
| ID104 | (H98) pTA962-*ftsZ1.D250A* | Carries plasmid for expression of *ftsZ1.D250A* (GTPase T7-loop mutant) | [13] |
| ID105 | (H98) pTA962-*ftsZ2.D231A* | Carries plasmid for expression of *ftsZ2.D231A* (GTPase T7-loop mutant) | This study |
| ID16 | (H98) pIDJL40-*ftsZ1* | Carries plasmid for expression of *ftsZ1-gfp* | [13] |
| ID49 | (H98) pIDJL114 | Carries plasmid for expression of *ftsZ1-mCherry* | This study |
| ID17 | (H98) pIDJL40-*ftsZ2* | Carries plasmid for expression of *ftsZ2-gfp* | This study |
| ID50 | (H98) pIDJL115 | Carries plasmid for expression of *ftsZ2-mCherry* | This study |
| ID153 | (H98) pIDJL40-*ftsZ1.D250A* | Carries plasmid for expression of *ftsZ1.D250A-gfp* | This study |
| ID225 | (H98) pHVID100 | Carries plasmid for expression of *ftsZ1.D250A-mCherry* | This study |
| ID226 | (H98) pHVID101 | Carries plasmid for expression of *ftsZ2.D231A-gfp* | This study |
| ID67 | (H98) pIDJL134 | Carries plasmid for dual expression of *ftsZ2-gfp* and *ftsZ1-mCherry* | This study |
| ID229 | (H98) pHVID103 | Carries plasmid for dual expression of *ftsZ2.D231A* and *ftsZ1-mCherry* | This study |
| ID233 | (H98) pHVID105 | Carries plasmid for dual expression of *ftsZ2-gfp* and *ftsZ1.D250A* | This study |
| ID227 | (H98) pHVID104 | Carries plasmid for dual expression of *ftsZ2.D231A-gfp* and *ftsZ1-mCherry* | This study |
| ID231 | (H98) pHVID106 | Carries plasmid for dual expression of *ftsZ2-gfp* and *ftsZ1.D250A-mCherry* | This study |
| ID133 | (ID76) pTA962 | Δ*ftsZ1* carrying pTA962 | This study |
| ID86 | (ID76) pTA962-*ftsZ1* | Δ*ftsZ1* and plasmid for expression of *ftsZ1* | This study |
| ID87 | (ID76) pTA962-*ftsZ2* | Δ*ftsZ1* and plasmid for expression of *ftsZ2* | This study |
| ID88 | (ID76) pTA962-*ftsZ1.D250A* | Δ*ftsZ1* and plasmid for expression of *ftsZ1.D250A* | This study |
| ID89 | (ID76) pIDJL40-*ftsZ1* | Δ*ftsZ1* and plasmid for expression of *ftsZ1-gfp* | This study |
| ID90 | (ID76) pIDJL40-*ftsZ2* | Δ*ftsZ1* and plasmid for expression of *ftsZ2-gfp* | This study |
| ID256 | (ID76) pIDJL114 | Δ*ftsZ1* and plasmid for expression of *ftsZ1-mCherry* | This study |
| ID254 | (ID76) pIDJL40-*ftsZ1.D250A* | Δ*ftsZ1* and plasmid for expression of *ftsZ1.D250A-gfp* | This study |
| ID255 | (ID76) pHVID100 | Δ*ftsZ1* and plasmid for expression of *ftsZ1.D250A-mCherry* | This study |
| ID277 | (ID76) pHVID105 | Δ*ftsZ1* and plasmid for dual expression of *ftsZ2-gfp* and *ftsZ1.D250A* | This study |
| ID276 | (ID76) pHVID106 | Δ*ftsZ1* and plasmid for dual expression of *ftsZ2-gfp* and *ftsZ1.D250A-mCherry* | This study |
| ID134 | (ID77) pTA962 | Δ*ftsZ2* carrying pTA962 | This study |
| ID91 | (ID77) pTA962-*ftsZ1* | Δ*ftsZ2* and plasmid for expression of *ftsZ1* | This study |
| ID92 | (ID77) pTA962-*ftsZ2* | Δ*ftsZ2* and plasmid for expression of *ftsZ2* | This study |
| ID93 | (ID77) pTA962-*ftsZ2.D231A* | Δ*ftsZ2* and plasmid for expression of *ftsZ2.D231A* | This study |
| ID94 | (ID77) pIDJL40-*ftsZ1* | Δ*ftsZ2* and plasmid for expression of *ftsZ1-gfp* | This study |
| ID95 | (ID77) pIDJL40-*ftsZ2* | Δ*ftsZ2* and plasmid for expression of *ftsZ2-gfp* | This study |
| ID257 | (ID77) pHVID101 | Δ*ftsZ2* and plasmid for expression of *ftsZ2.D231A-gfp* | This study |
| ID436 | (ID77) pIDJL114 | Δ*ftsZ2* and plasmid for expression of *ftsZ1-mCherry* | This study |
| ID279 | (ID77) pHVID103 | Δ*ftsZ2* and plasmid for dual expression of *ftsZ2.D231A* and *ftsZ1-mCherry* | This study |
| ID278 | (ID77) pHVID104 | Δ*ftsZ2* and plasmid for dual expression of *ftsZ2.D231A-gfp* and *ftsZ1-mCherry* | This study |
| ID112 | (ID77) Δ*ftsZ1* | Δ*ftsZ2* and Δ*ftsZ1* (Double deletion) | This study |
| ID463 | (ID112) pTA962 | Δ*ftsZ1* Δ*ftsZ2* carrying pTA962 | This study |
| ID464 | (ID112) pTA962-*ftsZ1* | Δ*ftsZ1* Δ*ftsZ2* and plasmid for expression of *ftsZ1* | This study |
| ID465 | (ID112) pTA962-*ftsZ2* | Δ*ftsZ1* Δ*ftsZ2* and plasmid for expression of *ftsZ2* | This study |
| ID473 | (ID112) pTA962-*ftsZ2* + *ftsZ1* | Δ*ftsZ1* Δ*ftsZ2* and plasmid for dual expression of *ftsZ2* and *ftsZ1* | This study |
| ID429 | (ID112) pIDJL134 | Δ*ftsZ1* Δ*ftsZ2* and plasmid for dual expression of *ftsZ2-gfp* and *ftsZ1-mCherry* | This study |
| ID430 | (ID112) pHVID104 | Δ*ftsZ1* Δ*ftsZ2* and plasmid for dual expression of *ftsZ2.D231A-gfp* and *ftsZ1-mCherry* | This study |
| ID431 | (ID112) pHVID106 | Δ*ftsZ1* Δ*ftsZ2* and plasmid for dual expression of *ftsZ2-gfp* and *ftsZ1.D250A-mCherry* | This study |

**Table S2**. Plasmids and oligonucleotides used in this study.

| Plasmid | Description/function | Oligonucleotides used in construction (5' to 3')* | Source |
|---|---|---|---|
| | | ***Plasmids for gene expression in H. volcanii*** | |
| pTA962 | *p.tna* expression vector for *H. volcanii* | | [16] |
| pIDJL40 | pTA962 with *gfp* (BamHI-NotI) | | [13] |
| pTA962-*ftsZ1* | *p.tna* control of *ftsZ1* | *ftsZ1-f*: CCCCCGGGAATTCATATGGACTCTATCGTCGGCGACGC<br>*ftsZ1-r*: CGCGGATCCCTACTCGACGTAGTCGATGTCTTCGAG | This study |
| pTA962-*ftsZ2* | *p.tna* control of *ftsZ2* | *ftsZ2-f*: CCCCCGGGAATTCATATGCAGGATATCGTTCGCGAGGCG<br>*ftsZ2-r*: CGCGGATCCTTACCGGATGACGTCGAGACCGTTG | This study |
| pTA962-*ftsZ1.D250A* | *p.tna* control of *ftsZ1.D250A* | | [13] |
| pTA962-*ftsZ2.D231A* | *p.tna* control of *ftsZ2.D231A* | *ftsZ2-f* and *ftsZ2-r*, above, and:<br>*Z2.D231A-f*: CAACCTCGACTACGCCGCCATGTCGACCATCATG<br>*Z2.D231A-r*: CATGATGGTCGACATGGCGGCGTAGTCGAGGTTG | This study |
| pIDJL40-*ftsZ1* | *p.tna* control of *ftsZ1-gfp* | | [13] |
| pIDJL40-*ftsZ2* | *p.tna* control of *ftsZ2-gfp* | *ftsZ2-f*, above.<br>*ftsZ2(NS)-r*: CGCGGATCCCCGGATGACGTCGAGACCGTTG | This study |
| pIDJL114 | *p.tna* control of *ftsZ1-mCherry* | *mCh-f*: GGCCGGATCCGCTGGCTCCGCTGCTGGTTC<br>*mCh-r*: GGAAGAATGCGGCCGCTTACTTGTACAGCTCGTCCATGCC | This study |
| pIDJL115 | *p.tna* control of *ftsZ2-mCherry* | As per pIDJL114 | This study |
| pIDJL40-*ftsZ1.D250A* | *p.tna* control of *ftsZ1.D250A-gfp* | *ftsZ1-f*, above.<br>*ftsZ1(NS)-r*: CGCGGATCCCTCGACGTAGTCGATGTCTTCGAG | This study |
| pHVID100 | *p.tna* control of *ftsZ1.D250A-mCherry* | | This study |
| pHVID101 | *p.tna* control of *ftsZ2.D231A-gfp* | *ftsZ2-f* and *ftsZ2(NS)-r*, above. | This study |
| pHVID102 | *p.tna* control of *ftsZ2.D231A-mCherry* | | This study |
| pTA962-*ftsZ2 + ftsZ1* | *p.tna* control of *ftsZ2 and ftsZ1* | | This study |
| pIDJL134 | *p.tna* control of *ftsZ2-gfp* and *ftsZ1-mCherry* | | This study |
| pHVID103 | *p.tna* control of *ftsZ2.D231A* and *ftsZ1-mCherry* | | This study |
| pHVID104 | *p.tna* control of *ftsZ2.D231A-gfp* and *ftsZ1-mCherry* | | This study |
| pHVID105 | *p.tna* control of *ftsZ2-gfp* and *ftsZ1.D250A* | | This study |
| pHVID106 | *p.tna* control of *ftsZ2-gfp* and *ftsZ1.D250A-mCherry* | | This study |
| | | ***Plasmids for genomic modification of H. volcanii*** | |
| pTA131 | Cloning vector with *pyrE2* marker, for *H. volcanii* genetic modification | | [17] |
| pIDJL74 | pTA131 with spliced *ftsZ1*-upstream flank and *p.tna-ftsZ1* cassette | *Z1USflank-f*: CCGGCCAAGCTTCTCGAAGCCGACGTCACGA<br>*Z1USflank-r*:<br>GCAGCACATCCCCCTTTCGCCAGATCTCCCCCTTGCGTCAGACATC<br>*PtnaUS-f*: CTGGCGAAAGGGGGATGTGCTGC<br>*ftsZ1-r*, above. | This study |
| pIDJL75 | pTA131 with spliced *ftsZ2*-upstream flank and *p.tna-ftsZ2* cassette | *Z2USflank-f*: CCGGCCAAGCTTCAGACCATGTTTACTGCCCGAAC<br>*Z2USflank-r*:<br>GCAGCACATCCCCCTTTCGCCAGATCTAGTTACACCTTTGCCCAGCC<br>G<br>*PtnaUS-f*, and *ftsZ2-r*, above. | This study |
| pIDJL96 | pIDJL74, with *p.fdx-hdrB* inserted at the BglII site (upstream of *p.tna*). For replacement of *ftsZ1* promoter. | | This study |
| pIDJL97 | pIDJL75, with *p.fdx-hdrB* inserted at the BglII site (upstream of *p.tna*). For replacement of *ftsZ2* promoter. | | This study |
| pIDJL128 | pIDJL74 with *ftsZ1* downstream flank replacing the *p.tna-ftsZ1*. | *Z1DSflank-f*: CCCCCAGATCTGTCGAGTAGTCGAGCCGTCCC<br>*Z1DSflank-r*: CCCCCGGATCCAGCGTGGGGAATCTCTTCGAG | This study |
| pIDJL129 | pIDJL75 with *ftsZ2* downstream flank replacing the *p.tna-ftsZ2*. | *Z2DSflank-f*: CCCCCAGATCTGTCATCCGGTAACGCCCTGTC<br>*Z2DSflank-r*: CCCCCGGATCCCTCAAGCAGGTCGCAAAGCAT | This study |
| pIDJL142 | pIDJL128 with *p.fdx-hdrB* marker between flanks (BglII). | | This study |
| pIDJL143 | pIDJL129 with *p.fdx-hdrB* marker between flanks (BglII). | | This study |

*Relevant restriction enzyme cut sites for cloning are underlined.

**Table S3.** Number of tubulin superfamily sequences identified in the indicated archaea*.

| Taxon represented | Species and strain | Abbr. | FtsZ1 | FtsZ2 | CetZ | Total |
|---|---|---|---|---|---|---|
| **Euryarchaeota** | | | | | | |
| Archaeoglobi | *Archaeoglobus fulgidus DSM 4304* | ARCFU | 1 | 1 | 1 | 3 |
| | *Ferroglobus placidus DSM 10642* | FERPA | 1 | 1 | 2 | 4 |
| | *Geoglobus acetivorans (taxid: 565033)* | GEOAC | 1 | 1 | 2 | 4 |
| Methanoliparia | *Euryarchaeota archaeon NM1a (taxid:2491083)* | EANM1 | 1 | 1 | 0 | 2 |
| Thermoplasmata | *Thermoplasma acidophilum DSM 1728* | THEAC | 1 (+1)* | 0 | 0 | 2 |
| | *Methanomassiliicoccus luminyensis B10* | METLB | 3 | 1 | 0 | 6 |
| Hadesarchaea | *Hadesarchaea archaeon DG-33* | HADES | 1 | 0 | 0 | 1 |
| Methanobacteria | *Methanobrevibacter ruminantium M1*** | METRM | 1 | 0 | 0 | 1 |
| | *Methanobacterium lacus (taxid:877455)*** | METLA | 1 | 0 | 0 | 1 |
| | *Methanosphaera stadtmanae DSM 3091*** | METST | 1 | 0 | 0 | 1 |
| | *Methanothermobacter thermautotrophicus str. Delta H*** | METTH | 1 | 0 | 0 | 1 |
| | *Methanothermus fervidus DSM 2088*** | METFV | 1 | 0 | 0 | 1 |
| Methanococci | *Methanocaldococcus jannaschii DSM 2661* | METJA | 1 | 1 | 0 | 2 |
| | *Methanococcus maripaludis S2* | METMP | 1 | 1 | 0 | 2 |
| Methanonatronarchaeia | *Methanonatronarchaeum thermophilum* | MENAT | 1 | 1 | 0 | 2 |
| Methanopyri | *Methanopyrus kandleri AV19*** | METKA | 1 | 0 | 0 | 1 |
| Nanohaloarchaeota | *Candidatus Nanosalina sp. J07AB43* | NANS0 | 1 | 1 | 0 | 2 |
| | *Candidatus Haloredivivus sp. G17* | HALSG | 0 | 1 | 0 | 1 |
| Halobacteria | *Haloarcula japonica DSM 6131* | HALJP | 1 | 1 | 3 | 6 |
| | *Natronomonas pharaonis DSM 2160* | NATPD | 1 | 1 | 2 | 5 |
| | *Halobacterium salinarum R1* | HALS3 | 1 | 1 | 3 | 8 |
| | *Halococcus saccharolyticus DSM 5350* | HALSC | 1 | 1 | 0 | 2 |
| | *Natronoarchaeum philippinense* | NATPH | 1 | 1 | 2 | 4 |
| | *Haloferax volcanii DS2* | HALVD | 1 | 1 | 6 | 8 |
| | *Haloquadratum walsbyi DSM16790* | HALWD | 1 | 1 | 1 | 3 |
| | *Halorubrum lacusprofundi ACAM 34* | HALLT | 1 | 1 | 3 | 5 |
| | *Natrialba magadii ATCC 43099* | NATMM | 1 | 1 | 4 | 6 |
| Methanomicrobia | *Methanocella arvoryzae MRE50* | METAR | 1 | 1 | 0 | 3 |
| | *Methanoculleus marisnigri JR1* | METMJ | 1 | 1 | 2 | 4 |
| | *Methanophagales archaeon (taxid: 2056316)* | METPH | 1 | 1 | 0 | 3 |
| | *Methanosarcina acetivorans C2A* | METAC | 1 | 1 | 1 | 3 |
| Theionarchaea | *Theionarchaea archaeon DG-70* | THEIO | 1 | 1 | 0 | 2 |
| Thermococci | *Pyrococcus furiosus DSM 3638* | PYRFU | 1 | 1 | 1 | 3 |
| | *Thermococcus kodakarensis KOD1* | THEKO | 1 | 1 | 1 | 3 |
| | *Palaeococcus pacificus DY20341* | PALPA | 1 | 1 | 1 | 3 |
| ***Asgard GROUP:*** | | | | | | |
| Heimdallarchaeota | *Candidatus Heimdallarchaeota archaeon* | HEIMD | 1 | 0 | 0 | 2 |
| Lokiarchaeota | *Lokiarchaeum sp. GC14_75* | LOKSG | 0 | 1 | 0 | 1 |
| Odinarchaeota | *Candidatus Odinarchaeota archaeon LCB_4* | ODINA | 1 | 1 | 0 | 3 |
| Thorarchaeota | *Candidatus Thorarchaeota archaeon AB_25* | THOAR | 1 | 1 | 0 | 3 |
| ***DPANN GROUP:*** | | | | | | |
| Aenigmarchaeota | *Candidatus Aenigmarchaeota archaeon CG1_02_38_14* | AENIG | 1 | 1 | 0 | 2 |
| Diapherotrites | *Candidatus Diapherotrites archaeon* | DIAPH | 2 | 0 | 0 | 2 |
| Micrarchaeota | *Candidatus Micrarchaeum acidiphilum ARMAN-2* | MICA2 | 2 | 2 | 0 | 4 |
| Pacearchaeota | *Candidatus Pacearchaeota archaeon CG1_02_30_18* | PACEA | 1 | 1 | 0 | 2 |
| Parvarchaeota | *Candidatus Parvarchaeum acidiphilum ARMAN-4* | PARA4 | 1 | 1 | 0 | 2 |
| Woesearchaeota | *Candidatus Woesearchaeota archaeon CG1_02_33_12* | WOESE | 1 | 1 | 0 | 2 |
| Nanoarchaeota | *Nanoarchaeum equitans Kin4-M* | NANEQ | (2) | 0 | 0 | 2 |
| ***TACK GROUP:*** | | | | | | |
| Bathyarchaeota | *Bathyarchaeota archaeon B23* | BATHA | 0 | 1 | 0 | 2 |
| Geothermarchaeota | *Candidatus Geothermarchaeota archaeon ex4572_27* | GEOAR | 0 | 1 | 0 | 2 |
| Korarchaeota | *Candidatus Korarchaeum cryptofilum OPF8* | KORCO | (1) | 0 | 0 | 7 |
| Marsarchaeota | *Candidatus Marsarchaeota G1 archaeon BE_D* | MARSA | 0 | 0 | 0 | 0 |
| Verstraetearcheota | *Candidatus Methanosuratus sp. (taxid: 2495426)* | VERST | 0 | 0 | 0 | 0 |
| **Crenarchaeota** | | | | | | |
| Acidilobales | *Acidilobus saccharovorans 345-15* | ACIS3 | 0 | 0 | 0 | 1 |
| Desulfurococcales | *Aeropyrum pernix K1* | AERPE | 0 | 0 | 0 | 0 |
| Fervidicoccales | *Fervidicoccus fontis Kam940* | FERFK | 0 | 0 | 0 | 0 |

| Taxon represented | Species and strain | Abbr. | FtsZ1 | FtsZ2 | CetZ | Total |
|---|---|---|---|---|---|---|
| Sulfolobales | *Sulfolobus acidocaldarius DSM 639* | SULAC | 0 | 0 | 0 | 0 |
| Thermoproteales | *Pyrobaculum aerophilum str. IM2* | PYRAE | 0 | 0 | 0 | 0 |
| **Thaumarchaeota** | | | | | | |
| Cenarchaeales | *Cenarchaeum symbiosum A* | CENSY | 0 | 0 | 0 | 1 |
| Nitrosopumilales | *Nitrosopumilus maritimus SCM1* | NITMS | 0 | 0 | 0 | 1 |
| | *Nitrosopumilales archaeon CG_4_10_14_0_8_um_filter_34_8* | NITAR | 0 | 0 | 0 | 1 |
| Nitrososphaeria | *Candidatus Nitrososphaera evergladensis SR1* | NITES | 0 | 0 | 0 | 1 |

*Grey text indicates taxa in which genome data might be incomplete (*i.e.,* in contig. or scaffold form), or the taxon or species is currently *Candidatus* status. Numbers in parentheses in the FtsZ1 column indicate deeply branching proteins that appear to be FtsZ family members (rather than CetZ or non-canonical), that nevertheless have uncertain designation as specifically as archaeal FtsZ1 or FtsZ2 or bacterial/plant FtsZ. Some species contain other non-canonical sequences, and these are included in the total.
**Species for which a complete genome is available that encodes one or more pseudomurein binding domains (pfam09373 in Methanobacteria) and/or have been shown to have a pseudomurein wall (Methanobacteria and *Methanopyrus kandleri* AV19).

**Table S4.** Average percent identity amongst domains of the archaeal FtsZ1, FtsZ2 and CetZ protein families*.

| | **N-tail** | | | **N-term. (GTP binding)** | | | **C-term. (polymerization)** | | | **C-tail** | | | **Overall** | | |
|---|---|---|---|---|---|---|---|---|---|---|---|---|---|---|---|
| | FtsZ1 | FtsZ2 | CetZ | FtsZ1 | FtsZ2 | CetZ | FtsZ1 | FtsZ2 | CetZ | FtsZ1 | FtsZ2 | CetZ | FtsZ1 | FtsZ2 | CetZ |
| FtsZ1 | 24 | 18 | N/A | 64 | 51 | 26 | 55 | 43 | 21 | 23 | 18 | 11 | 54 | 44 | 23 |
| FtsZ2 | 18 | 20 | N/A | 51 | 60 | 25 | 43 | 52 | 20 | 18 | 18 | 11 | 44 | 51 | 22 |
| CetZ | N/A | N/A | N/A | 26 | 25 | 43 | 21 | 20 | 40 | 11 | 11 | 25 | 23 | 22 | 44 |

*Data were averaged from ClustalX percent-identity matrices (excluding self-alignments), derived from separate multiple alignments of the four domains and the whole protein (overall), including all proteins from these three families identified in the 60 archaea listed in Table S3. CetZ proteins do not have the N-terminal tail (N/A).

**Table S5.** Identified non-synonymous and intergenic small variants in genomes of *H. volcanii* ID76 (Δ*ftsZ1*), ID77 (Δ*ftsZ2*), and ID112 (Δ*ftsZ1* Δ*ftsZ2*), compared to the draft reference genome of H98*.

| **Locus** | **Annotation/region** | **Variant position (in DS2)** | **DS2/H98 reference** | **ID76 (Δ*ftsZ1*)** | **ID77 (Δ*ftsZ2*)** | **ID112 (Δ*ftsZ1* Δ*ftsZ2*)** | **Amino acid change** |
|---|---|---|---|---|---|---|---|
| HVO_1307 | Hypothetical protein | 1192773 | ACCGAC CCCGCC GCCTCC GCTCTC GACTCC GACCCC GCCGCC | ACCGAC CCCGCC GCCTCC GCTCTC GACTCC GACCCC GCCGCC / **ACCGA CCCCG CCGCC** | ACCGAC CCCGCC GCCTCC GCTCTC GACTCC GACCCC GCCGCC | ACCGAC CCCGCC GCCTCC GCTCTC GACTCC GACCCC GCCGCC | Deletion of **SALDSDP AA** (12-20) |
| HVO_1279 *(hdrA)* | Dihydrofolate reductase | 1166715 | G | G / **C** | G | G / **C** | E11**D** |
| HVO_2948 *(pheS)* | Phenylalanyl-tRNA ligase α-chain | 2783019 | C | C | **T** | **T** | G38**D** |
| HVO_0809 *(metS)* | Methionyl-tRNA ligase (MetG family) | 728964 | C | C | **G** | **G** | G167**A** |
| HVO_0424 | ABC transporter ATP-binding protein (ABCE1 family). | 380991 | A | A | A | **G** | D143**G** |
| HVO_1711 *(sgaI)* | Glucoamylase | 1578118 | A | A | A | **G** | D268**G** |
| HVO_1018 *(recJ3)* | RecJ-like exonuclease | 927660 | TCGGCG GCGGCG TCTCCG GCGGCG GC | TCGGCG GCGGCG TCTCCG GCGGCG GC | TCGGCG GCGGCG TCTCCG GCGGCG GC | TCGGCG GCGGCG TCTCCG GCGGCG GC / **TCGGC GGCGG CGTCTC CGGCG GCGGC GTCTCC GGCGG CGGC** | Insertion of **VSGGG** after G595 |
| Intergenic region (1595913..1 596083) | Region of unknown function between and downstream of genes HVO_1725 (*orc5* - Orc1/Cdc6-type DNA replication protein) and HVO_1726 (hypothetical protein). [Note: the *oriC2* origin is upstream of *orc5*.] | 1595982 | A | A | A | **G** | |

*The nucleotide and corresponding amino acid changes are shown in bold text when they differ from the H98 reference sequence. The forward-slash separates two sequences at a heterozygous site. The sequences shown in the *DS2/H98 reference* column are the same as the *H. volcanii* DS2 complete genome (NC_013967), and their positions in the DS2 sequence are indicated in the *Variant position* column. Additional differences between DS2 and H98 are not shown; genome sequence data have been deposited at NCBI under BioProject PRJNA681931.

#### Supplementary Video Legends

**Video S1. Time-lapse microscopy of FtsZ1 and FtsZ2 depletion and restoration.** FtsZ depletion was achieved by firstly growing *H. volcanii* ID56 (*p.tna-ftsZ1*) and ID57 (*p.tna-ftsZ2*) in media with Trp (2 mM Trp). The cells were then washed in fresh media without Trp, and samples were placed on a soft agarose gel media pad, without Trp, using the submerged-sandwich technique for time-lapse imaging (left two panels). For FtsZ restoration (right two panels), the *p.tna-ftsZ* strains were initially grown without Trp in batch cultures, and then restoration was initiated by adding Trp to 0.2 mM for *ftsZ1* induction, and 2 mM Trp for *ftsZ2* induction, to the agarose pad. Depletion caused cells to grow without dividing, and some cells of both strains displayed occasional budding-like events instead. Restoration of division occurred in both strains, with the giant cells dividing at multiple locations, quite asynchronously and occasionally asymmetrically. Cell growth rate declines in the latter part of the videos, possibly due to local depletion of nutrients.

**Video S2. Time-lapse microscopy of division/fragmentation of FtsZ1-depleted cells.** *H. volcanii* ID56 (*p.tna-ftsZ1*) that had been previously depleted of FtsZ1 by continuous mid-log culturing in the absence of Trp were time-lapse imaged, and examples of dividing/fragmenting cells were identified. These cells show some division events and unusual ways of generating of cell fragments.

**Video S3. Time-lapse microscopy of fragmentation of FtsZ2-depleted cells.** *H. volcanii* ID57 (*p.tna-ftsZ2*) that had been previously depleted of FtsZ2 by continuous mid-log culturing in the absence of Trp were time-lapse imaged, and examples of fragmenting cells were identified. These cells show some blebbing-like events and unusual ways of generating of cell fragments.

**Video S4. 3D imaging of *H. volcanii* wild-type and Δ*ftsZ1* Δ*ftsZ2* strains.** Confocal laser-scanning microscopy of wild-type and Δ*ftsZ1* Δ*ftsZ2* double-mutant live cells suspended in soft-agarose gel. Wild-type plate (top left) and rod (bottom left) cells showing their flattened morphology. Giant plates lose much of their flattened morphology in liquid culture (top-right), whereas giant rod-like cells appear to maintain the flatness—albeit with some apparently flexibility (in this case a somewhat twisted shape). In this field, a cytoplasmic bridge appears between the pole of a large cell and a smaller adjacent cell. This may be an intermediate state during the late stage of separation.

**Video S5. Time-lapse microscopy of Δ*ftsZ1* – expansion of giant plates on agarose.** *H. volcanii* ID76 mid-log cultures were time-lapse imaged. Some cells show a budding-like process. Note that while the cells showed no clear evidence of division while supported on these soft-gel pads, the confinement at the gel-glass interface can somewhat obscure the detection of individual abutting cells. However, it is useful to compare the results obtained with efficiently dividing cells (Video S1, right panels). In some giant cells, the phase-contrast revealed a dynamic cell structure and possible buckling of the cell surface.

**Video S6. Time-lapse microscopy of Δ*ftsZ1* – occasional division/fragmentation**. *H. volcanii* ID76 mid-log cultures were time-lapse imaged using the submerged sandwich technique, and cells undergoing occasional division or highly acentral division/fragmentation events were identified.

**Video S7. Time-lapse microscopy of Δ*ftsZ2* – expansion of giant cells.** *H. volcanii* ID77 mid-log cultures were time-lapse imaged. Most cells of this strain show the giant plate morphotype, expanding to form extremely large irregular cells when supported on the gel surface. Fragmentation of one cell may be seen in the upper-central panel.

**Video S8. Time-lapse microscopy of Δ*ftsZ1* Δ*ftsZ2* – expansion of giant plates and polar tubulation of filaments.** *H. volcanii* ID112 mid-log cultures were time-lapse imaged. Tubulation-and-fission or budding-like processes were frequently observed at the poles of the filamentous cells.

**Video S9. Time-lapse microscopy of FtsZ1-GFP during multiple rounds of division.** *H. volcanii* ID16 mid-log cultures (with 0.2 mM Trp) were time-lapse imaged over several generations (30 min frame intervals), revealing almost continuous FtsZ1-GFP localization at midcell during multiple cycles of division. In this selected field, a rare cell that fails division may also be seen, which exhibits a complex, dynamic network of aster-like central cluster of filaments containing FtsZ1-GFP. This pattern was common in other giant plates observed during this study.

**Video S10. Time-lapse microscopy of FtsZ1-GFP shows dynamic behavior in midcell rings.** *H. volcanii* ID16 mid-log cultures (with 0.2 mM Trp) were time-lapse imaged (10 min frame intervals). FtsZ1-GFP shows uneven localization around the ring and the fluorescence intensity changes over time, consistent with ongoing polymer assembly and disassembly in the ring. Cells are also clearly seen dividing unilaterally and bilaterally in this field.

**Video S11. FtsZ1 and FtsZ2 dynamically co-localize at midcell during division.** *H. volcanii* ID67 was sampled from mid-log cultures (grown with 0.2 mM Trp) were imaged by time-lapse microscopy of FtsZ1-mCherry (red) and FtsZ2-GFP (green). The proteins both show dynamic movement in the ring and generally co-localize. They show somewhat differing localization intensity within the ring, but both FtsZ rings close down along with the visible constriction of the envelope.
